# A survey of bacterial and fungal communities of table olives

**DOI:** 10.64898/2025.12.16.694624

**Authors:** Eugenio Parente, Rocchina Pietrafesa, Francesca De Filippis, Alessandra De Vivo, Maria Grazia Labella, Marina Hidalgo, Emanuela Lavanga, Annamaria Ricciardi

## Abstract

Table olives are produced from a large number of olive varieties subjected to different trade preparations, resulting in a highly heterogeneous family of fermented foods. To characterise the diversity of bacterial and fungal communities and its relationship with variety, ripeness, and trade preparation, we surveyed 363 samples from 40 producers across 6 countries, combining physicochemical measurements, viable counts, and amplicon-based metagenomics. This is the largest survey of table olive microbial communities to date and includes the first culture-independent characterisation of microbial communities for several Italian PDO and non-PDO varieties, most notably Oliva di Gaeta. The contrast between alkali-treated and naturally fermented olives was the dominant structuring factor, with HALAB (Halophilic and Alkalophilic Lactic Acid Bacteria) and other halophiles enriched in alkali-treated varieties and a diverse array of *Lactobacillaceae* and *Pseudomonadota* characterising naturally fermented olives. Despite these consistent signals, striking variability was observed within the same variety and even within the same producer, driven by stochastic colonization events, house microbiota, and the widespread use of small fermentation vessels. This variability obscured variety-specific microbial signatures and prevented reliable discrimination of Italian PDO varieties from similar non-PDO counterparts using amplicon-based approaches. The ecological and taxonomic complexity documented here, encompassing bacterial and fungal genera with largely untapped starter and flavour potential, provides the foundation for the development of variety-specific microbiome-based starter cultures.

**Highlights:**

- We report a metataxonomic survey of microbial communities in 363 table olive samples
- Alkali treatment and natural fermentation drive distinct microbial community structures
- House microbiota and stochastic colonization generate high within-variety variability
- Microbiome data provide ecological foundations for olive microbiome-based starters

## 1. Introduction

Fermented foods have been part of the human diet for millennia and their diversity is huge (Gänzle, 2022; Hernández-Velázquez et al., 2024; Tamang et al., 2020). Table olives are among the most ancient fermented vegetables from the Mediterranean basin (Langgut and Garfinkel, 2022) and this has resulted in the evolution of a great variety of products (Anagnostopoulos and Tsaltas, 2022; Perpetuini et al., 2020). Fermentation of table olives has multiple functions: removing the bitter glucoside oleuropein, enhancing safety and shelf life through acid production and inhibition of spoilage microorganisms, and improving sensory properties. Five main trade preparations of table olives are recognized by the International Olive Oil Council (https://www.internationaloliveoil.org/wp-content/uploads/2019/11/COI-OT-NC1-2004-Eng.pdf). Treated olives, also known as Spanish style olives, are treated with sodium hydroxide to remove oleuropein before being washed and fermented in brine. Green specialties, like Picholine style olives, on the other hand, are stored at low temperature in brine after the alkali treatment, thus limiting fermentation. Natural olives (Greek style) include both green and ripe olives directly fermented in brines. Dried/shrivelled olives and olives darkened by oxidation are preserved by methods other than fermentation.

Because of their long history, table olives production is strongly associated with local varieties and practices. In Italy four varieties (Oliva di Gaeta, Nocellara del Belice, Bella di Daunia, Oliva Ascolana del Piceno) are included in the Protected Denomination of Origin (PDO) food list and Oliva Taggiasca Ligure has recently received Protected Geographical Indication status (https://ec.europa.eu/agriculture/eambrosia/geographical-indications-register/). Many more are included in the Italian List of Traditional Agricultural Products (https://www.gazzettaufficiale.it/eli/gu/2024/03/13/61/sg/pdf). Spain, Greece, France, Portugal and Turkey also produce a wide variety of table olives, including PDO products. Sensory quality, safety and stability of table olives are affected by several factors, including olive ripeness, trade preparation, and by the growth of microorganisms, either present as natural contaminants on the fruits, salt, brines, and contact surfaces or added as starter cultures (Anagnostopoulos and Tsaltas, 2022; Campus et al., 2018). Geographic origin is also believed to be a contributing factor for initial drupe contamination (Argyri et al., 2020; Ferrocino et al., 2025; Kamilari et al., 2023). Current trends in table olive production aim at controlling the microbiota involved in fermentation and spoilage by sanitizing the raw materials, by adding starter and functional cultures, and/or by reducing salt content (Campus et al., 2018). Addition of starters has been successfully tested for several varieties and, typically, involves the inoculation of lactic acid bacteria (*Lactiplantibacillus pentosus* or *Lactipl. plantarum, Lacticaseibacillus paracasei*: Alfonzo et al., 2023b; Comunian et al., 2017; Lanza et al., 2020, 2021; Paba et al., 2020; Pino et al., 2018, 2019; Randazzo et al., 2017; Rodríguez-Gómez et al., 2023; Ruiz-Barba and Jimenéz-Diaz, 2012; Servili et al., 2006; Tzamourani et al., 2024; Vougiouklaki et al., 2024; Zago et al., 2013), yeasts (several species belonging to the genera *Candida, Wyckerhamomyces, Pichia, Cyteromyces* have been tested: Ciafardini and Zullo, 2019; Montaño et al., 2021; Ruiz-Barba et al., 2024; Tufariello et al., 2019) or their combinations (Benítez-Cabello et al., 2020; Garrido-Fernández et al., 2021). Back-slopping practices (Martorana et al., 2015) have also been described. The composition and dynamics of microbial communities in table olives have been extensively reviewed (Anagnostopoulos and Tsaltas, 2022; Campus et al., 2018; Portilha-Cunha et al., 2020; Tsoungos et al., 2023; Vaccalluzzo et al., 2020). The microbiota of table olives is complex and variable, usually due to the lack of control of unit operations affecting dispersal and selection, and includes microbial groups which are considered beneficial, like lactic acid bacteria, yeasts, and other halophilic microorganisms (Halophilic and Alkalophilic Lactic Acid Bacteria, HALAB, and halophilic *Pseudomonadota*), but also microorganisms associated with spoilage, like some halophiles (*Alkalibacterium, Marinilactobacillus, Celerinatantimonas, some Vibrio*), propionibacteria, clostridia and yeasts (Arroyo-López et al., 2012, 2021; Ballester et al., 2021; de Castro et al., 2018; Penland et al., 2021). We have recently reviewed metataxonomic studies on table olives and found that a core microbiota of approximately 30 genera can be identified, with wide variations in the composition among trade preparations and varieties (Ricciardi et al., 2025).

With a few exceptions (Penland et al., 2020, 2021, 2022) studies on table olives microbiota are relatively small (<50 samples) and often focus on a limited number of varieties (see reviews by Ricciardi et al., 2025; Tsoungos et al., 2023 for details). This, and the variety of sampling plans and experimental approaches used, complicates the interpretation and the integration of these data to obtain a comprehensive view of the factors affecting table olive microbial community assembly and dynamics. The effect of trade preparation on the structure and dynamics of bacterial and/or fungal communities has been evaluated for a few varieties (Cocolin et al., 2013; Maoloni et al., 2022; Ruiz-Barba et al., 2023a, 2023b; Zinno et al., 2017), and there is evidence that identical varieties produced in different geographic locations (Kamilari et al., 2023; Kazou et al., 2020) or even within the same geographical region (Lucena-Padrós and Ruiz-Barba, 2019), or in different years by the same producer (Penland et al. 2020) have measurable differences in microbial community composition, although subsets of core taxa can be identified. This information, together with source tracking studies may assist in understanding the relative importance of dispersal and selection in the assembly of table olive microbial communities and provide new avenues for the selection of microbiome – based starters (Nikoloudaki et al., 2024).

Even with all the complexity and variability in table olive microbiomes, a recent study (López-García et al., 2025), leveraging on a large combined dataset and on machine learning approaches, has been successful in discriminating table olives bacterial profiles based on olive processing type, cultivar, country of origin, and isolation matrix. This opens the possibility that the composition of microbiota can be used in table olive authenticity studies and help discriminating PDO varieties from their non-PDO counterparts.

This motivated us to carry out a large survey of table olives belonging to a wide array of varieties and trade preparations from 6 countries (Italy, Greece, Spain, Cyprus, Peru, Egypt) with a focus on Italian PDO and non-PDO varieties, and including both sound and spoiled samples with the following objectives:

a. obtaining a comprehensive view of the bacterial and fungal microbial communities of table olives, and understanding the variability due to systematic effects, like olive variety, ripeness and trade preparation, and the extent of random variability between and within producing companies.
b. using black natural Itrana olives as a case study to evaluate if detectable differences between PDO and non-PDO products exist

## 2 Materials and methods

### 2.1 Sampling

Table olive samples (n = 340) were obtained from 40 producers across 6 countries (Italy, Greece, Spain, Cyprus, Peru, Egypt). Producers were required to deliver olives ready for packaging in their fermentation brine,, and to fill a questionnaire reporting essential metadata (location of the company, olive ripeness and variety, the techniques used for transformation, addition of starter cultures, if any; spoilage; duration of fermentation and of storage post fermentation, if applicable). None of the samples used in this study had undergone a heat treatment. Twenty sample, for which the fermentation brine was not available, were dispatched in the final packages, with the storage brine. Furthermore, 23 metagenomic DNA from table olives were obtained from Instituto de la Grasa, CSIC (Seville, Spain) and used for amplicon targeted metagenomics, in order to increase coverage of Spanish varieties. Summary data are shown in Table 1, while detailed data are shown in Supplementary Table 1. All samples were dispatched in insulated containers with ice packs and reached our laboratory within 48 (samples from Italy) to 72 h. They were stored at 4°C for no more than 24 h before being used for microbiological analyses and extraction of DNA.

**Table 1.**
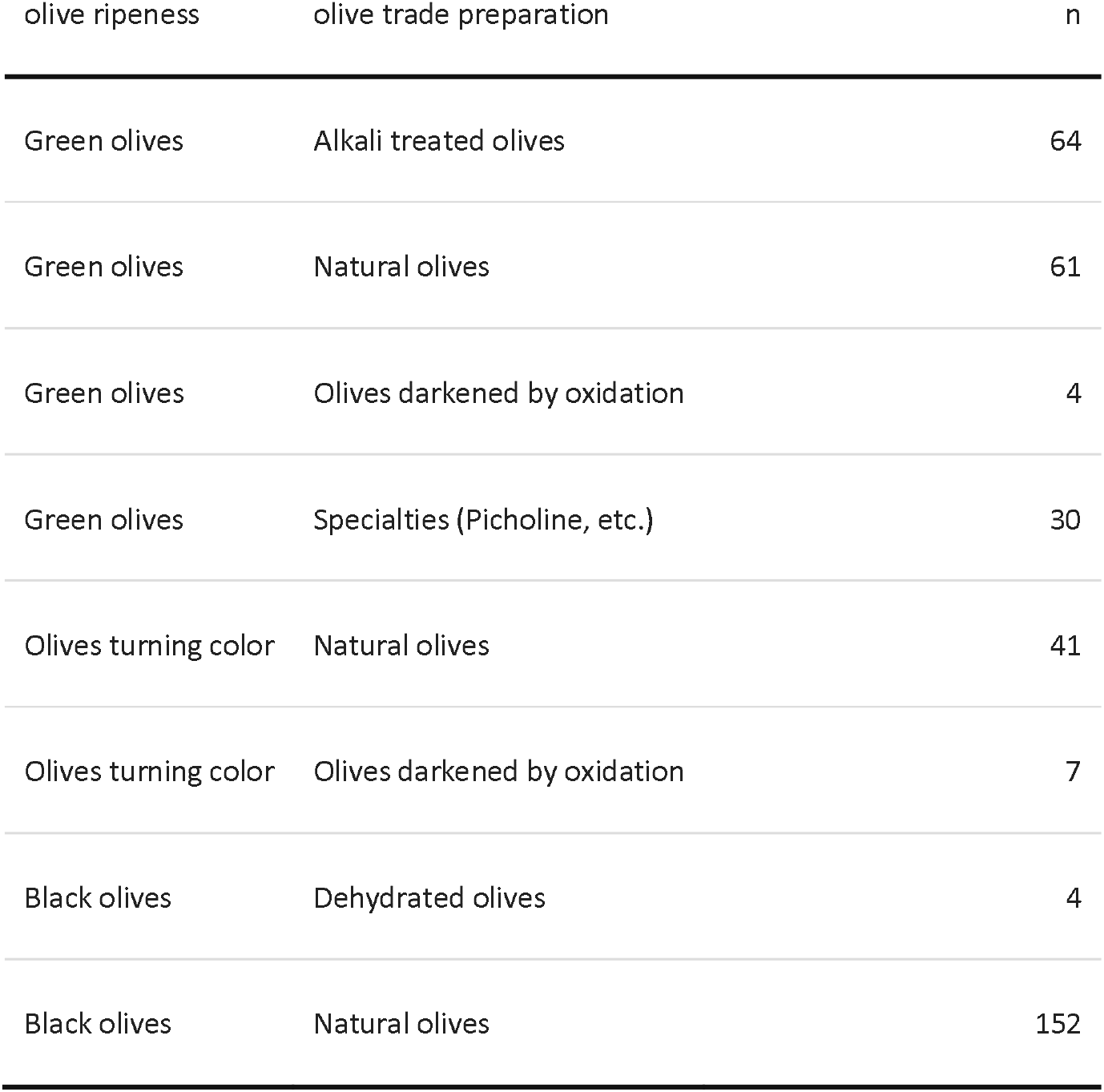
The samples used in this project, by olive ripeness and trade preparation.

| olive ripeness | olive trade preparation | n |
| --- | --- | --- |
| Green olives | Alkali treated olives | 64 |
| Green olives | Natural olives | 61 |
| Green olives | Olives darkened by oxidation | 4 |
| Green olives | Specialties (Picholine, etc.) | 30 |
| Olives turning color | Natural olives | 41 |
| Olives turning color | Olives darkened by oxidation | 7 |
| Black olives | Dehydrated olives | 4 |
| Black olives | Natural olives | 152 |

Supernatants (12,000 x g) of the brine were stored at -20°C for 7-30 gg before the measurement of pH, titratable acidity and chlorides.

### 2.2 Chemical and physicochemical analyses

pH and chlorides were measured in brines except for samples of dehydrated olives; the latter were homogenized (1:4) with deionized water in a Stomacher Lab Blender for 1 min using a filter bag, and pH and chlorides were measured in the filtered liquid phase. pH was measured using a digital pH meter (CyberScan pH 110, Eutech Instrument Pte Ltd., Thermo Fisher Scientific, Monza, Italy) with a combination electrode (Double Pore Slim electrode, Hamilton Company, Reno, Nevada, USA). NaCl concentration was estimated from the measurement of chlorides on diluted (D = 10-50) brines using a selective electrode (Crison ISE Chlorid 51340400) with a reference electrode (Crison 5241, KNO_3_ 1 mol/L as a reference electrolyte), connected to an Orion 420 A plus Ion-analyzer. 5 mol/L NaNO_3_ was used as ionic strength adjuster (1 ml / 50 mL diluted brine) and a calibration curve (0.001-0.1 mol/L NaCl). Titratable acidity was only measured in brines using phenophthalein (Sigma) as an indicator using NaOH 0.1 mol/L and a digital burette (Brand Titrette, Brand, Wertheim, Germany).

Two technical replicates were used for each measurement.

### 2.3 Enumeration of microbial groups, isolation and identification

Counts were performed on a 1:1 mixture of olives (pits were aseptically removed) and brine after homogenization (2 min) in a filtered bag (Nasco Whirlpak, Pleasant Prairie, WI) using a Stomacher Lab Blender 400 (Cole-Parmer, now Antylia Scientific, Vernon Hills, IL, USA) in peptone water (Bacteriological peptone 0.1% w/v, NaCl 0.85% w/v).

Serial ten-fold dilutions were performed in peptone water. Lactic acid bacteria were enumerated in modified MRS agar (Lee and Lee, 2008; Ricciardi et al., 2014) with 0.02% sodium azide (mMRS+NA, 25°C, 72 h), yeasts and moulds on Glucose Yeast Extract agar with 100 mg/L chloramphenicol (GYEAC, 25°C, 5 days), halophilic microorganisms on Plate Count Agar Standard with 5% NaCl (w/v) (PCA + NaCl, 30°C, 48 h), contaminants on *Pseudomonas* agar base (PAB, 25°C, 48 h), and *Enterobacteriaceae* in Violet Red Bile Glucose Agar (VRBGA, 30°C, 24 h). All microbiological media were inoculated using a spiral plater (WASP Spiral Plater, bioMérieux Italia SpA, Bagno a Ripoli, Italy) with the exception of VRBGA, for which conventional pour plating was used. Colony counts were performed using a digital colony counter (EasyCount 2, bioMérieux Italia).

Colonies (1-2 from at least one counting plate) were isolated at random and streaked for purification on the same medium and, after control of cell morphology by light microscopy, identification was carried out with MALDI-ToF Biotyper Sirius (Bruker Co, Mallerica, MA, USA) using the “Extended Direct Transfer” SOP provided by the producer.

### 2.4 Microbiological media, ingredients and reagents

All microbiological media and ingredients were obtained from Oxoid (Basingstoke, United Kingdom) and reagents from Sigma-Aldrich (Merck KGaA, Darmstadt, Germany).

### 2.5 Amplicon targeted metagenomics

Metagenomic DNA was extracted from both olives and brines (when applicable) in an attempt to recover both planktonic microorganisms and microorganisms attached to olive surfaces, while minimizing contamination with fruit DNA. Olives (40 g) were mixed with an equal volume of brine (for dehydrated olives, sterile distilled water) to which sterile Tween 80 (final concentration 0.1% vol/vol) was aseptically added. After shaking for 10 min on a rotary shaker (100 rpm) in ice, the liquid was centrifuged (8,000xg for 10 min) and the pellet was washed twice in sterile NaCl 0.85% and immediately frozen at -80°C. DNA extraction and purification were performed using DNeasy PowerFood Microbial Kit (QIAGEN Italia, Milan, Italy) followed by quantitation using NanoDrop One spectrophotometer (Thermo-Fisher Scientific, Milan, Italia). Library preparation and sequencing was performed by Novogene (Cambridge, UK). The V3-V4 region of the 16S rRNA gene was amplified using Novogene’s proprietary primers Bakt_341F_Nov (5’-CCTAYGGGRBGCASCAG-3’) and Bakt_805R_Nov (5’-GGACTACNNGGGTATCTAAT-3’). The ITS2 region was amplified using primers ITS3F (5’-GCATCGATGAAGAACGCAGC-3’) and ITS4R (5’-TCCTCCGCTTATTGATATGC-3’) for Fungi (White et al., 1990). Paired end sequencing (2×250 bp) was performed on Illumina NovaSeq X. Raw sequences were processed using a pipeline based on DADA2 (Callahan et al., 2016) using R (R Core Team, 2025). Model scripts are available at https://github.com/ep142/FoodMicrobionet/tree/master/dada2_pipeline. Taxonomic assignment was carried out using functions assignTaxonomy() and addSpecies(), which implement the naïve Bayeasian classifier (Wang et al., 2007), with SILVA 138.2 (Quast et al., 2013) for bacteria and UNITE v10 general release 2024-04-04 for fungi (Abarenkov et al., 2023) as taxonomic reference databases. The results were assembled in a phyloseq object (McMurdie and Holmes, 2013) which was then used for further analysis. Extraction blanks and a mock community (ZymoBIOMICS Microbial Community DNA Standard, Zymo Research, Invine, CA, USA) were also included. Contaminating ASVs were removed using package “decontam” (Davis et al., 2018). Non-target sequences were removed prior to further statistical processing using phyloseq::subset_taxa().

### 2.6 Statistical analysisi

Statistical analyses and graphs were produced with scripts combining the functions of the packages stats (R Core Team, 2025), PCAtools (Blighe and Lun, 2025), phyloseq (McMurdie and Holmes, 2013), ggplot2 (Wickham, 2016), ggpubr (Kassambara, 2025) and rstatix (Kassambara, 2023). Non-parametric tests (Wilcoxon-Signed Rank Sum test) were used for comparing distributions of physico-chemical and chemical variables and counts (as log(cfu/g brine and olives)). Holm correction for multiple testing was used. Alpha diversity analyses, ordinations and inferential analyses on microbial community data were performed using functions of the packages phyloseq (McMurdie and Holmes, 2013), DESeq2 (Love et al., 2014) and vegan (Oksanen et al., 2025). Heatmaps were generated using package ComplexHeatmap (Gu, 2022).

To account for unequal sequencing depth, rarefaction was performed before calculation of alpha-diversity. The threshold for rarefaction (15,000 and 20,000 sequences for bacteria and fungi, respectively) was chosen based on rarefaction curves (data not shown). Total Sum Scaling was performed before ordinations and PERMANOVA.

## 3 Results

### 3.1 Sampling

A summary of the samples grouped according to ripeness and trade preparation is shown in Supplementary Table 1. Most of the samples (natural olives, 44%, Spanish-style, alkali treated Spanish-style olives, 19%) had undergone a fermentation step, while 27% had undergone an alkali treatment. The latter included both Spanish-style olives and specialties like Nocellara del Belice produced with the “Castelvetrano” method (Perpetuini et al., 2020). The remaining samples were produced with other methods, including drying. Spoiled samples were also made available by some producers (Supplementary Table 3).

### 3.2 pH, titratable acidity and salt content

The pH of brines and olives in our study ranged from 3.7 to 8.1 (see Supplementary Table 4, Supplementary Table 5 for summary statistics). Even within a single variety, the pH was highly variable (see Supplementary Figure 1 and Supplementary Table 5 for ranges and standard deviation values). The null hypothesis of no differences in pH distribution among groups defined by combination of ripeness and trade preparation was rejected (Wilcoxon rank-sum test, WSRT, with multiple-testing correction; p < 0.001). Black dehydrated olives and specialties showed a higher median pH.

Results for titratable acidity (TA) of brines reflect those of pH with the exception of olives darkened by oxidation, to which gluconate was added. A box plot of the distribution of TA for the main varieties is shown in Supplementary Figure 2, while summary statistics are reported in Supplementary Table 6: the titratable acidity ranged from 0.3 to 24 g lactic acid/L, with significantly higher median values (WSRT) for natural olives and significantly lower values for alkali treated olives and specialties in particular. Again, high variability was observed even within the same variety as shown by the spread of the points in Supplementary Figure 2. There was a weak (-0.58) but highly significant (p<0.001) correlation between pH and the log_10_(TA).

Summary statistics for salt content in brines and olives are shown in Supplementary Tables 7 and 8 while a box plot for the main varieties is shown in Supplementary Figure 3. NaCl concentrations ranged from 2.2 g/L to near saturation (246 g/L), whereas median values in groups of samples sharing the same ripeness stage and trade preparation varied between 23 and 82 g/L. High variability was observed. For Oliva di Gaeta samples, which were the most numerous, NaCl content of brines varied between 30 and 116 g/L. Dehydrated olives and olives darkened by oxidation had significantly lower median NaCl values (WSRT).

### 3.3 Microbial counts

Viable microbial counts were obtained using selective or elective media for undesirable microorganisms (*Enterobacteriaceae*, contaminants on PAB) and potentially beneficial microorganisms. Based on MALDI-ToF or molecular identification of isolates (Supplementary Table 9) we found that both mMRS+NA and GYEAC were likely to be selective (all isolates were LAB or yeasts, respectively), while PAB (Supplementary Figure 5) supported the growth of yeasts (which formed pinpoint colonies) and PCA+NaCl supported the growth of both yeast and lactic acid bacteria together with staphylococci and other halophiles. The latter is therefore of limited use when counts of yeasts and LAB are high.

Counts of *Enterobacteriaceae* are shown in Supplementary Figure 4. For most samples counts were below the detection limit (<10 cfu/mL), while values > 100 cfu/g (brine + olives) were found in several samples of naturally fermented or alkali treated olives. Counts of LAB are shown in Figure 1 while summary statistics are shown in Supplementary Tables 10 and 11. Several samples (30%) had viable counts below the detection limit of our method (200 cfu/g), with 10^9^ cfu/g detected as maximum counts. As for other variables, variability within olive varieties was very high but some varieties showed significantly (p<0.05) higher (WSRT) counts compared to others (for example, Bella di Daunia vs Halkidiki for Green alkali treated olives; Kalamata and Konservolea vs Oliva di Gaeta for black naturally fermented olives). High counts of LAB were also found in alkali treated olives which are stored at low temperature after the alkali treatment, like Nocellara del Belice produced with the Castelvetrano method (Alfonzo et al., 2024).

**Figure 1.**
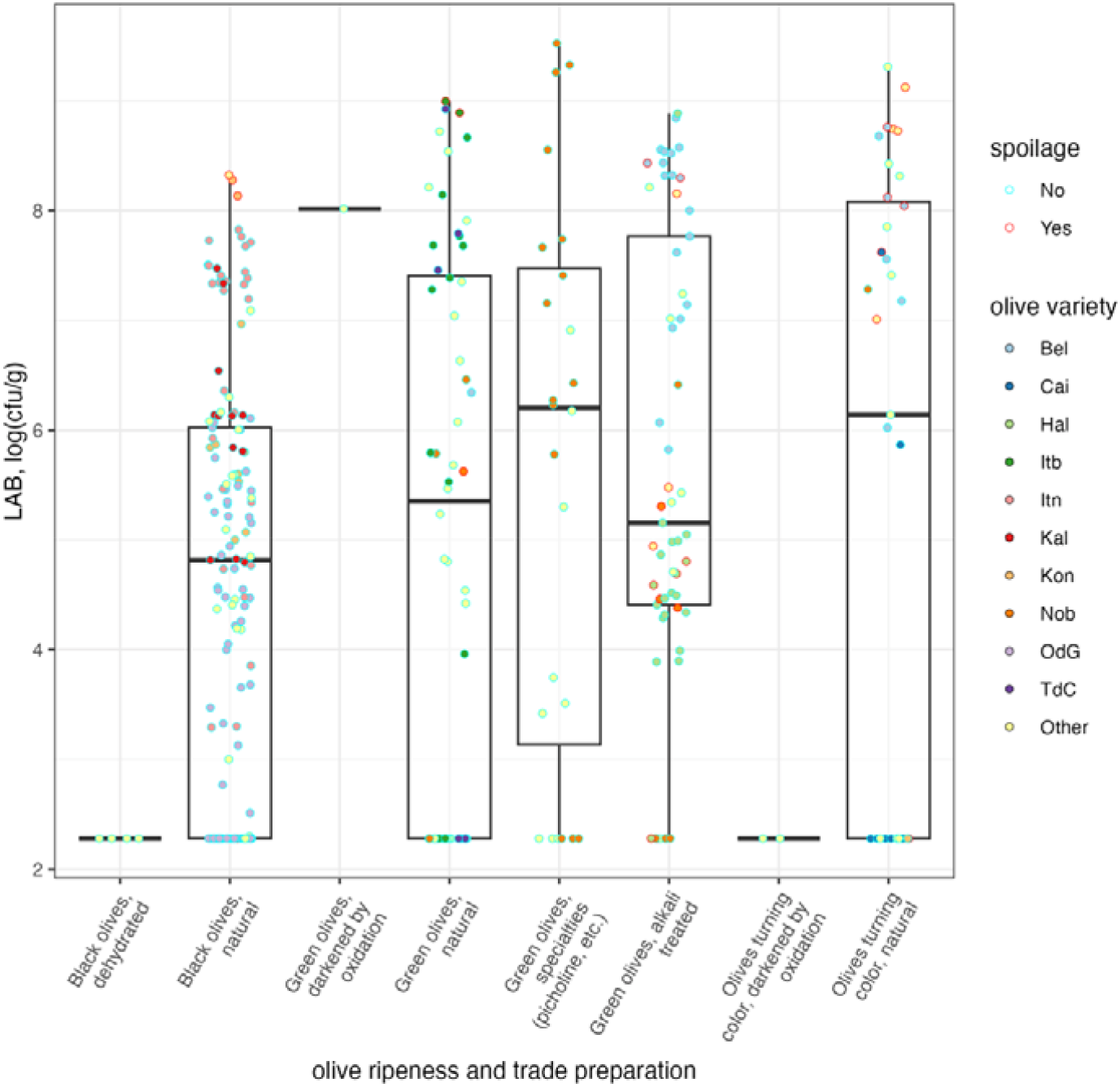
Distribution of viable counts of lactic acid bacteria in table olives used in this study.

Summary results for yeast counts (moulds were only occasionally detected) are reported in Supplementary Tables 12 and 13, and their distribution is shown in Figure 2: median values for the groups defined by combinations of ripeness and trade preparation ranged between 5.02 and 6.85 log(cfu/g) and only 6% of the samples had counts below the detection limit of the method while counts exceeding 10^8^ cfu/g were observed thus confirming that yeasts are among the main microbial groups in table olives. In fact, counts exceeding 10^5^ cfu/g were observed even in varieties which do not undergo fermentation, like Nocellara del Belice, dried olives and olives darkened by oxidation.

**Figure 2.**
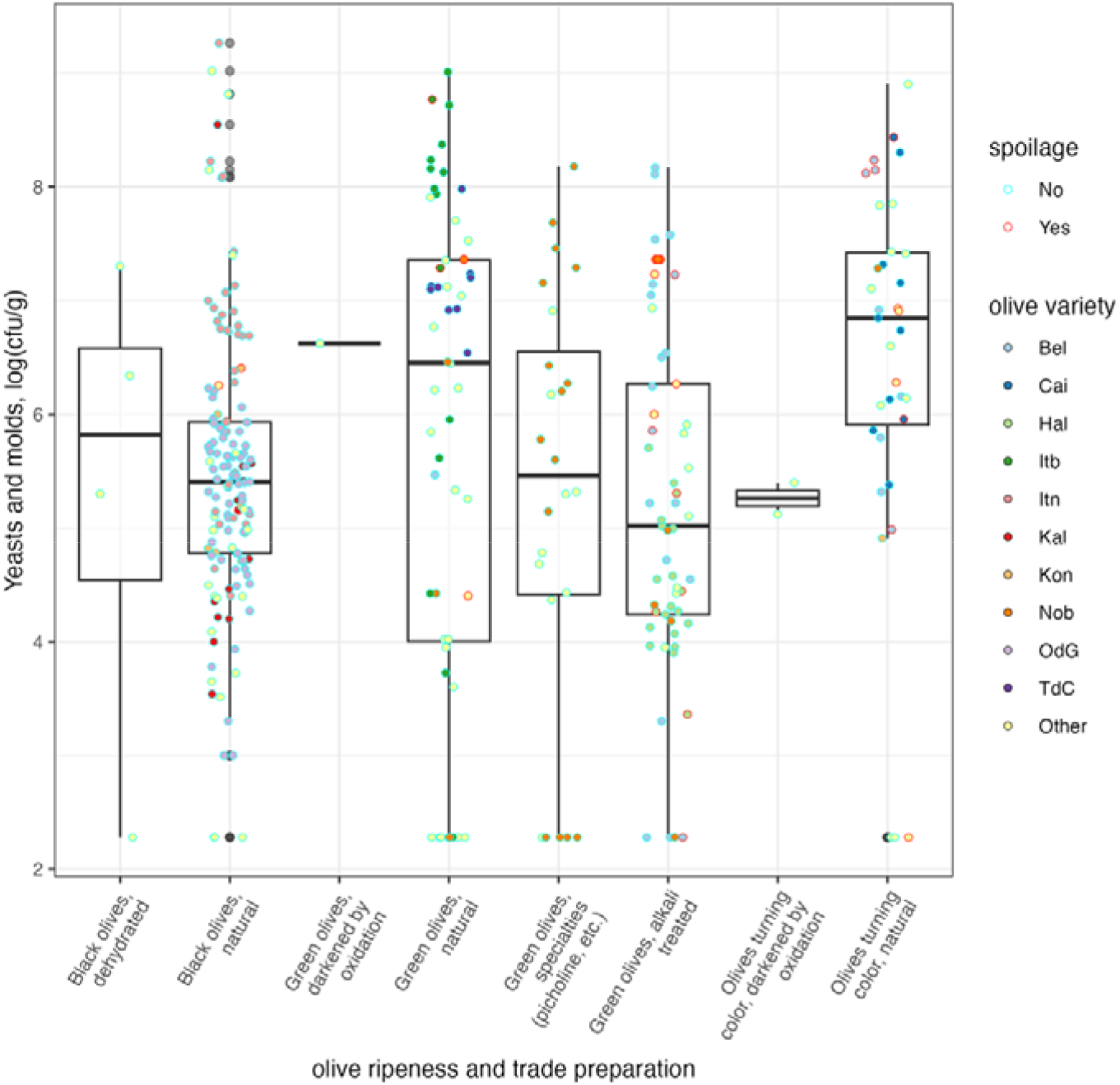
Distribution of viable counts of yeast and moulds in table olives used in this study.

To explore the correlation structure among counts and chemical and chemico-physical parameters, Pearson’s correlation matrix was estimated and a correlation test with Holm’s correction for multiple testing was carried out to test the null hypothesis that correlation coefficients were not significantly different from 0. All coefficients were <|0.5|, with the highest absolute values for the correlation between counts of contaminants and halophilic microorganisms (0.45) and pH and titratable acidity (-0.37) although many coefficients were significantly different from 0. As a result of the general lack of correlation structure, the first two components of a Principal Component Analysis carried out on the correlation matrix explained only 43% of the variance, with no clear separation between different olive groups (Supplementary Figure 6). This again, might be the result of the very high variability within groups.

### 3.4 The composition of microbial communities of table olives

#### 3.4.1 Alpha-diversity of microbial communities

Extraction of DNA proved to be challenging for some low biomass samples, but we were able to carry out sequencing of the V3-V4 region of the 16S rRNA gene (336 samples) and of the ITS2 region (337 samples) of most of the samples. Mock communities (2) and blanks (3) were included for quality control. The number of sequences after QC filtering, ASV inference and chimera removal was 17.2×10^6^ for bacteria and 17.0×10^6^ for fungi. 90% of the samples had 29.3 x10^3^ sequences or more for bacteria, while the 90^th^ percentile for fungi was 34.9 x10^3^ sequences. The ASV abundance tables for both bacteria and fungi, after removal of contaminants and non-target sequences, were highly sparse (99.3 and 99.6% for bacteria and fungi, respectively). To avoid massive loss of rare taxa during prevalence and abundance filtering and loss of power in inferential tests we decide to perform taxonomic agglomeration prior to statistical analysis rather than operating on ASVs.

Our pipeline for 16S rRNA sequences theoretically allowed taxonomic assignment down to the species level. However, this is not always possible for *Lactobacillaceae*, even when using a custom database (Parente et al., 2023). Therefore, alpha and beta-diversity analysis for bacterial communities was performed after agglomeration at the genus level. Chao1 values calculated on the bacterial communities are shown in Figure 3, while summary statistics are reported in Supplementary Tables 14 and 15.

**Figure 3.**
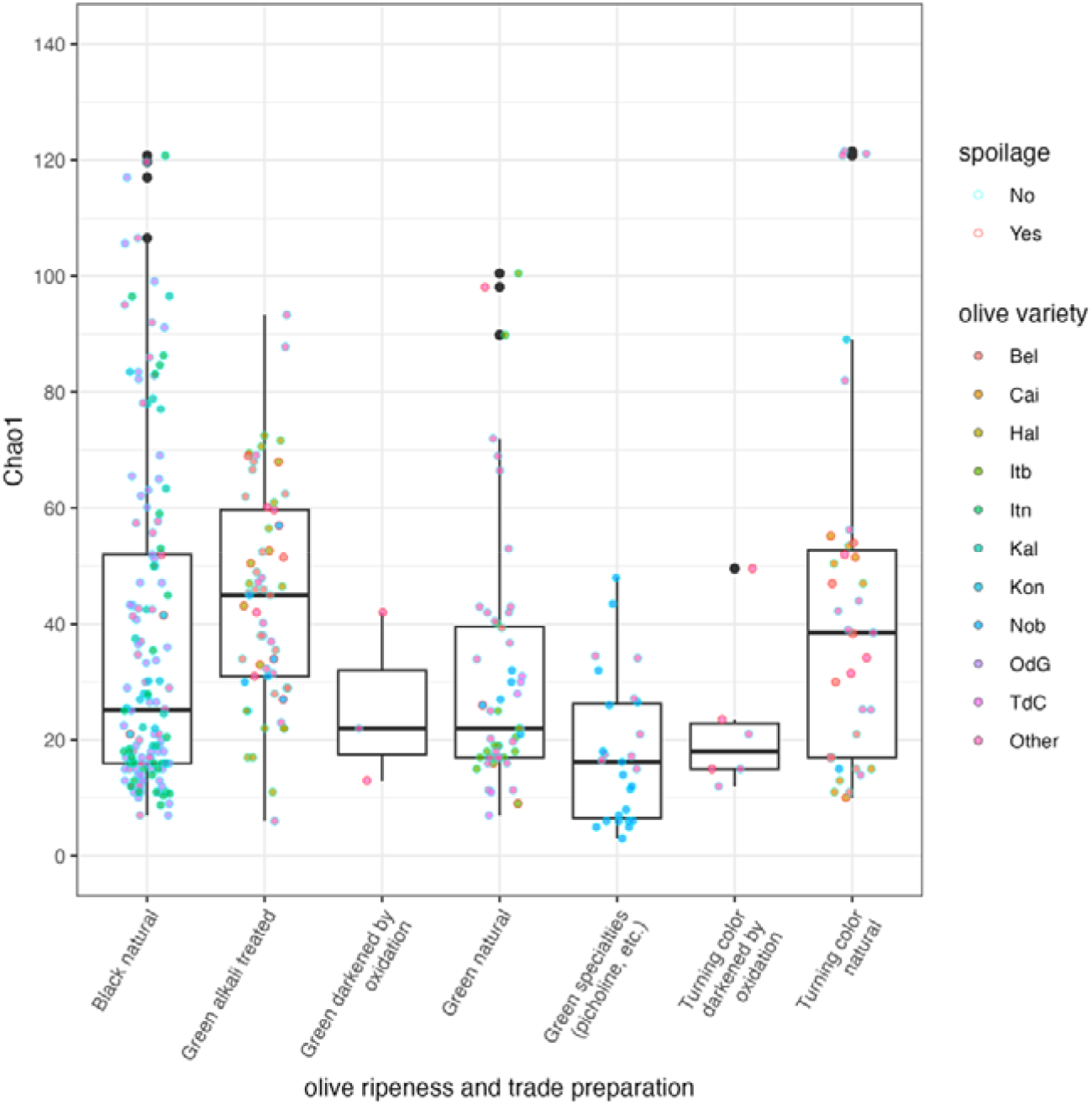
Alpha-diversity (Chao1) of bacterial communities in table olives analysed in this study.

Chao1, estimated using the estimate_richness() function of the phyloseq package after rarefaction, ranged from 3 to 21 (with median values for different groups ranging from 16 to 45). A large variability existed within each group and even within the same variety but some trends were evident. There was a trend in the decrease of diversity from Alkali treated olives, to natural olives, to olives darkened by oxidation, to specialties. We decided to investigate if between-firm and within-firm variability was a contributing factor. Distribution of Chao1 is shown for Itrana, Bella di Daunia and Halkidiki in Supplementary Figures 7, 8 and 9. Within the same variety the diversity of bacterial communities varied between different producers. We used Spearman’s ρ to evaluate correlations between diversity, physical-chemical parameters, and microbial counts for specific ripeness and trade preparation combinations (Supplementary Table 16). With the exception of a strong negative correlation between pH and diversity in Specialties - where diversity increased as pH decreased, likely due to activity of LAB - all other correlations were low (|ρ| < 0.40). While significantly different from 0, the remaining correlations often differed in sign across different trade preparations, indicating that the effect of factors like acidification or NaCl concentration on Chao1 differed in different groups.

Since our pipeline allowed assignment at the species level for the vast majority of sequences (69.4% of the ASVs, 97.7% of the sequences), diversity indices for fungi were estimated before agglomeration at the genus level. The distribution of Chao1 values is shown in Figure 4, while summary statistics are in Supplementary Tables 17 and 18.

**Figure 4.**
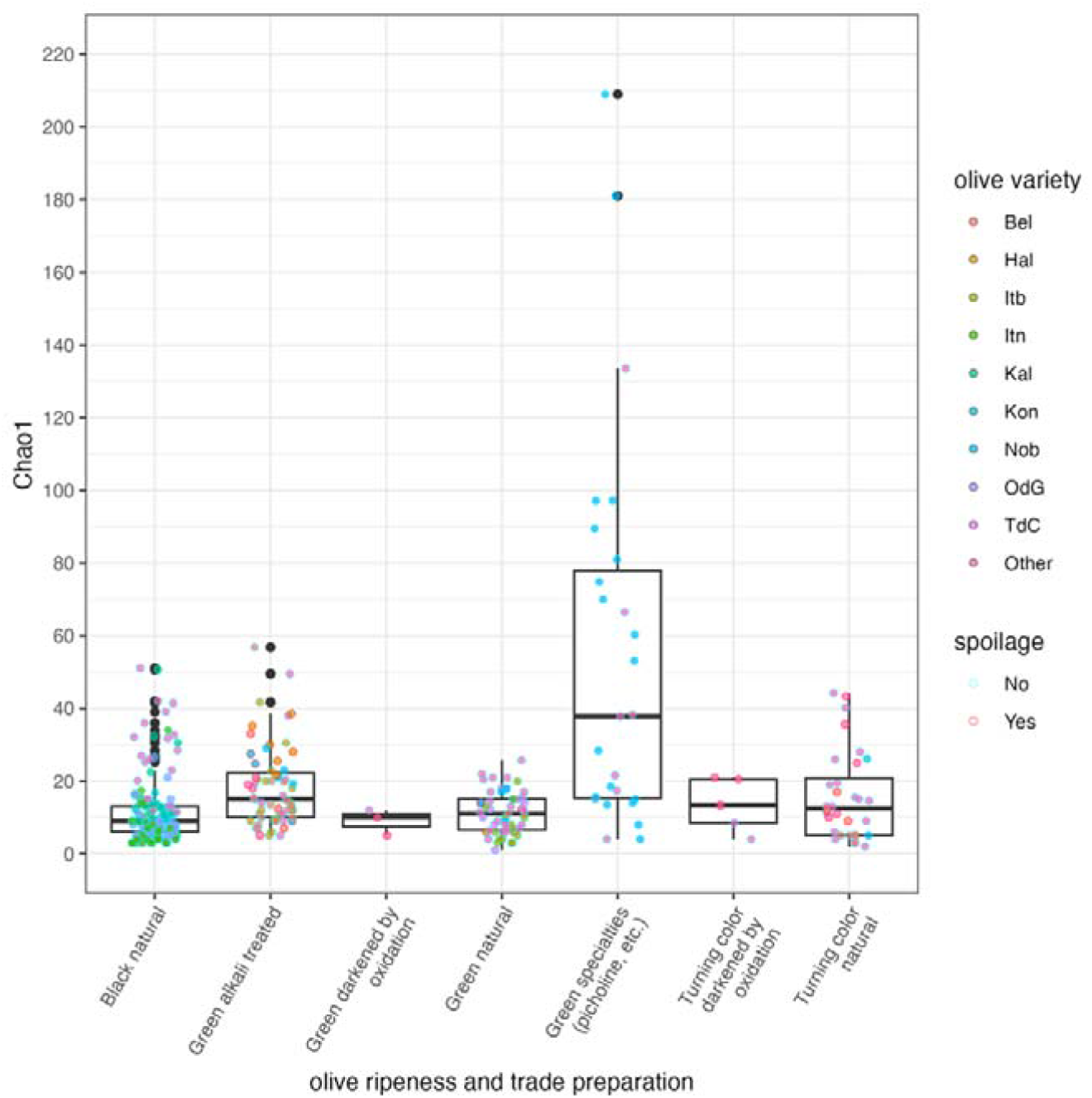
Alpha-diversity (Chao1) of fungal communities in table olives used in this study.

Chao1 ranged from 1 to 209 (with median values for different groups ranging from 9 to 41, lower than those for bacteria for most groups). A large variability existed within each group and even within the same variety, but some trends were evident: the median value for Chao1 decreased from Specialties to alkali treated olives, to natural olives (Figure 4). Significant differences among the distributions were found using WSRT for all the comparisons between alkali treated olives (including specialties) and natural olives (p<0.02). We investigated if between-firm and within-firm variability was a contributing factor. As for bacterial communities, distribution of Chao1 values within the same variety (Itrana olives, Bella di Daunia and Halkidiki olives: Supplementary Figures 10, 11 and 12) varied among different producers.

We also evaluated the correlation (Spearman ρ) between Chao1 and selected variables within groups of ripeness and trade preparation (Supplementary Table 19): a rather strong negative correlation was found between Chao1 and yeasts and moulds counts and duration of fermentation for most groups, suggesting that, as the fermentation duration increases, diversity of fungi decreases. Again, differences exist among varieties, possibly because of systematic combinations of specific factors affecting dispersal and selection: in fact, in Oliva di Gaeta, there was a weak but significant positive correlation (0.35) between pH and Chao1; as for bacteria, higher pH is associated with higher diversity.

After prevalence and abundance filtering (> 1% maximum abundance and >1% prevalence), >99.8% of the sequences were preserved for bacteria (Supplementary Figure 13). The large number of genera retained highlights the high microbial diversity found in table olives. The top 24 bacterial taxa on the basis of abundance and prevalence and are shown in Supplementary Table 20. The distribution of the abundance of LAB and HALAB is shown in Figure 5. Although abundances of the main genera are highly variable within each olive group, *Alkalibacterium* and *Marinilactibacillus* are abundant and prevalent in alkali treated olives, while a diverse set of Lactobacillaceae, among which *Lactiplantibacillus* is clearly both prevalent and abundant, dominate naturally fermented olives communities. Several heterofermentative LAB are often (*Lentilactobacillus*) or occasionally (*Limosilactobacillus, Levilactobacillus, Paucilactobacillus* and *Leuconostoc*) dominant or subdominant. It is interesting to note that, even in samples where *Lactiplantibacillus* was not dominant, *Lactiplantibacillus pentosus* was the most frequently isolated species (Supplementary Table 9).

**Figure 5.**
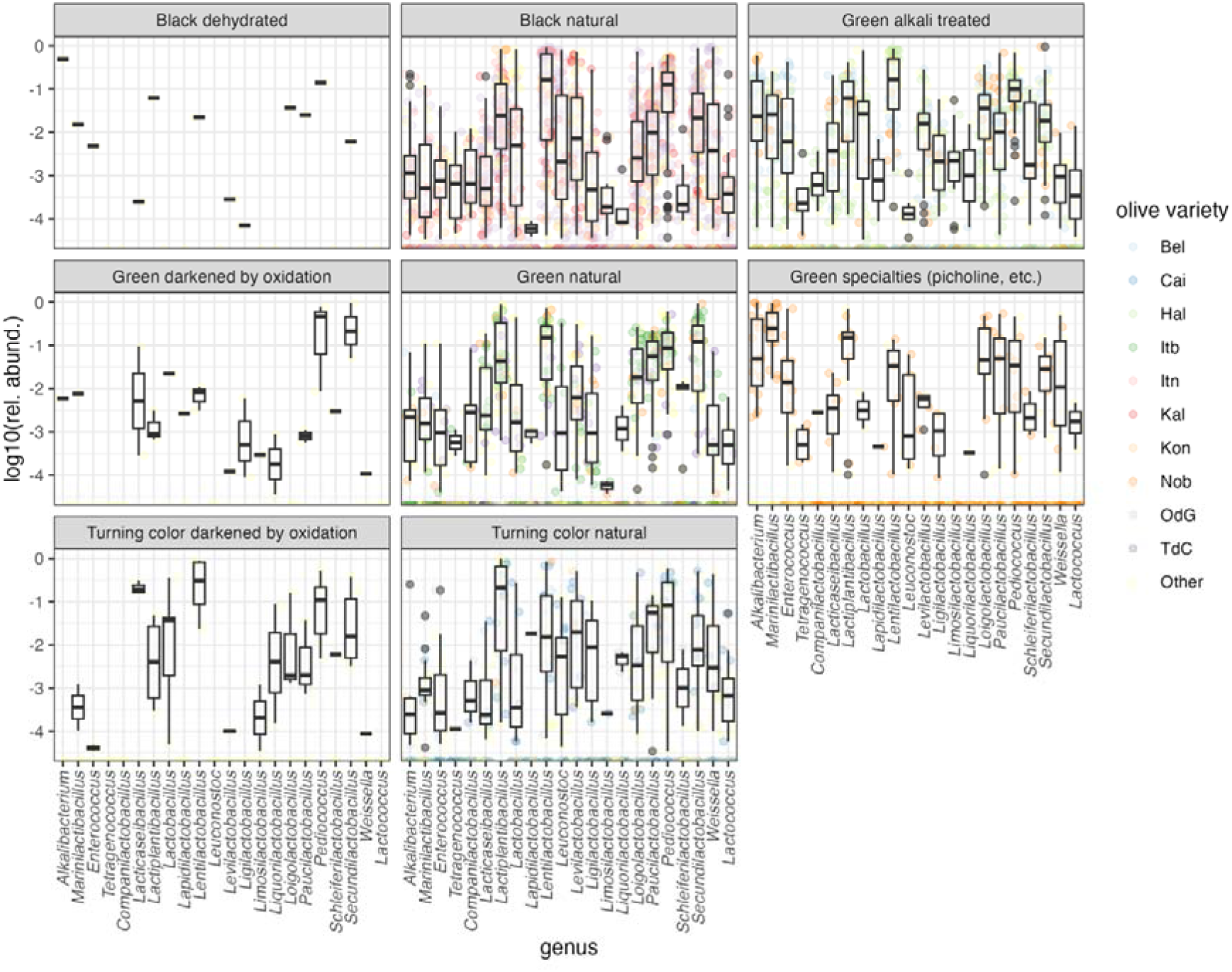
Distribution of the relative abundance of LAB and HALAB in table olives. Genera are ordered by alphabetical order within family.

The distribution of non-LAB halophilic genera is shown in Supplementary Figure 14. *Celerinatantimonas* was the most prevalent and abundant halophilic *Pseudomonadota*, but other Gram-negatives, such as *Halomonas, Vibrio, Salinicola* and *Idiomarina* were prevalent and occasionally abundant.

In a recent review we have found that DNA of genera including pathogenic bacteria can be found, albeit at very low abundance, in table olives (Ricciardi et al., 2025b). We repeated the analysis for this survey and found that only in very few samples DNA sequences assigned to genera *Salmonella, Yersinia* and Listeria were detectable, usually at very low abundance, with one notable exception (Supplementary Figure 15).

Bar plots of the relative abundances of the 24 most abundant and prevalent genera are shown in Supplementary Figures 16 (pooled data for all varieties, by ripeness and trade preparation group) and 17 to 21 (main varieties within ripeness and trade preparation groups). Some general patterns are evident: HALAB were more frequent and abundant in alkali treated olives, while LAB, mainly *Lentilactobacillus, Lactiplantibacillus* and *Pediococcus* were dominating members in naturally fermented olives. However, variety-specific patterns appear. Two examples, for Bella di Cerignola (used for different types of Bella di Daunia PDO) and Itrana (used for the production of two non-PDO types and of Oliva di Gaeta PDO) are shown in Supplementary Figures 22 and 23. In both cases, while there is some similarity in the composition of the bacterial community within a given variety, differences between firms are visually evident.

Abundance and prevalence filtering was performed on fungal communities using approaches similar to those described above. Our pipeline allowed the assignment at the species level for the vast majority of ASVs, but given the debate on the reliability of species assignment using regions of the ribosomal RNA operon in fungi (Kauserud, 2023; Rué et al., 2023; Tedersoo et al., 2022) we decided to perform this analysis both at the species and genus levels. Abundance and prevalence filtering at the species level retained 195 taxa out of 795, with 99.6% of sequences, mostly belonging to phylum Ascomycota (Supplementary Figure 24). Twenty-nine taxa had a prevalence > 0.1 and a maximum abundance > 0.01 and may be considered as the core mycobiota of table olives (Supplementary Table 10). Six species (*Pichia membranifaciens, P. manshurica, Candida boidinii, Dekkera custersiana, Wickerhamomyces anomalus, Saccharomyces bayanus*) have a prevalence >0.5 and a maximum relative abundance >0.90, but with one exception, their median abundance is <0.01: in fact, they may dominate some samples but may have a low abundance in others (see Supplementary Figure 25 for a boxplot of the distribution of core species). The most abundant species were all cultivable, and were in fact isolated from counting plates (Supplementary Table 9).

To simplify further descriptive analysis, we performed aggregation at the genus level. Core fungal genera are shown in Supplementary Figure 26 and, remarkably, they largely overlap with those found in a recent review (Ricciardi et al., 2025). To avoid losing genera which are important only in one group of olives we used the same approach used for bacterial communities (the union of the sets of taxa which passed the filter in each group was retained): 107 genera were retained. Figure 6 shows a box plot of the most important families (*Pichiaceae, Saccharomycetales_fam_Incertae_sedis* and *Phaffomycetaceae*), while other genera are shown in Supplementary Figures 27 and 28.

As for bacterial genera there is a very wide variability within each ripeness and trade preparation group and sometimes even within the same variety. However, some patterns appear to emerge: *Pichia* is by far the most abundant and prevalent genus in all types of olives which have undergone a fermentation (including green alkali treated olives) while the distribution of other important genera is more variable and often clearly multimodal, with very high abundance in some samples and very low in others. Specialties clearly have the largest variety of fungal genera, possibly due to the lower duration of fermentation/storage and the less selective conditions.

**Figure 6.**
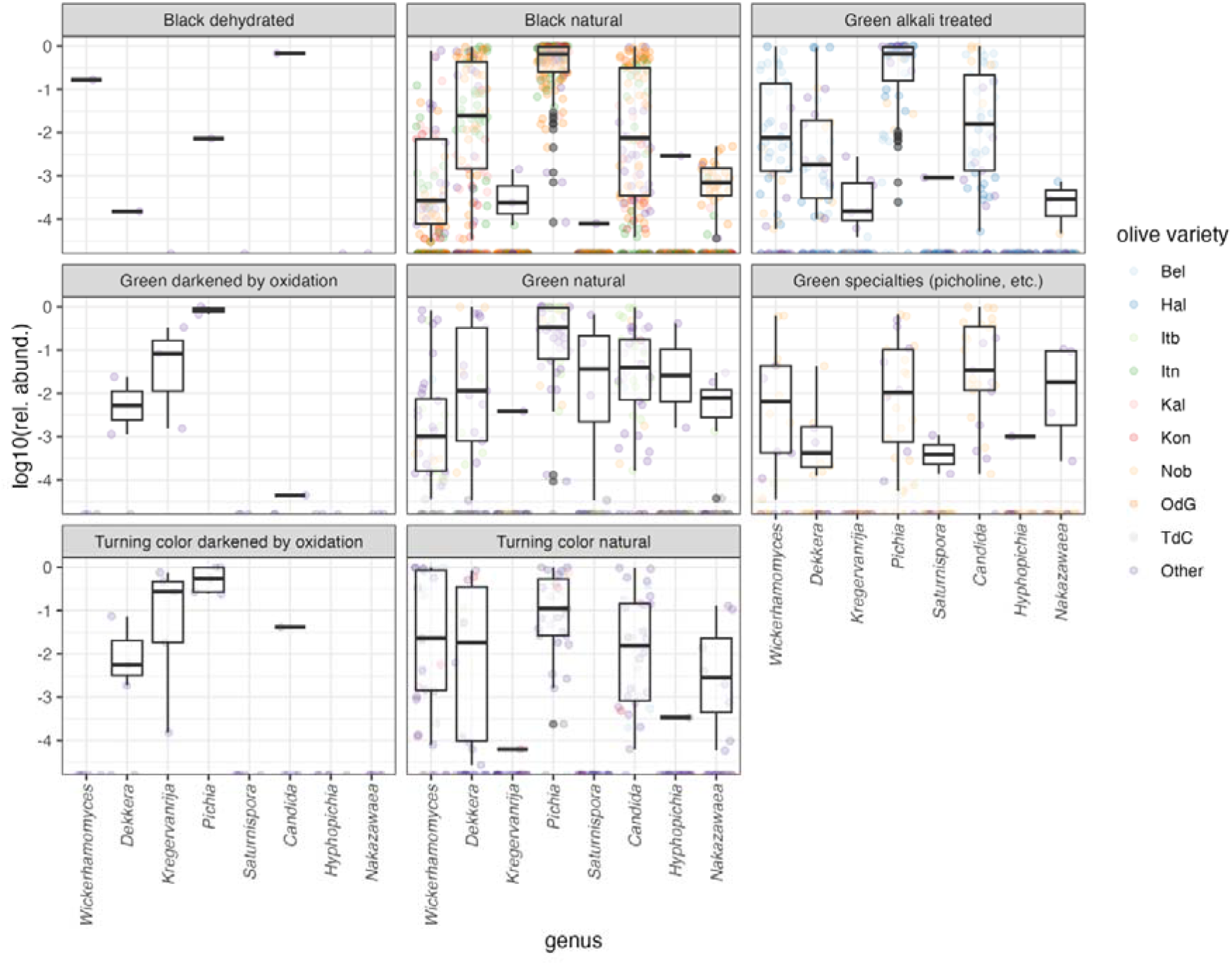
Distribution of fungal genera belonging to families *Pichiaceae, Saccharomycetales fam Incertae sedis* and *Phaffomycetaceae* in table olives used in this study. The main varieties are shown (see Supplementary Table 2 for abbreviations).

A summary bar plot with the relative abundance of the main fungal genera in ripeness/trade preparation groups is shown in Supplementary Figure 29. Overall, *Pichia* dominated most of the groups, and the sum of relative abundance of *Pichia, Candida, Wickerhamomyces, Saccharomyces, Zygotorulaspora* and *Dekkera* usually exceeded 75%. On the other hand, there are notable differences among varieties within each group (Supplementary Figures 30-34). The high fungal diversity in specialties, even within the same firm (most samples of Nocellara del Belice were provided by one firm over two years) is evident from Supplementary Figure 33. While the composition of fungal communities of samples of the same combination of olive variety and trade preparation is remarkably similar, samples from some firms have a substantially different composition (Supplementary Figures 35-36).

#### 3.4.2 Beta-diversity of bacterial and fungal communities

In order to compare the microbiota across samples and varieties, we used a four-step analytical approach. First, we calculated a Bray-Curtis distance matrix, which is a statistical measure used to quantify how much two microbial communities differ in both the types and the abundance of taxa present. Second, we visualized these relationships using non-monotonic Multidimensional Scaling (nMDS). This technique creates a spatial map where the distance between points represents their biological similarity; samples that cluster together have more similar microbiota. Third, we evaluated the homogeneity of group variances (beta-dispersion) and applied Permutational Multivariate Analysis of Variance (PERMANOVA) to test whether the differences between ripeness stages and trade preparations were statistically significant or simply due to random variation.

Finally, we carried out differential abundance analysis with DESeq2 to identify specific taxa whose abundance was significantly different is pairwise comparisons of ripeness/trade preparation groups. nMDS of bacterial communities in three dimensions resulted in a low stress (0.13), thus explaining a high proportion of the variance, and the configuration for the first two dimensions is shown in Figure 7. The variability within each ripeness/trade preparation group is evident: ripeness/trade preparation groups tend to occupy a specific area of the graph but there was an overlap between cluster of samples belonging to each group. No clear pattern for the clustering of spoiled samples (which were often, but not always, far from non-spoiled samples) is evident, but this may be due to the variety of spoilage types in our dataset. The between-firms and within-firms variability even within a single olive variety was substantial: selected examples are shown in Supplementary Figures 37-39.

**Figure 7.**
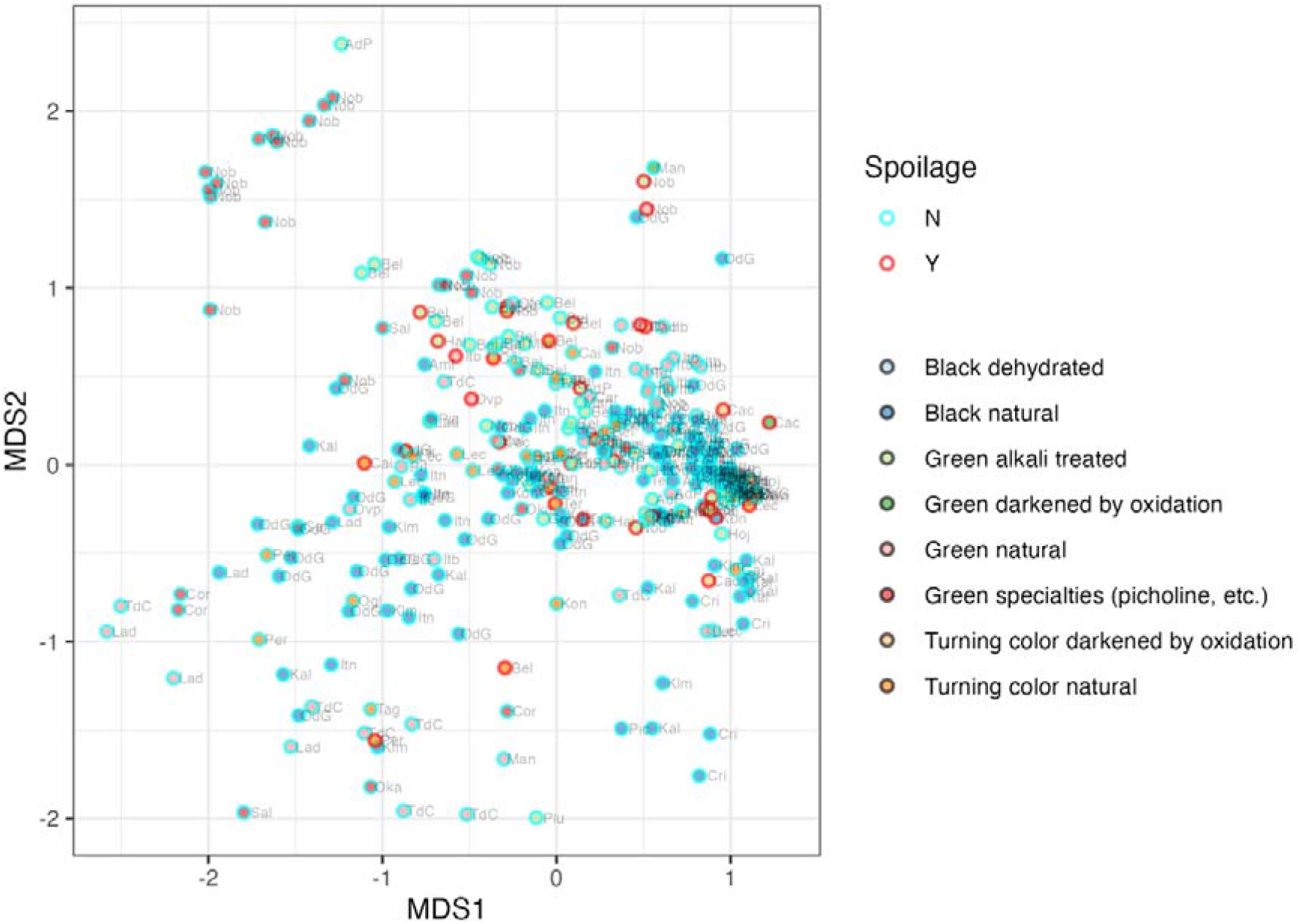
Configuration plot for the first two dimensions of a non-monotonic Multidimensional Scaling of the Bray-Curtis distance matrix of the composition of bacterial communities of table olives. See Supplementary Table 2 for abbreviations of olive varieties.

However, inferential methods (PERMANOVA with adonis2) showed that ripeness and trade preparation significantly (0.001) affected the composition of the bacterial communities. We further examined if varieties within a specific ripeness/trade preparation group or producers within a specific olive variety influenced the bacterial composition. Using PERMANOVA, we found that variety within groups significantly affected the composition of bacterial communities, except for ‘turning color’ olives that were naturally fermented (Supplementary Table 22). The producing firm was a significant factor shaping the microbiota for all varieties tested, including Itrana nera, Kalamata, Bella di Daunia and Halkidiki (Supplementary Table 23). These results indicate that house microbiota or a producer’s specific practices can create distinct microbial profiles even within the same variety. Statistical testing confirmed that the observed microbial shifts were primarily caused by true differences in community composition (group centroids) rather than simply reflecting higher variability within certain groups (beta-dispersion), with the exception of the comparison for varieties within green natural olives and firms within Kalamata olives. We also used PERMANOVA to test the null hypothesis of lack of differences in the structure of bacterial communities for Itrana nera and Oliva di Gaeta (its PDO counterpart). Olives of the two varieties were provided by 6 different firms, one of which, the largest in the PDO area, F23, provided 90% of the samples for these varieties. When only samples from F23 were used (n = 80) the null hypothesis was rejected with p = 0.025. When all firms were included, significant differences were still found but, due to unbalanced sampling and between and within firm variability, further sampling and the use supervised machine learning approaches are needed to confirm that differences in microbiota can indeed be used to differentiate PDO from non PDO Itrana olives. Finally, we used differential abundance analysis (DESeq2 with alpha = 0.01) to pinpoint the specific taxa driving the differences between groups. The results are shown in Supplementary Figures 40 to 47 and in Supplementary Table 24 and summarized in the heatmap presented in Figure 8. This approach allowed us to detect significant microbial signatures regardless of their overall rank in the community; indeed, many taxa with low total abundance were identified as key biological indicators because they exhibited large effect sizes (high log2 fold-changes) between preparations. The analysis revealed distinct microbial profiles for selected trade preparation: Alkali-treated olives were primarily distinguished by a significant enrichment of HALAB and other salt-tolerant and alkaliphilic microorganisms (*Enterococcus, Alkalibacterium, Marinilactobacillus, Halolactibacillus, Natronobacter, Alkalibacterium*). Some LAB (*Lentilactobacillus, Pediococcus, Secundilactobacillus*) were characteristically less abundant specialties and enriched in black natural olives. The latter showed a distinct prevalence of Pseudomonadota, including deleterious microorganisms like Enterobacteriaceae, and a diverse array of *Lactobacillaceae*. Specifically, heterofermentative genera—including *Levilactobacillus, Leuconostoc*, and *Weissella*—were significantly more abundant in black natural olives compared to all green varieties.

**Figure 8.**
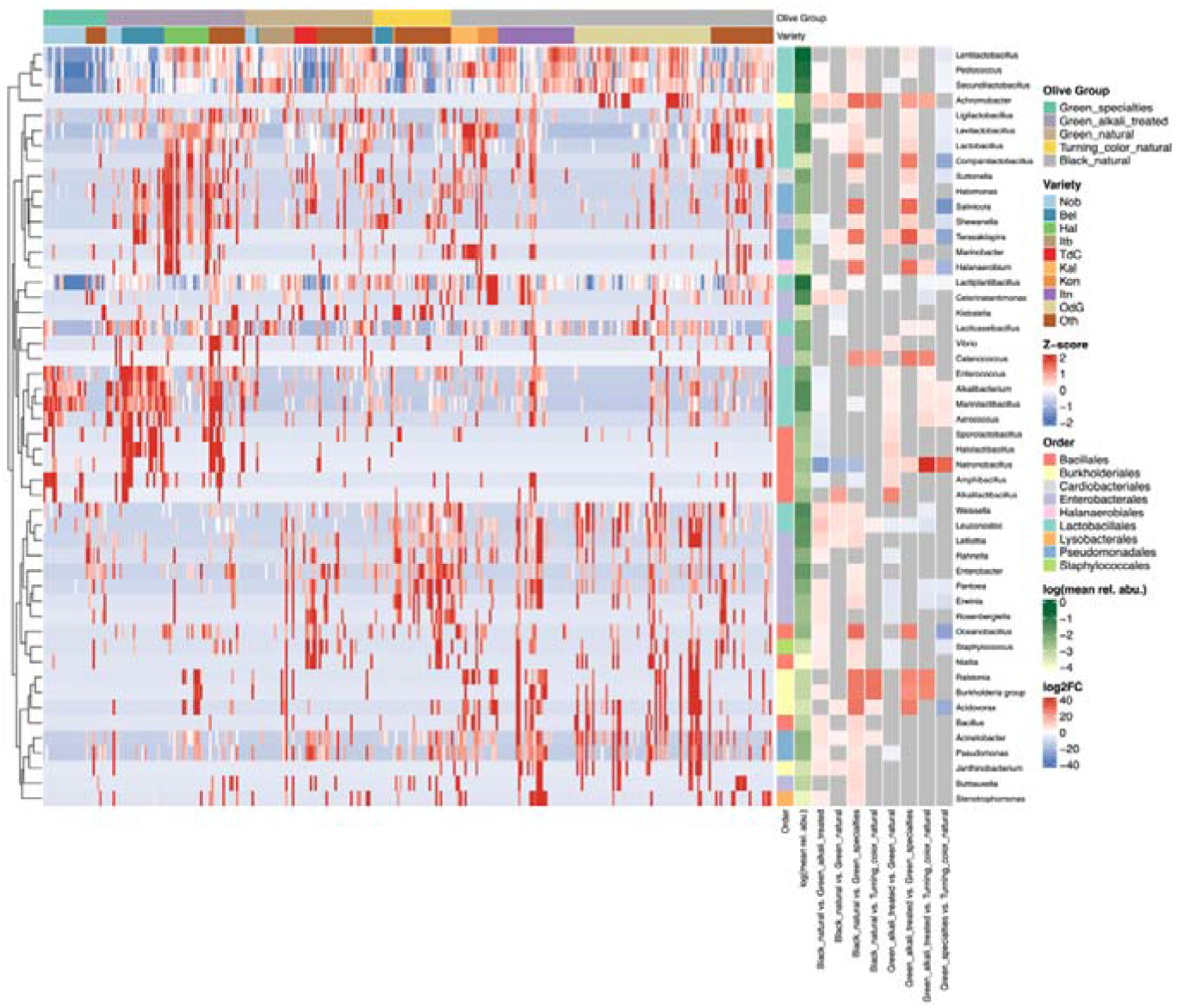
Differential abundance of bacterial genera across olive ripeness and trade preparation groups. Heatmap showing Z-score transformed variance-stabilized counts (VST) of all bacterial genera significantly differentially abundant (DESeq2, adjusted p-value < 0.01) in at least one pairwise contrast between olive groups. Columns represent individual samples grouped by ripeness/trade preparation (Green natural, Green specialties, Green alkali treated, Turning color natural, Black natural, Ripe trade) and ordered by olive variety within each group. Rows represent bacterial genera and are ordered by Spearman correlation-based hierarchical clustering. Row annotation bars (right) show the taxonomic order of each genus, the mean log10 relative abundance across all samples, and the log2 fold change for each pairwise contrast; grey cells in the log2 fold change bars indicate that the genus was not significantly differentially abundant in that contrast. Heatmap colors represent Z-scores capped at ±2 for visualization purposes; values exceeding this range are shown in the most extreme color.

We observed a clear split in the *Lactobacillaceae* family based on fruit ripeness; *Lactiplantibacillus* is significantly more abundant in black natural olives than in alkali treated green olives (including specialties), but *Lacticaseibacillus* is more abundant in Green alkali treated olives compared to Black natural.

The same approach was used for fungal communities. We first visualized community relationships using non-constrained ordination (nMDS) (Figure 9). Most ripeness/trade preparation groups exhibited overlapping, suggesting shared fungal community characteristics. By examining preparations separately (Supplementary Figures 48-50), we noted that composition of fungal is often highly variable within-variety. This was especially true for Oliva di Gaeta (Supplementary Figure 48). Although spoiled samples frequently appeared distinct within a given producer, our limited sample size for defective olives prevented definitive statistical conclusions regarding spoilage markers.

**Figure 9.**
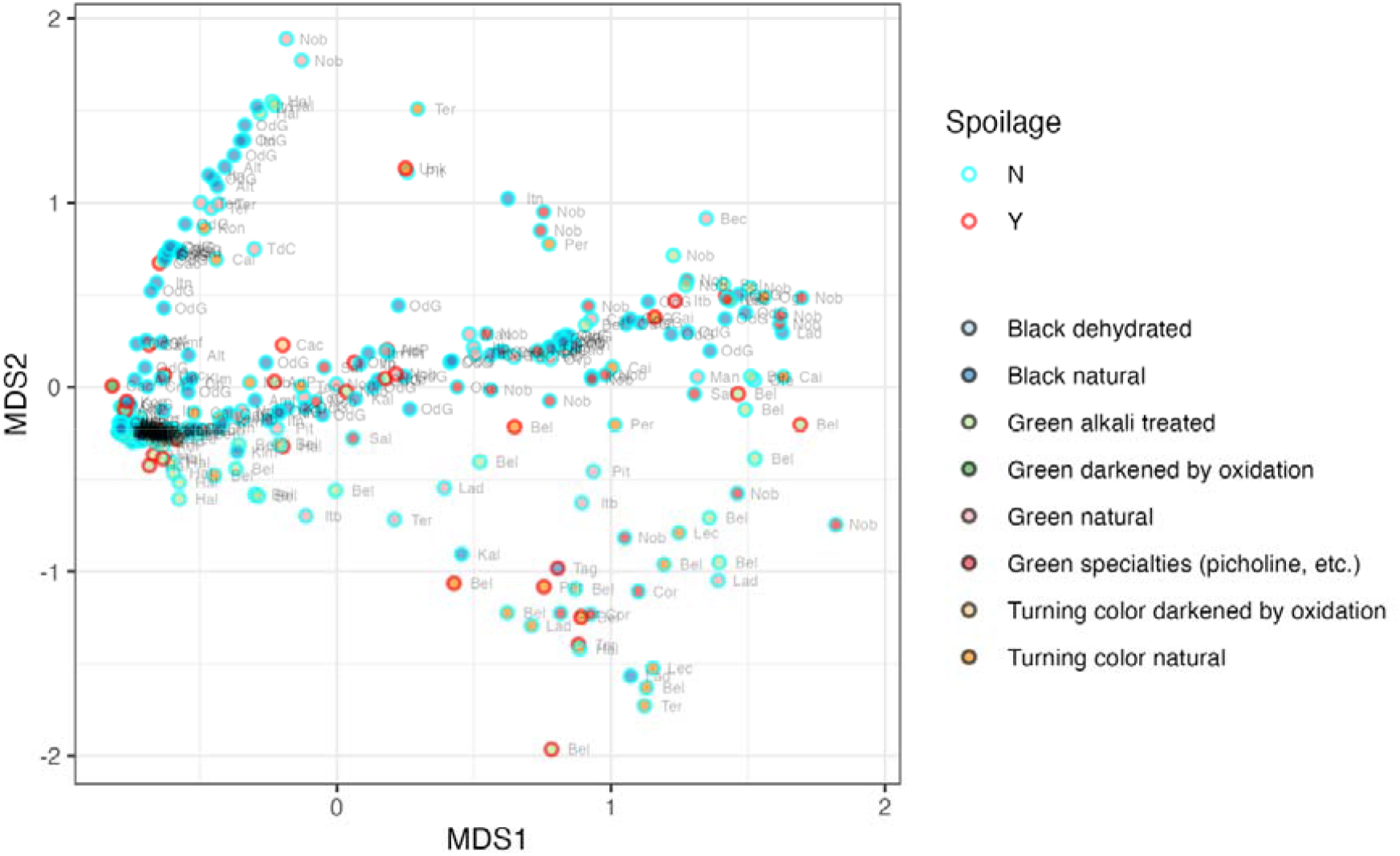
Configuration plot for the first two dimensions of a non-monotonic Multidimensional Scaling of the Bray-Curtis distance matrix of the composition of fungal communities of table olives. See Supplementary Table 2 for abbreviations of olive varieties.

As for bacteria, PERMANOVA confirmed that the combination of ripeness and trade preparation significantly shaped composition of fungal community (p = 0.001), as did the olive variety within the main groups (Supplementary Table 25).

However, in the test of effect of ripeness and trade preparation inhomogeneity of dispersion is a contributing factor, largely due to the high variance in Green and Black natural olives (see also Figure 9).

The influence of the producer was particularly prominent. For Kalamata, Itrana nera, and Halkidiki, the producing firm was a significant factor (Supplementary Table 26). This indicates that house microbiota and local practices are major drivers in determining which fungi dominate the fermentation.

As for bacteria significant differences between the fungal communities of black Itrana varieties (non-PDO Itrana nera and PDO Oliva di Gaeta) were found both when all firms were included and when only firm F23 was used (p = 0.002) thus showing that even within the same firm, systematic differences in microbiota are due to starting material and production practices.

We then carried out differential abundance analysis using DeSeq2 with alpha = 0.01. The results are shown in Supplementary Figures 50 to 57 and in Supplementary Table 27 and summarized in the heatmap shown in Figure 10.

Even with all the variability within each group, a pattern is evident. A first group of genera (*Saturnispora, Gibellulopsis, Foliophoma, Acremonium, Chytridiomycota incertae sedis, Fungi gen incertae sedis, Alternaria, Vishniacozyma, Cladosporium, Aureobasidium and Fusarium*), whose origin is likely the carposphere or the environment and whose average abundance is characteristically low, were enriched in Green specialties, in which fermentation is slowed or prevented by low temperature.

**Figure 10.**
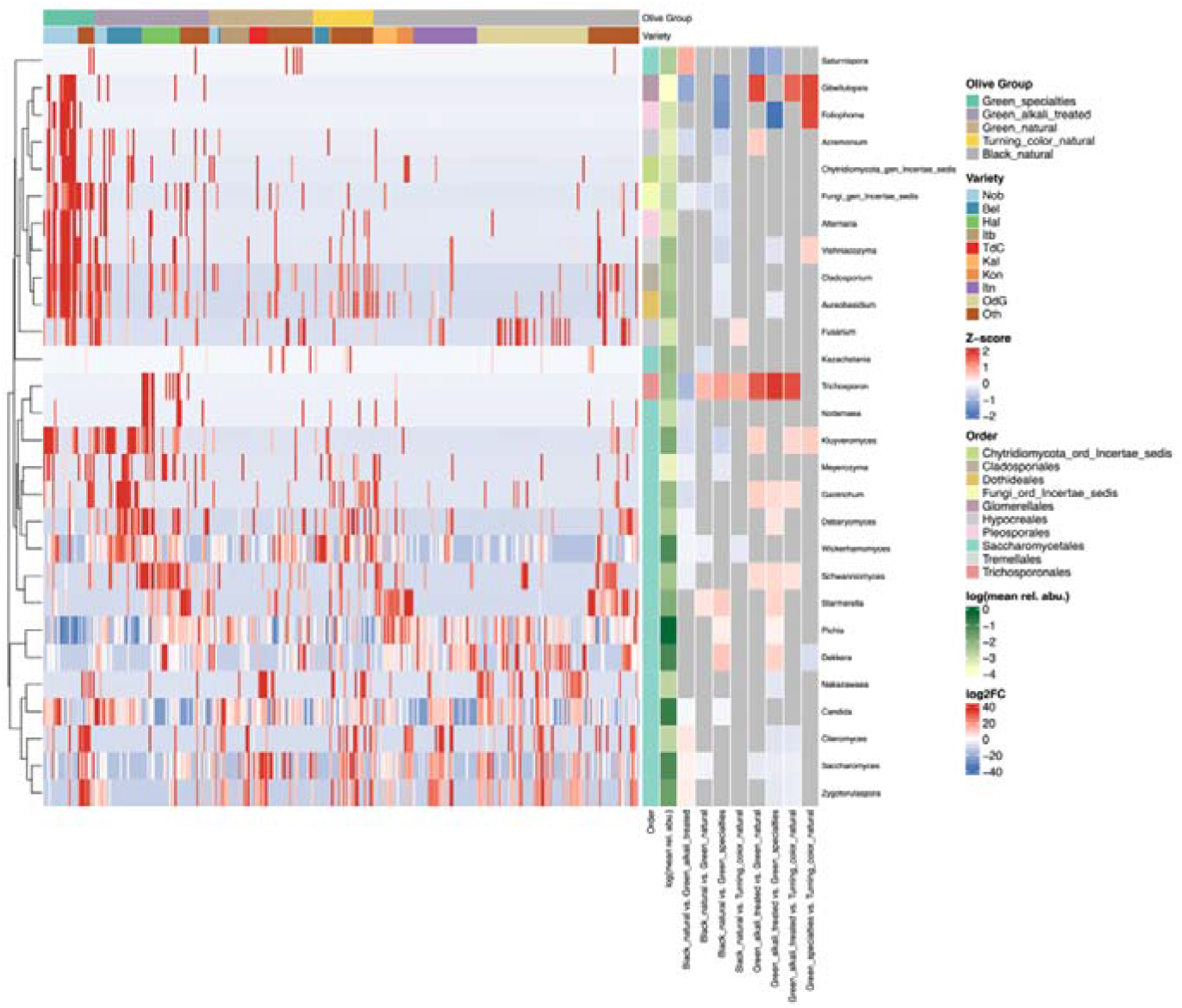
Differential abundance of bacterial genera across olive ripeness and trade preparation groups. Heatmap showing Z-score transformed variance-stabilized counts (VST) of all bacterial genera significantly differentially abundant (DESeq2, adjusted p-value < 0.01) in at least one pairwise contrast between olive groups. Columns represent individual samples grouped by ripeness/trade preparation (Green natural, Green specialties, Green alkali treated, Turning color natural, Black natural, Ripe trade) and ordered by olive variety within each group. Rows represent bacterial genera and are ordered by Spearman correlation-based hierarchical clustering. Row annotation bars (right) show the taxonomic order of each genus, the mean log10 relative abundance across all samples, and the log2 fold change for each pairwise contrast; grey cells in the log2 fold change bars indicate that the genus was not significantly differentially abundant in that contrast. Heatmap colors represent Z-scores capped at ±2 for visualization purposes; values exceeding this range are shown in the most extreme color.

A second group, which included both abundant and prevalent genera (*Trichosporon, Kodamea, Meyerozyma, Kluyveromyces, Geotrichum, Debaryomyces, Wickerhamomyces, Schwanniomyces*) was significantly more abundant in one or more alkali treated varieties compared to natural olives. The genera *Schwanniomyces* (which is abundant in Halkidiki olives) and *Starmerella* (which is abundant in Halkidiki, Konservolea and Kalamata) stand out becaue of their association with specific varieties, and this may explain its pattern in some contrasts. The last group includes genera whose pattern is more variable. *Pichia*, the most abundant and prevalent genus, is clearly less abundant in specialties but its abundance is variable in other groups. Significant differences may be obscured by the agglomeration at the genus level, since the pattern of abundance of the species within this genus varies in different groups (see Supplementary Figure 25). *Dekkera* was found to be more abundant in black and turning color natural compared to specialties. *Citeromyces* was more abundant in all types of naturally fermented olives compared to green alkali treated olives. The abundance of the genus *Candida* was lower on average in black natural compared to both green alkali treated and green specialties: however, this may not be true for all varieties and all species, since *C. boidinii* was very abundant in some samples of black natural olives (see Supplementaty Figure 25).

## 4 Discussion

This work presents the largest survey to date of the bacterial and fungal communities of table olives, with a particular focus on Italian varieties. Among these, Oliva di Gaeta — produced from naturally fermented black Itrana olives and one of the most commercially important Italian table olive varieties — is described here for the first time using culture-independent approaches, thus extending significantly previously available findings based on culture dependent approaches (Sacchi et al., 2022; Servili et al., 2006; Tofalo et al., 2012). For several other Italian PDO and non-PDO varieties, including Bella di Daunia, Ascolana del Piceno, Nocellara Etnea, Nocellara del Belíce and Tonda di Cagliari, prior metataxonomic data were limited to small, single-producer studies (Campus et al., 2025; Cocolin et al., 2013; De Angelis et al., 2015; Maoloni et al., 2022; Randazzo et al., 2017; Zinno et al., 2017), while for many other minor varieties (see Supplementary Table 2) this is the first study reporting data con bacterial and fungal communities. The dataset also includes substantial representation of Greek (Halkidiki, Kalamata, Conservolea) and Spanish (Manzanilla, Aloreña and others) varieties, providing a broader comparative framework and enabling, for the first time, a direct cross-variety analysis at this scale. A key finding emerging from this dataset is the striking variability in microbiota composition observed not only across varieties, but within the same variety and even within the same producer. This variability is consistent with a strong role of stochastic colonization events, which can obscure the selective pressures — salt concentration, ripeness, trade preparation — that might otherwise generate variety-specific microbial signatures. This has direct implications for the development of control strategies: harnessing the diversity of these communities through microbiome-based starter cultures offers a promising avenue, provided that the rich diversity of flavours and aromas that characterizes artisanal Italian table olives is preserved rather than homogenized.

We confirmed that the composition of microbial communities in table olives is shaped by physicochemical factors associated with olive ripeness, and trade preparation (Anagnostopoulos and Tsaltas, 2022; López-García et al., 2025; Perpetuini et al., 2020), which exert consistent selective pressures across producers and varieties, as confirmed by our inferential analysis.

The strongest structuring factor in our dataset was the contrast between alkali-treated and naturally fermented olives. PERMANOVA and differential abundance analysis confirmed that the combination of ripeness and trade preparation significantly affected the composition of both bacterial and fungal communities. Naturally fermented black olives were characterised by a diverse array of *Lactobacillaceae* — particularly heterofermentative genera including *Levilactobacillus, Leuconostoc* and *Weissella* — alongside a higher prevalence of *Pseudomonadota*, including *Enterobacteriaceae*. The diversity of lactic and non-lactic microbiota of natural olives and the widespread occurrence of *Enterobacteriaceae* and other *Pseudomonatota* has been confirmed by several authors (see Portilha-Cunha et al., 2020; Ricciardi et al., 2025; Tsoungos et al., 2023, for recent reviews). Among Pseudomonadota *Celerinatantimonas* is especially notable because of its spoilage potential (de Castro et al., 2022) and its very strong association with black natural olives (Kazou et al. 2020; Penland et al., 2020, 2021), a pattern which is confirmed by our data (Supplementary Figure 14, Figure 8). Alkali-treated olives, by contrast, were distinguished by enrichment of Halophilic and Alkalophilic Lactic Acid Bacteria (HALAB), including *Enterococcus, Alkalibacterium, Marinilactobacillus, and Halolactibacillus*. Within alkali-treated olives, a further important distinction emerged between Spanish-style olives, which undergo fermentation after lye treatment, and Castelvetrano-style olives, in which low-temperature storage suppresses fermentation (Alfonzo et al., 2024). The signature presence of these genera in alkali treated olives has been confirmed by several authors (Arroyo-López et al., 2021; López-García et al., 2022; Lucena-Padrós and Ruiz-Barba, 2016) and emerges as a very robust signal in a recent study using machine-learning approaches (López-García et al., 2025), especially when these are compared with directly brined counterparts of the same variety (Ruiz-Barba et al., 2023bb). The latter were characterised by the most consistent dominance of HALAB and, strikingly, by the highest fungal diversity in the dataset, with enrichment of fungal genera which have been associated with the carposphere and the environment (Ferrocino et al., 2025; Penland et al., 2021)— a pattern consistent with the less selective conditions prevailing in the absence of fermentation-driven community succession, which in naturally fermented olives progressively reduces fungal diversity as fermentation time increases. These broad contrasts provide the interpretive framework within which finer-grained effects — olive variety, producer identity, and stochastic colonization — should be understood. However, even within the same variety and production process, significant differences between producers were observed, again supported by inferential methods. This producer-level variability is consistent with the existence of a house microbiota — a resident microbial community associated with each producer’s facility and equipment — whose composition introduces a further layer of differentiation, a well-known phenomenon in spontaneous fermentation processes (Bokulich et al., 2015; Comasio et al., 2020; Li et al., 2022), which has also been confirmed in several studies on table olives (Kazou et al., 2020, Lucena-Padrós and Ruiz Barba, 2019; Penland et al., 2021). The widespread use of small fermentation vessels (240–280 L plastic fermenters at uncontrolled temperature) likely amplifies this effect: in small, semi-closed systems, stochastic colonization events have an outsized influence on the trajectory of each individual fermentation, producing divergent communities even from nominally identical starting conditions.

Culture-based enumeration and culture-independent analysis provided complementary but not always concordant pictures of the microbial communities of table olives. Among the media tested, only mMRS+NA and GYEAC showed sufficient selectivity for LAB and yeasts, respectively, while PCA+NaCl supported the growth of both groups and is therefore of limited diagnostic value when their counts are high. LAB counts were highly variable and frequently below the detection limit, consistent with the known decline of LAB over the course of fermentation and their scarcity in high-salt brines (Benítez-Cabello et al., 2020; Sánchez et al., 2018; Traina et al., 2024). Yeast counts were more consistently high across varieties and trade preparations, including in varieties which do not undergo fermentation, confirming the ubiquity of yeasts as a major microbial group in table olives (Anagnostopoulos and Tsaltas, 2022; Arroyo-López et al., 2012; Giavalisco et al., 2024). A noteworthy discrepancy emerged between culture-based and amplicon-based results for LAB. *Lactiplantibacillus* was frequently recovered even in samples where it was not the dominant taxon according to amplicon sequencing: only 29% of samples had a *Lactiplantibacillus* fraction among *Lactobacillaceae* sequences exceeding 0.20, and only 7% exceeded 0.80, yet *Lactiplantibacillus* was the sole genus isolated in 87% of samples. Colony picking was performed randomly, so this discrepancy most likely reflects either a systematic cultivability advantage of *Lactiplantibacillus* on mMRS+NA over other LAB, and relatively low recovery of colonies from other highly abundant genera (*Leuconostoc, Pediococcus, Lentilactobacillus*, among others) may reflect the fact that these genera were not viable or able to grow at the time of enumeration and isolation; a visibility bias in colony picking due to differences in colony morphology on this medium is less likely.

*Enterobacteriaceae* counts were below the detection limit in most samples, but values exceeding 100 cfu/g were occasionally found in naturally fermented and alkali-treated olives. Metataxonomic analysis revealed the presence of DNA assigned to several *Enterobacteriaceae* genera at high relative abundance in some varieties even at the end of fermentation; however, whether this represents viable cells, viable but non-culturable forms, or residual DNA from dead cells cannot be determined from these data alone. The same argument applies to the detection of genomic DNA of pathogenic or potentially pathogenic bacteria. We confirmed our previous finding of the occurrence of sequences assigned to bacterial genera containing pathogenic microorganisms (Ricciardi et al., 2025). The occurrence of *Salmonella* at relative abundance close to 0.1% in at least two samples, together with previous reports on outbreaks caused by *Salm. enterica* from olives (Donachie et al., 2018) raises some concerns on the occurrence of this pathogen in olive producing environments although we cannot infer the presence of viable pathogenic microorganisms from this data. It is well known that the low abundance of pathogenic microorganisms in foods and the low taxonomic resolution of amplicon targeted metagenomics pose challenges to the use of metagenomic approaches in the detection of microorganisms relevant for food safety (Ferrocino et al., 2022; Wang et al., 2025).

Bacterial and fungal alpha diversity, as measured by Chao1, were both highly variable within and between groups, consistent with the high variability in community composition documented throughout this study. Bacterial diversity showed a decreasing trend from alkali-treated olives to naturally fermented olives to specialties, while the opposite trend was observed for fungi — specialties harboured the highest fungal diversity, reflecting the less selective conditions prevailing in the absence of prolonged fermentation. Fermentation-driven acidification and the progressive dominance of competitive fermentative taxa reduce fungal diversity over time, as supported by the significant negative correlation between fungal Chao1 and fermentation duration observed across most groups. However, previous findings on the effect of fermentation time on diversity of fungal communities are contradictory (López-García et al., 2024; Ruiz-Barba et al., 2023a, 2023b).

Producer identity was a significant contributor to within-variety variability in diversity for both bacteria and fungi, consistent with the house microbiota effect discussed above. Chao1 values for both bacteria and fungi were broadly comparable to those reported for similar varieties and trade preparations in previous studies (Cocolin et al., 2013; De Angelis et al., 2015; de Castro et al., 2018; Kamilari et al., 2023; Maoloni et al., 2022; Medina et al., 2016; Penland et al., 2020, 2021), although differences in bioinformatic pipelines and diversity indices limit direct comparisons.

Among LAB, *Lactiplantibacillus* was the most prevalent and abundant genus in naturally fermented olives, consistent with its well-documented adaptation to the brine environment and its widespread use as a starter and functional culture in table olives (Giavalisco et al., 2024; Portilha-Cunha et al., 2020; Zotta et al., 2022). However, it was absent or outnumbered by other genera in a substantial proportion of samples, underscoring that the LAB ecology of naturally fermented olives is more complex than the dominant role assigned to this genus in much of the starter literature. *Lacticaseibacillus*, which also has documented starter and functional potential in table olives (Pino et al., 2019), was among the dominant genera particularly in alkali-treated varieties — a pattern confirmed by our differential abundance analysis — suggesting that starter development for alkali-treated olives should more systematically consider this genus alongside *Lactiplantibacillus*. Several heterofermentative LAB — particularly *Lentilactobacillus*, and occasionally *Limosilactobacillus, Levilactobacillus, Paucilactobacillus* and *Leuconostoc* — were frequently subdominant or dominant in naturally fermented olives. Their contribution to flavour development and fermentation dynamics remains poorly understood, although *Levilactobacillus* and *Leuconostoc* have been evaluated as starters in Greek olives (Chytiri et al., 2020; Vougiouklaki et al., 2024). The frequent dominance of *Lentilactobacillus* is of particular concern given its association with spoilage and biogenic amine production in protein-rich environments (O’Sullivan et al., 2015; Penland et al., 2021; Suzzi and Gardini, 2003), and warrants systematic investigation of its role across olive varieties and trade preparations. Finally, the occasional occurrence of HALAB in naturally fermented olives — where alkaline and high-salt conditions are less sustained — is an unexpected finding whose ecological and technological significance deserves further study.

The core mycobiota of table olives — dominated by *Pichia, Candida, Wickerhamomyces*, and *Saccharomyces* — was largely consistent with previous surveys (Ricciardi et al., 2025; Tsoungos et al., 2023), although their abundance was highly variable within and across varieties, and the pattern of individual *Pichia* species differed between trade preparations and varieties. Several findings extend or add to the existing literature. *Dekkera/Brettanomyces*, which is frequently reported in table olives (Medina et al., 2018; Michailidou et al., 2021; Tsoungos et al., 2023), was subdominant in several naturally fermented varieties in our dataset, a pattern that warrants attention given its spoilage potential in other fermented products (Tubia et al., 2018). *Zygotorulaspora*, whose role in the aromatic profile of table olives has been documented (Arroyo-López et al., 2016; Mougiou et al., 2023; Montaño et al., 2021) and which has been used as a starter in Nyons olives (Penland et al., 2022), was occasionally dominant in our dataset, reinforcing its relevance as a candidate for microbiome-based starter development. *Schwanniomyces* showed a strong association with Halkidiki olives, extending a previous report (Argyri et al., 2020), while *Starmerella* was consistently associated with Greek varieties (Conservolea, Kalamata and, to a lesser extent, Halkidiki) — a finding not previously reported and suggesting a variety- or origin-specific nature for this genus. *Debaryomyces*, which has been evaluated as starter cultures (Chytiri et al., 2020, Tarantini et al., 2024), was also among the genera identified in our dataset, further supporting the potential for diversifying yeast starter selection beyond the currently dominant *Wickerhamomyces* and *Saccharomyces* (Giavalisco et al., 2024). Together, these findings highlight the richness of the fungal communities of table olives and their largely untapped potential for variety-specific starter development.

The beta-diversity analysis confirmed, at a global level, the patterns already evident from the differential abundance analysis: ripeness and trade preparation were the dominant structuring factors, while olive variety within trade preparation groups and producer identity within varieties introduced additional, statistically significant layers of differentiation. The striking within-group variability visible in the ordination plots — even for samples of the same variety from the same producer — reflects the combined influence of stochastic colonization, small fermentation vessel effects, and house microbiota, as discussed above. Technical replication confirmed that this variability is biological rather than methodological, with Bray-Curtis distances between technical replicates being very low (data not shown). Our dataset included spoiled samples across several varieties, and fungal communities of spoiled samples were often visually distinct from those of unspoiled samples in the ordination plots. However, the limited number of spoiled samples and the heterogeneity of spoilage types prevented inferential analysis, and no firm conclusions on microbial markers of spoilage can be drawn from these data, although other authors have identified consistent spoilage markers for selected varieties (Arroyo-López et al., 2021; de Castro et al., 2018, 2022; Penland et al., 2021). This remains an important open question, particularly for fungal communities where dissimilatiries between spoiled and non-spoiled samples suggests that diagnostic signals may exist.

Several authors have proposed that metataxonomic analysis of microbial communities may assist in identifying the geographical origin of table olives, invoking the concept of terroir — originally developed for wine — to describe the influence of environment and raw material on microbial community composition and, ultimately, organoleptic properties (Argyri et al., 2020; Belda et al., 2017; Ferrocino et al., 2025; Kamilari et al., 2023; Kazou et al., 2020). Our data are partially consistent with this view: producers of the same variety share common microbial patterns, indicating that ecological factors associated with raw material and production environment do shape community structure. However, significant and measurable differences among producers within the same variety — confirmed by PERMANOVA — indicate that house microbiota and producer-specific practices introduce variation that cannot be attributed to terroir alone. This confounding between terroir and house microbiota effects has important implications for authentication: producers within the same PDO area can harbour more divergent microbial communities than PDO and non-PDO producers of the same variety, making microbiota-based discrimination inherently challenging. Indeed, while statistically significant differences in both bacterial and fungal communities between PDO Oliva di Gaeta and its non-PDO counterpart Itrana nera were detectable — both within the largest single producer and across all producers — the signal was weak and heavily influenced by between-firm variability. Similar results were obtained for Bella di Daunia and Nocellara del Belíce. These findings suggest that, with current amplicon-based approaches and unbalanced sampling designs, Italian PDO table olive varieties cannot be reliably discriminated from similar non-PDO varieties on the basis of microbiota composition alone. Realising the potential of microbiota-based authentication will require larger, balanced sampling designs, integration with metabolomic data (Beteinakis et al., 2024), and the application of supervised machine learning approaches (López-García et al., 2025) capable of extracting weak but consistent signals from highly variable community data. The availability of large, well annotated databases, like OliveFMBN (Ricciardi et al, 2025), in which the data of this survey have been integrated, may support efforts in this field.

The high variability in microbial community composition documented in this study, combined with the limited control that artisanal and semi-industrial producers currently exert over fermentation conditions, provides a strong rationale for the development of starter cultures tailored to specific olive varieties and trade preparations. The current starter market for table olives is dominated by a small number of commercial producers, virtually all of whom rely on *Lactiplantibacillus* as the primary — and often sole — inoculant. Examples are the Lal’Olive Crispy *Lactipl. pentosus* OM13 (Alfonzo et al., 2023a; Vege-Start 10 (Corsetti et al., 2012) and the well documented functional strain *Lactipl. pentosus* LPG1 (Benítez-Cabello et al., 2020). While the fermentative, biopreservative, and functional potential of this species in table olives is well established (Giavalisco et al., 2024; Portilha-Cunha et al., 2020; Zotta et al., 2022), our data suggest that this approach captures only a fraction of the ecologically and technologically relevant microbial diversity present in table olives. The continued reliance on a narrow range of *Lactiplantibacillus*-based starters risks progressive homogenization of the organoleptic diversity that currently characterises artisanal varieties — a diversity that is itself a reflection of the complex, variety-specific microbial communities documented here.

Microbiome-based starters offer a promising alternative (Nikoloudaki et al., 2024): rather than relying on one or a few well-characterised strains, this approach uses microbiome data to design complex starter communities that capture the diversity and ecological interactions of the native microbiota of a specific variety and production context. Our findings provide ecological foundations for this approach. For naturally fermented olives, candidate bacterial starters should look beyond *Lactiplantibacillus* to include *Lacticaseibacillus*, heterofermentative LAB such as *Levilactobacillus* and *Leuconostoc*. The role of HALAB is more controversial since some of them have been associated with spoilage (Arroyo-López et al., 2021). Fungal communities are an equally important: genera such as *Wickerhamomyces, Zygotorulaspora*, and *Debaryomyces* have documented starter or flavour potential (Benítez-Cabello et al., 2020; Giavalisco et al, 2024) and are consistent members of the mycobiota of specific varieties. The development of mixed bacterial-fungal starters grounded in variety-specific microbiome data represents a natural extension of this work, although a full characterisation of microbial interactions and their functional consequences lies beyond the scope of the present study.

Realising this potential will require not only larger and more targeted microbiome surveys, but also functional studies linking community composition to fermentation dynamics and organoleptic outcomes, and the development of competitive, variety-adapted strains capable of driving fermentation consistently without suppressing the ecological diversity that underpins the quality and identity of artisanal table olives.

## 5 Conclusions

This study presents the largest metataxonomic survey of table olive microbial communities to date, providing the first culture-independent characterisation of several Italian PDO and non-PDO varieties — most notably Oliva di Gaeta — and substantially extending existing knowledge for others, including Bella di Daunia, Ascolana del Piceno, and Nocellara del Belíce. The dominant structuring factors shaping both bacterial and fungal communities were olive ripeness and trade preparation, with the contrast between alkali-treated and naturally fermented olives representing the strongest and most consistent signal; within this framework, variety- and producer-level effects introduced additional, statistically significant differentiation consistent with the combined influence of ecological selection and house microbiota. However, the striking variability observed within the same variety and even within the same producer — driven by stochastic colonization events amplified by small-scale fermentation conditions — obscures variety-specific microbial signatures and means that, with current amplicon-based approaches, Italian PDO varieties cannot be reliably discriminated from similar non-PDO counterparts on the basis of microbiota composition alone, although integration with metabolomic data and supervised machine learning approaches may improve authentication in the future. The ecological and taxonomic complexity documented here — encompassing a diverse array of LAB, HALAB, halophilic Gram-negatives, and fungal genera with documented starter and flavour potential — provides the foundation for a new generation of variety- and preparation-specific microbiome-based starter cultures that move beyond the current reliance on *Lactiplantibacillus*-only commercial products; realising this potential will require functional studies linking community composition to fermentation dynamics and organoleptic outcomes, and the development of mixed bacterial-fungal starters grounded in variety-specific microbiome data.

## Data availability

Sequence data have been deposited in NCBI SRA with bioproject accession PRJNA1292811. The accession list of the runs used in this study is available in Supplementary data and figures. The metataxonomic data have been integrated in the OliveFMBN database (https://doi.org/10.5281/zenodo.13729346)

## Acknowledgements

The authors gratefully acknowledge the skilled technical assistance of Ms. Anita Aliano. This work was carried out within the PRIN 2022 Project METAOlive 2022NN28ZZ and received funding from the European Union Next-GenerationEU, CUP C53D23005460006 (PIANO NAZIONALE DI RIPRESA E RESILIENZA (PNRR) – MISSIONE 4 COMPONENTE 2, INVESTIMENTO 1.4 – D.D. 1048 14/07/2023).

