## Supplementary tables and figures for "A survey of bacterial and fungal communities of table olives"

**Supplementary Table 1.** The table olive samples used in this study, in detail.

| OLIVE RIPENESS | OLIVE TRADE PREPARATION | STYLE | VARIETY | N | COUNTRY |
|---|---|---|---|---|---|
| **GREEN OLIVES** | Alkali treated olives | Whole | Halkidiki | 22 | Greece |
| **GREEN OLIVES** | Alkali treated olives | Whole | Bella di Daunia | 19 | Italy |
| **GREEN OLIVES** | Alkali treated olives | Whole | Nocellara del Belice | 7 | Italy |
| **GREEN OLIVES** | Alkali treated olives | Whole | Ascolana del Piceno DOP | 6 | Italy |
| **GREEN OLIVES** | Alkali treated olives | Whole | Manzanilla | 4 | Spain |
| **GREEN OLIVES** | Alkali treated olives | Whole | Hojiblanca | 4 | Spain |
| **GREEN OLIVES** | Alkali treated olives | Whole | Plum | 2 | Greece |
| **GREEN OLIVES** | Alkali treated olives | Whole | Cacereña | 2 | Italy |
| **GREEN OLIVES** | Alkali treated olives | Pitted | Halkidiki | 1 | Italy |
| **GREEN OLIVES** | Alkali treated olives | Whole | Gordal | 1 | Spain |
| **GREEN OLIVES** | Natural olives | Whole | Itrana bianca | 15 | Italy |
| **GREEN OLIVES** | Natural olives | Whole | Tonda di Cagliari | 10 | Italy |
| **GREEN OLIVES** | Natural olives | Whole | Termite di Bitetto | 5 | Italy |
| **GREEN OLIVES** | Natural olives | Whole | Nocellara del Belíce | 5 | Italy |
| **GREEN OLIVES** | Natural olives | Whole | Pit´ze Carroga | 4 | Italy |
| **GREEN OLIVES** | Natural olives | Cracked | Ladoelia | 4 | Cyprus |
| **GREEN OLIVES** | Natural olives | Whole | Manzanilla | 3 | Cyprus, Spain |
| **GREEN OLIVES** | Natural olives | Whole | Hojiblanca | 2 | Spain |
| **GREEN OLIVES** | Natural olives | Whole | Olive verdi di Paternò | 2 | Italy |
| **GREEN OLIVES** | Natural olives | Whole | Aloreña | 2 | Spain |
| **GREEN OLIVES** | Natural olives | Whole | Arbequina | 2 | Spain |
| **GREEN OLIVES** | Natural olives | Whole | Leccino | 2 | Italy |
| **GREEN OLIVES** | Natural olives | Cracked | Manzanilla | 1 | Cyprus |

| | | | | | |
|---|---|---|---|---|---|
| **GREEN OLIVES** | Natural olives | Whole | Ascolana del Piceno DOP | 1 | Italy |
| **GREEN OLIVES** | Natural olives | Whole | Bella di Daunia | 1 | Italy |
| **GREEN OLIVES** | Natural olives | Whole | Oliva di Gaeta | 1 | Italy |
| **GREEN OLIVES** | Natural olives | Whole | Cacereña | 1 | Spain |
| **GREEN OLIVES** | Natural olives | Sliced | Aloreña | 1 | Spain |
| **GREEN OLIVES** | Natural olives | Whole | Carolea | 1 | Italy |
| **GREEN OLIVES** | Natural olives | Whole | Bella di Cerignola | 1 | Italy |
| **GREEN OLIVES** | Natural olives | Whole | Nocellara Messinese | 1 | Italy |
| **GREEN OLIVES** | Olives darkened by oxidation | Whole | Cacereña | 4 | Spain |
| **GREEN OLIVES** | Olives darkened by oxidation | Whole | Manzanilla | 1 | Spain |
| **GREEN OLIVES** | Specialties (Picholine, etc.) | Whole | Nocellara del Belice | 21 | Italy |
| **GREEN OLIVES** | Specialties (Picholine, etc.) | Cracked | Okal | 3 | Spain |
| **GREEN OLIVES** | Specialties (Picholine, etc.) | Cracked | Cornenzuelo de Jaén | 3 | Spain |
| **GREEN OLIVES** | Specialties (Picholine, etc.) | Cracked | Salella schiacciata | 3 | Italy |
| **OLIVES TURNING COLOR** | Natural olives | Whole | Caiazzana | 11 | Italy |
| **OLIVES TURNING COLOR** | Natural olives | Whole | Bella di Daunia | 8 | Italy |
| **OLIVES TURNING COLOR** | Natural olives | Whole | Leccino | 6 | Italy |
| **OLIVES TURNING COLOR** | Natural olives | Whole | Peranzana | 5 | Italy |
| **OLIVES TURNING COLOR** | Natural olives | Whole | Termite di Bitetto | 3 | Italy |
| **OLIVES TURNING COLOR** | Natural olives | Pitted | Leccino | 2 | Italy |
| **OLIVES TURNING COLOR** | Natural olives | Whole | Unknown | 1 | NA |

| | | | | | |
|---|---|---|---|---|---|
| **OLIVES TURNING COLOR** | Natural olives | Whole | Ladoelia | 1 | Cyprus |
| **OLIVES TURNING COLOR** | Natural olives | Whole | Konservolia | 1 | Italy |
| **OLIVES TURNING COLOR** | Natural olives | Whole | Taggiasca | 1 | Italy |
| **OLIVES TURNING COLOR** | Natural olives | Whole | Nocellara del Belice | 1 | Italy |
| **OLIVES TURNING COLOR** | Natural olives | Whole | Ogliarola Garganica | 1 | Italy |
| **OLIVES TURNING COLOR** | Olives darkened by oxidation | Whole | Cacereña | 4 | Spain |
| **OLIVES TURNING COLOR** | Olives darkened by oxidation | Whole | Hojiblanca | 3 | Spain |
| **OLIVES TURNING COLOR** | Olives darkened by oxidation | Whole | Manzanilla | 2 | Spain |
| **BLACK OLIVES** | Dehydrated olives | Whole | Oliva infornata di Ferrandina | 4 | Italy |
| **BLACK OLIVES** | Natural olives | Whole | Oliva di Gaeta | 65 | Italy |
| **BLACK OLIVES** | Natural olives | Whole | Itrana nera | 33 | Italy |
| **BLACK OLIVES** | Natural olives | Whole | Kalamata | 15 | Greece, Italy |
| **BLACK OLIVES** | Natural olives | Whole | Konservolia | 10 | Greece |
| **BLACK OLIVES** | Natural olives | Whole | Kalamon | 7 | Greece |
| **BLACK OLIVES** | Natural olives | Whole | Aitana | 4 | Italy |
| **BLACK OLIVES** | Natural olives | Whole | Amfissa nere | 4 | Greece |
| **BLACK OLIVES** | Natural olives | Whole | Criolla | 4 | Peru |
| **BLACK OLIVES** | Natural olives | Whole | Ladoelia | 3 | Cyprus |
| **BLACK OLIVES** | Natural olives | Whole | Picual | 2 | Italy, Egypt |
| **BLACK OLIVES** | Natural olives | Whole | Termite di Bitetto | 2 | Italy |
| **BLACK OLIVES** | Natural olives | Whole | Taggiasca | 1 | Italy |
| **BLACK OLIVES** | Natural olives | Whole | Peranzana | 1 | Italy |

| | | | | | |
|---|---|---|---|---|---|
| **BLACK OLIVES** | Natural olives | Seasoned | Leccino | 1 | Italy |

**Supplementary Table 2**. Abbreviations for olive varieties used in text and figures. Note that some varieties are produced with multiple methods (alkali treated, natural, etc.)

| OLIVE VARIETY | ABBREVIATION |
|---|---|
| **AITANA** | Alt |
| **ALOREÑA** | Alo |
| **AMFISSA BLACK** | Amf |
| **ARBEQUINA** | Arb |
| **ASCOLANA DEL PICENO PDO** | AdP |
| **BELLA DI CERIGNOLA** | Bec |
| **BELLA DI DAUNIA** | Bel |
| **CACEREÑA** | Cac |
| **CAIAZZANA** | Cai |
| **CAROLEA** | Car |
| **CORNENZUELO DE JAÉN** | Cor |
| **CRIOLLA** | Cri |
| **GORDAL** | Gor |
| **HALKIDIKI** | Hal |
| **HOJIBLANCA** | Hoj |
| **ITRANA BIANCA** | Itb |
| **ITRANA NERA** | Itn |
| **KALAMATA** | Kal |
| **KALAMON** | Klm |
| **KONSERVOLIA** | Kon |
| **LADOELIA** | Lad |
| **LECCINO** | Lec |
| **MANZANILLA** | Man |

| | |
|---|---|
| **NOCELLARA MESSINESE** | Nom |
| **NOCELLARA DEL BELICE** | Nob |
| **OGLIAROLA GARGANICA** | Ogl |
| **OKAL** | Oka |
| **OLIVA DI GAETA** | OdG |
| **OLIVA INFORNATA DI FERRANDINA** | Ofe |
| **OLIVE VERDI DI PATERNÒ** | Ovp |
| **PERANZANA** | Per |
| **PICUAL** | Pic |
| **PIT´ZE CARROGA** | Pit |
| **PLUM** | Plu |
| **SALELLA SCHIACCIATA** | Sal |
| **TAGGIASCA** | Tag |
| **TERMITE DI BITETTO** | Ter |
| **TONDA DI CAGLIARI** | TdC |
| **UNKNOWN** | Unk |

**Supplementary Table 3**. Distribution of spoiled samples among the main olive categories (combination of ripening stage and olive trade preparation).

| SPOILAGE | BLACK OLIVES, DEHYDRATED | BLACK OLIVES, NATURAL | GREEN OLIVES, ALKALI TREATED | GREEN OLIVES, DARKENED BY OXIDATION | GREEN OLIVES, NATURAL | GREEN OLIVES, SPECIALTIES (PICHOLINE, ETC.) | OLIVES TURNING COLOR, DARKENED BY OXIDATION | OLIVES TURNING COLOR, NATURAL |
|---|---|---|---|---|---|---|---|---|
| **DISCOLOURATION** | 0 | 0 | 0 | 0 | 0 | 0 | 0 | 1 |
| **FLOR** | 0 | 0 | 0 | 0 | 0 | 0 | 0 | 3 |
| **GAS** | 0 | 1 | 2 | 0 | 2 | 0 | 0 | 5 |
| **GAS POCKETS** | 0 | 0 | 2 | 0 | 0 | 0 | 0 | 0 |
| **LARVAE** | 0 | 0 | 0 | 0 | 1 | 0 | 0 | 0 |
| **OTHER** | 0 | 0 | 3 | 3 | 0 | 0 | 0 | 0 |
| **PUTRID** | 0 | 0 | 5 | 0 | 1 | 0 | 1 | 1 |
| **PUTRID/COLOR** | 0 | 0 | 1 | 0 | 0 | 0 | 0 | 0 |
| **SPOTS** | 0 | 0 | 3 | 0 | 0 | 0 | 0 | 1 |
| **ZAPATERA / PALMICHE** | 0 | 3 | 3 | 0 | 0 | 0 | 2 | 0 |
| **NONE** | 4 | 148 | 49 | 2 | 61 | 30 | 6 | 30 |

**Supplementary Table 4**. Summary statistics for the pH of brines and olives.

| RIPENING STAGE + TRADE PREPARATION | MEDIAN | MIN | MAX |
|---|---|---|---|
| **OLIVES TURNING COLOR, NATURAL** | 4.08 | 3.62 | 5.46 |
| **GREEN OLIVES, ALKALI TREATED** | 4.23 | 3.46 | 6.89 |
| **BLACK OLIVES, NATURAL** | 4.05 | 3.27 | 5.74 |
| **GREEN OLIVES, SPECIALTIES (PICHOLINE, ETC.)** | 5.01 | 3.77 | 6.63 |
| **BLACK OLIVES, DEHYDRATED** | 5.21 | 4.76 | 5.43 |
| **GREEN OLIVES, NATURAL** | 4.20 | 3.58 | 8.08 |
| **OLIVES TURNING COLOR, DARKENED BY OXIDATION** | 3.70 | 3.68 | 3.71 |
| **GREEN OLIVES, DARKENED BY OXIDATION** | 3.93 | 3.93 | 3.93 |

**Supplementary Table 5**. Summary statistics on the pH of brines and olives of olive varieties in this study. See Supplementary Table 2 for abbreviations of table varieties.

| Ripening stage + trade preparation | Olive var. short | n | Mean | Median | Min | Max | Std. dev. |
|---|---|---|---|---|---|---|---|
| Black olives, dehydrated | Ofe | 4 | 5.15 | 5.21 | 4.76 | 5.43 | 0.32 |
| Black olives, natural | Alt | 4 | 3.96 | 3.89 | 3.88 | 4.19 | 0.15 |
| Black olives, natural | Amf | 4 | 3.73 | 3.57 | 3.53 | 4.27 | 0.36 |
| Black olives, natural | Itn | 33 | 4.30 | 4.15 | 3.87 | 5.02 | 0.38 |
| Black olives, natural | Kal | 15 | 3.85 | 3.84 | 3.32 | 5.10 | 0.41 |
| Black olives, natural | Klm | 7 | 4.18 | 4.16 | 3.71 | 4.70 | 0.31 |
| Black olives, natural | Kon | 10 | 4.05 | 4.05 | 3.84 | 4.23 | 0.12 |
| Black olives, natural | Lad | 3 | 5.39 | 5.39 | 5.05 | 5.74 | 0.49 |
| Black olives, natural | Lec | 1 | 4.02 | 4.02 | 4.02 | 4.02 | - |
| Black olives, natural | OdG | 65 | 4.24 | 4.11 | 3.69 | 4.86 | 0.39 |
| Black olives, natural | Per | 1 | 4.28 | 4.28 | 4.28 | 4.28 | NA |
| Black olives, natural | Pic | 2 | 3.48 | 3.48 | 3.27 | 3.70 | 0.30 |
| Black olives, natural | Tag | 1 | 4.09 | 4.09 | 4.09 | 4.09 | - |
| Black olives, natural | Ter | 2 | 4.04 | 4.04 | 4.04 | 4.05 | 0.01 |
| Green olives, alkali treated | AdP | 6 | 4.20 | 4.13 | 4.00 | 4.56 | 0.22 |
| Green olives, alkali treated | Bel | 19 | 4.58 | 4.45 | 4.12 | 6.10 | 0.48 |
| Green olives, alkali treated | Hal | 23 | 4.05 | 3.90 | 3.74 | 5.41 | 0.39 |
| Green olives, alkali treated | Hoj | 4 | 3.84 | 3.84 | 3.84 | 3.84 | - |
| Green olives, alkali treated | Man | 4 | 3.80 | 3.80 | 3.79 | 3.82 | 0.02 |
| Green olives, alkali treated | Nob | 7 | 5.10 | 5.41 | 3.72 | 6.89 | 1.08 |
| Green olives, alkali treated | Plu | 2 | 3.59 | 3.59 | 3.46 | 3.72 | 0.18 |
| Green olives, darkened by oxidation | Man | 1 | 3.93 | 3.93 | 3.93 | 3.93 | - |

| Ripening stage + trade preparation | Olive var. short | n | Mean | Median | Min | Max | Std. dev. |
|---|---|---|---|---|---|---|---|
| Green olives, natural | AdP | 1 | 3.97 | 3.97 | 3.97 | 3.97 | - |
| Green olives, natural | Alo | 3 | 4.30 | 4.30 | 4.30 | 4.30 | - |
| Green olives, natural | Arb | 2 | 3.80 | 3.80 | 3.80 | 3.80 | - |
| Green olives, natural | Bec | 1 | 3.58 | 3.58 | 3.58 | 3.58 | - |
| Green olives, natural | Bel | 1 | 4.52 | 4.52 | 4.52 | 4.52 | - |
| Green olives, natural | Car | 1 | 4.28 | 4.28 | 4.28 | 4.28 | - |
| Green olives, natural | Hoj | 2 | 3.74 | 3.74 | 3.74 | 3.74 | - |
| Green olives, natural | Itb | 15 | 4.23 | 4.07 | 3.77 | 4.87 | 0.35 |
| Green olives, natural | Lad | 4 | 4.57 | 4.54 | 4.17 | 5.02 | 0.36 |
| Green olives, natural | Lec | 2 | 3.73 | 3.73 | 3.73 | 3.74 | 0.01 |
| Green olives, natural | Man | 4 | 5.49 | 4.37 | 4.03 | 8.08 | 2.25 |
| Green olives, natural | Nob | 5 | 4.23 | 4.30 | 3.74 | 4.76 | 0.41 |
| Green olives, natural | Nom | 1 | 3.95 | 3.95 | 3.95 | 3.95 | - |
| Green olives, natural | Ovp | 2 | 3.92 | 3.92 | 3.67 | 4.18 | 0.36 |
| Green olives, natural | Pit | 4 | 3.77 | 3.76 | 3.75 | 3.80 | 0.03 |
| Green olives, natural | TdC | 10 | 4.39 | 4.55 | 3.90 | 4.64 | 0.31 |
| Green olives, natural | Ter | 5 | 4.46 | 4.66 | 3.91 | 4.76 | 0.37 |
| Green olives, specialties (picholine, etc.) | Cor | 3 | 4.61 | 4.65 | 4.32 | 4.85 | 0.27 |
| Green olives, specialties (picholine, etc.) | Nob | 21 | 5.61 | 5.79 | 3.77 | 6.63 | 0.88 |
| Green olives, specialties (picholine, etc.) | Oka | 3 | 4.50 | 4.45 | 4.45 | 4.59 | 0.08 |
| Green olives, specialties (picholine, etc.) | Sal | 3 | 4.77 | 4.78 | 4.46 | 5.07 | 0.31 |

| Ripening stage + trade preparation | Olive var. short | n | Mean | Median | Min | Max | Std. dev. |
|---|---|---|---|---|---|---|---|
| Olives turning color, darkened by oxidation | Hoj | 3 | 3.68 | 3.68 | 3.68 | 3.68 | - |
| Olives turning color, darkened by oxidation | Man | 2 | 3.71 | 3.71 | 3.71 | 3.71 | - |
| Olives turning color, natural | Bel | 8 | 4.19 | 4.02 | 3.82 | 5.46 | 0.53 |
| Olives turning color, natural | Cai | 11 | 4.28 | 4.22 | 3.81 | 4.92 | 0.45 |
| Olives turning color, natural | Kon | 1 | 3.77 | 3.77 | 3.77 | 3.77 | - |
| Olives turning color, natural | Lad | 1 | 4.10 | 4.10 | 4.10 | 4.10 | - |
| Olives turning color, natural | Lec | 8 | 4.12 | 4.14 | 3.71 | 4.44 | 0.24 |
| Olives turning color, natural | Nob | 1 | 3.62 | 3.62 | 3.62 | 3.62 | - |
| Olives turning color, natural | Ogl | 1 | 4.36 | 4.36 | 4.36 | 4.36 | - |
| Olives turning color, natural | Per | 5 | 4.64 | 4.66 | 4.49 | 4.77 | 0.14 |
| Olives turning color, natural | Tag | 1 | 3.62 | 3.62 | 3.62 | 3.62 | - |
| Olives turning color, natural | Ter | 3 | 4.17 | 4.11 | 4.03 | 4.38 | 0.18 |
| Olives turning color, natural | Unk | 1 | 3.96 | 3.96 | 3.96 | 3.96 | - |

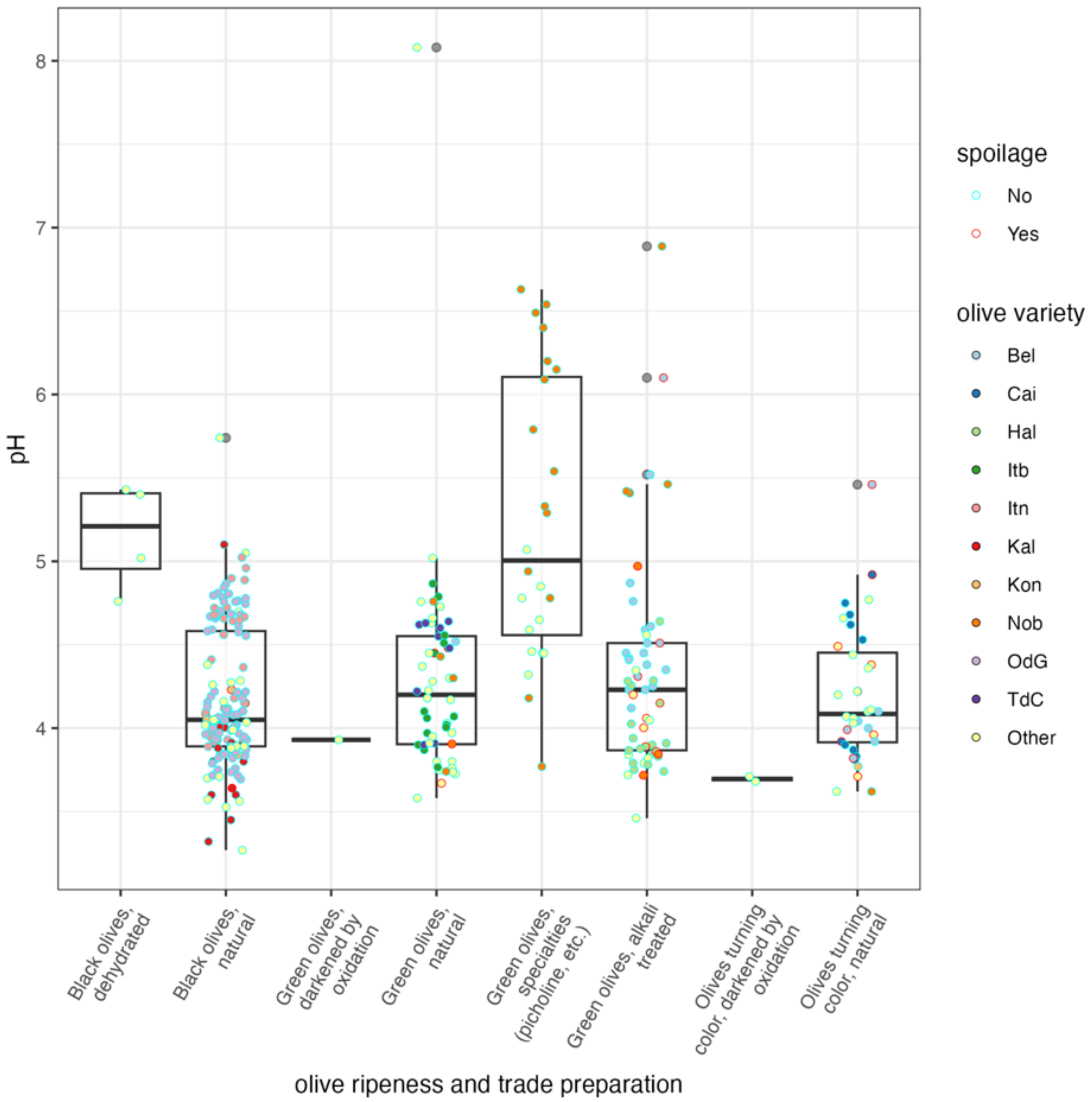


**Supplementary Figure 1**. Distribution of pH of brines (all samples except dehydrated olives, where pH was measured after homogenizing in water) for different combinations of olive ripeness and trade preparation. Different colors are used to distinguish the main varieties (those with at least 10 samples) from the others (which are pooled in the group "Other"). For olive variety abbreviations see Supplementary Table 2. The border of the symbols indicates if the samples were spoiled or not (regardless of the spoilage type).

**Supplementary table 6**. Summary statistics for the titratable acidity of brines (g/L of brine).

| Ripening stage + trade preparation | median | min | max |
|---|---|---|---|
| Olives turning color, natural | 4.59 | 2.19 | 16.26 |
| Green olives, alkali treated | 5.19 | 1.01 | 22.87 |
| Black olives, natural | 8.56 | 0.56 | 32.74 |
| Green olives, specialties (picholine, etc.) | 1.14 | 0.27 | 9.59 |
| Green olives, natural | 6.15 | 1.23 | 55.77 |
| Olives turning color, darkened by oxidation | 29.48 | 26.71 | 32.25 |
| Green olives, darkened by oxidation | 15.17 | 15.17 | 15.17 |

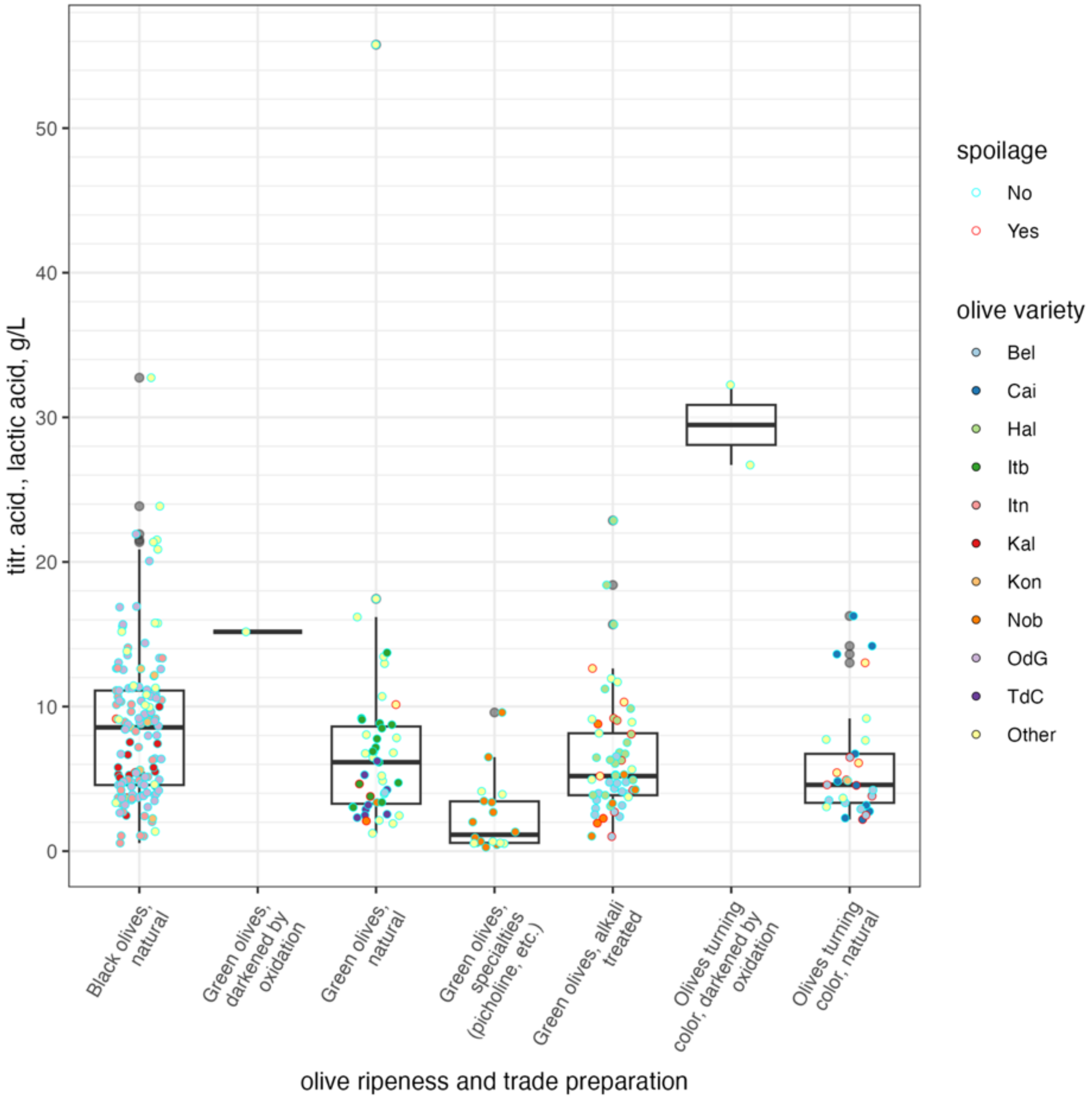


**Supplementary Figure 2**. Distribution of titratable acidity of brines (all samples except dehydrated olives, g/L lactic acid) for different combinations of olive ripeness and trade preparation. Different colors are used to distinguish the main varieties (those with at least 10 samples) from the others (which are pooled in the group "Other"). For olive variety abbreviations see Supplementary Table 2. The border of the symbols indicates if the samples were spoiled or not (regardless of the spoilage type).

**Supplementary Table 7**. Summary statistics for the salt content of brines and olives (g/L of brine or g/kg of olives for dried olives).

| Ripening stage + trade preparation | median | min | max |
|---|---|---|---|
| Black olives, dehydrated | 38.22 | 12.50 | 65.48 |
| Black olives, natural | 81.97 | 9.79 | 169.71 |
| Green olives, alkali treated | 78.14 | 34.56 | 157.56 |
| Green olives, darkened by oxidation | 33.23 | 33.23 | 33.23 |
| Green olives, natural | 54.61 | 15.51 | 246.77 |
| Green olives, specialties (picholine, etc.) | 32.39 | 2.15 | 99.32 |
| Olives turning color, darkened by oxidation | 22.98 | 17.64 | 28.33 |
| Olives turning color, natural | 59.38 | 37.41 | 147.92 |

**Supplementary Table 8.** Summary statistics on the NaCl content of brines and olives of olive varieties in this study (g/L of brine or g/kg of olives for dried olives). See supplementary Table 2 for abbreviations of table varieties.

| Ripening stage + trade preparation | Olive variety | n | mean | median | min | max | sd |
|---|---|---|---|---|---|---|---|
| Black olives, dehydrated | Ofe | 4 | 38.6 | 38.2 | 12.5 | 65.5 | 23.0 |
| Black olives, natural | Alt | 4 | 79.0 | 79.4 | 69.0 | 88.3 | 8.3 |
| Black olives, natural | Amf | 4 | 111.0 | 114.7 | 84.2 | 130.7 | 19.4 |
| Black olives, natural | Itn | 33 | 52.3 | 41.6 | 9.8 | 113.8 | 29.5 |
| Black olives, natural | Kal | 15 | 99.2 | 101.8 | 32.6 | 169.7 | 29.5 |
| Black olives, natural | Klm | 7 | 77.9 | 82.9 | 52.8 | 97.1 | 15.2 |
| Black olives, natural | Kon | 10 | 109.8 | 114.5 | 57.5 | 152.5 | 26.3 |
| Black olives, natural | Lad | 3 | 15.5 | 15.5 | 15.1 | 15.9 | 0.5 |
| Black olives, natural | Lec | 1 | 49.4 | 49.4 | 49.4 | 49.4 | - |
| Black olives, natural | OdG | 65 | 79.5 | 81.7 | 29.9 | 116.4 | 19.9 |
| Black olives, natural | Per | 1 | 124.6 | 124.6 | 124.6 | 124.6 | - |
| Black olives, natural | Pic | 2 | 109.9 | 109.9 | 94.2 | 125.7 | 22.3 |
| Black olives, natural | Tag | 1 | 57.6 | 57.6 | 57.6 | 57.6 | - |
| Black olives, natural | Ter | 2 | 83.7 | 83.7 | 77.0 | 90.4 | 9.5 |
| Green olives, alkali treated | AdP | 6 | 79.0 | 78.6 | 65.4 | 94.2 | 9.6 |
| Green olives, alkali treated | Bel | 19 | 72.9 | 68.1 | 34.6 | 129.5 | 22.2 |
| Green olives, alkali treated | Hal | 23 | 86.0 | 77.5 | 43.4 | 157.6 | 28.4 |
| Green olives, alkali treated | Hoj | 4 | 83.8 | 83.8 | 83.8 | 83.8 | - |
| Green olives, alkali treated | Man | 4 | 73.2 | 73.2 | 55.8 | 90.6 | 24.6 |
| Green olives, alkali treated | Nob | 7 | 83.4 | 89.3 | 45.9 | 104.0 | 20.5 |
| Green olives, alkali treated | Plu | 2 | 88.3 | 88.3 | 79.0 | 97.6 | 13.1 |

| Ripening stage + trade preparation | Olive variety | n | mean | median | min | max | sd |
|---|---|---|---|---|---|---|---|
| Green olives, darkened by oxidation | Man | 1 | 33.2 | 33.2 | 33.2 | 33.2 | - |
| Green olives, natural | AdP | 1 | 87.8 | 87.8 | 87.8 | 87.8 | - |
| Green olives, natural | Alo | 3 | 42.3 | 42.3 | 42.3 | 42.3 | - |
| Green olives, natural | Arb | 2 | 58.4 | 58.4 | 58.4 | 58.4 | - |
| Green olives, natural | Bec | 1 | 54.1 | 54.1 | 54.1 | 54.1 | - |
| Green olives, natural | Bel | 1 | 110.0 | 110.0 | 110.0 | 110.0 | - |
| Green olives, natural | Car | 1 | 38.9 | 38.9 | 38.9 | 38.9 | - |
| Green olives, natural | Hoj | 2 | 74.8 | 74.8 | 74.8 | 74.8 | - |
| Green olives, natural | Itb | 15 | 53.5 | 47.2 | 30.3 | 101.2 | 24.4 |
| Green olives, natural | Lad | 4 | 73.8 | 16.5 | 15.5 | 246.8 | 115.3 |
| Green olives, natural | Lec | 2 | 118.3 | 118.3 | 116.7 | 119.9 | 2.2 |
| Green olives, natural | Man | 4 | 53.4 | 64.7 | 16.2 | 79.3 | 33.0 |
| Green olives, natural | Nob | 5 | 70.9 | 67.0 | 42.7 | 106.5 | 26.8 |
| Green olives, natural | Nom | 1 | 52.0 | 52.0 | 52.0 | 52.0 | - |
| Green olives, natural | Ovp | 2 | 53.1 | 53.1 | 41.9 | 64.2 | 15.7 |
| Green olives, natural | Pit | 4 | 36.1 | 35.0 | 34.8 | 38.4 | 2.0 |
| Green olives, natural | TdC | 10 | 58.8 | 61.7 | 43.7 | 77.9 | 11.0 |
| Green olives, natural | Ter | 5 | 94.6 | 92.0 | 52.2 | 123.5 | 29.7 |
| Green olives, specialties (picholine, etc.) | Cor | 3 | 25.2 | 31.3 | 2.1 | 42.1 | 20.7 |
| Green olives, specialties (picholine, etc.) | Nob | 21 | 40.8 | 34.6 | 18.9 | 99.3 | 21.3 |
| Green olives, specialties (picholine, etc.) | Oka | 3 | 31.7 | 40.5 | 6.1 | 48.5 | 22.5 |

| Ripening stage + trade preparation | Olive variety | n | mean | median | min | max | sd |
|---|---|---|---|---|---|---|---|
| Green olives, specialties (picholine, etc.) | Sal | 3 | 16.4 | 19.1 | 9.6 | 20.4 | 5.9 |
| Olives turning color, darkened by oxidation | Hoj | 3 | 17.6 | 17.6 | 17.6 | 17.6 | - |
| Olives turning color, darkened by oxidation | Man | 2 | 28.3 | 28.3 | 28.3 | 28.3 | - |
| Olives turning color, natural | Bel | 8 | 79.7 | 78.6 | 49.1 | 108.5 | 23.2 |
| Olives turning color, natural | Cai | 11 | 48.2 | 45.8 | 39.2 | 62.7 | 7.6 |
| Olives turning color, natural | Kon | 1 | 48.5 | 48.5 | 48.5 | 48.5 | - |
| Olives turning color, natural | Lad | 1 | 55.0 | 55.0 | 55.0 | 55.0 | - |
| Olives turning color, natural | Lec | 8 | 64.8 | 68.0 | 37.4 | 82.3 | 15.1 |
| Olives turning color, natural | Nob | 1 | 53.8 | 53.8 | 53.8 | 53.8 | - |
| Olives turning color, natural | Ogl | 1 | 75.8 | 75.8 | 75.8 | 75.8 | - |
| Olives turning color, natural | Per | 5 | 89.5 | 77.8 | 72.3 | 118.4 | 25.2 |
| Olives turning color, natural | Tag | 1 | 147.9 | 147.9 | 147.9 | 147.9 | - |
| Olives turning color, natural | Ter | 3 | 57.7 | 50.3 | 46.7 | 76.2 | 16.1 |
| Olives turning color, natural | Unk | 1 | 55.7 | 55.7 | 55.7 | 55.7 | - |

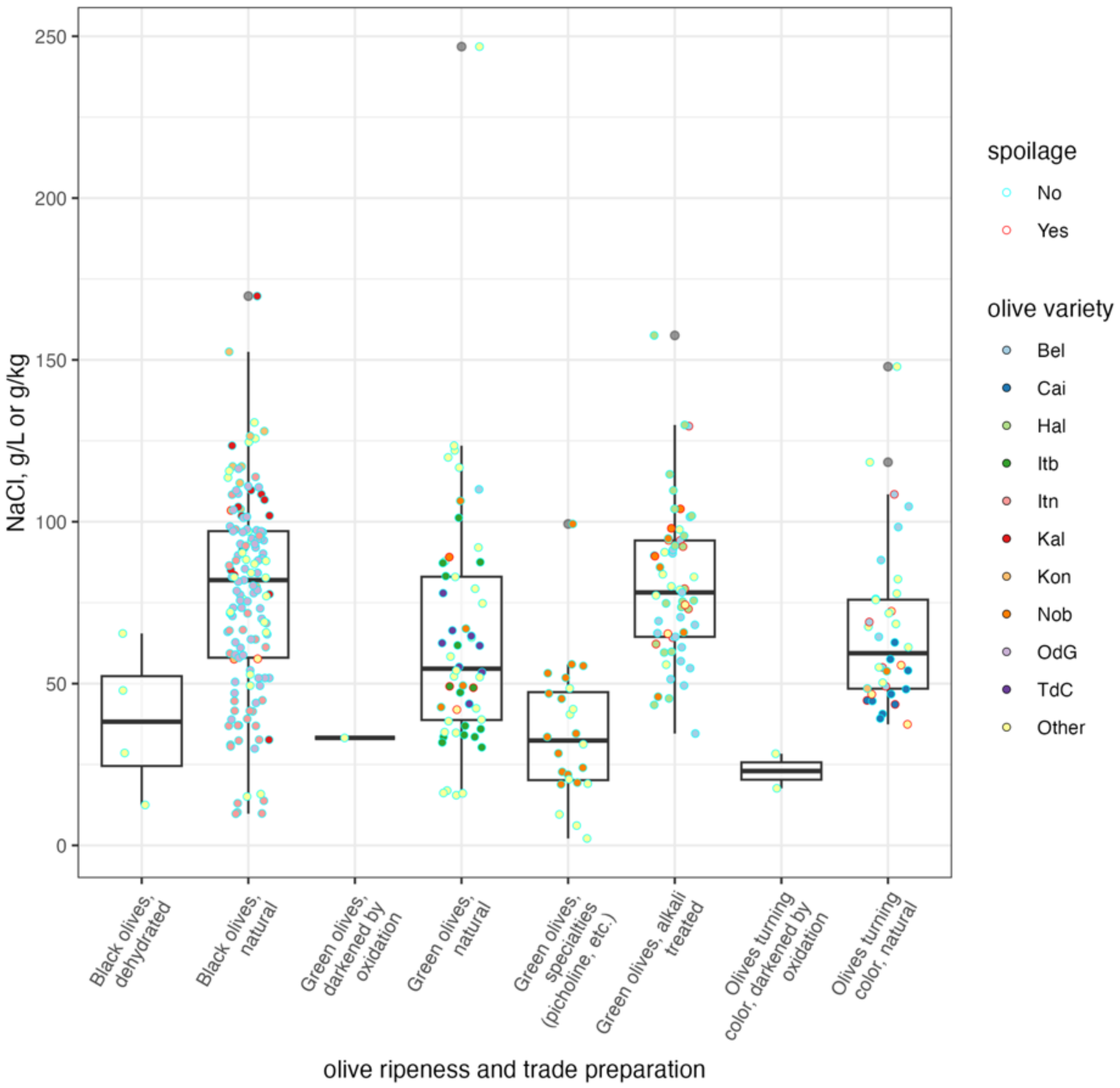


**Supplementary Figure 3**. Distribution of salt content of brines and olives (g/L or g/kg NaCl) for different combinations of olive ripeness and trade preparation. Different colors are used to distinguish the main varieties (those with at least 10 samples) from the others (which are pooled in the group "Other"). For olive variety abbreviations see Supplementary Table 2. The border of the symbols indicates if the samples were spoiled or not (regardless of the spoilage type).

**Supplementary table 9**. Summary data on the identification of isolates obtained from enumeration plates.

| Count of Identif. MALDI-ToF | | |
|---|---|---|
| substrate_from | Identif. MALDI-ToF | Total |
| Pseudomonas Selective Agar Base (PSAB) | *Aerococcus viridans* | 1 |
| | *Candida boidinii* | 8 |
| | *Enterococcus casseliflavus* | 1 |
| | no identification available | 4 |
| | *Pichia manshurica* | 2 |
| | *P. membranifaciens* | 2 |
| | *Psychrob. namhaensis* | 1 |
| | *Stenot. rhizophila* | 1 |
| | *Wickerhamomyces anomalus* | 4 |
| PSA total | | 24 |
| Glucose Yeast Extract Agar with chloramphenicol (GYEAC) | *Brettanomyces anomalus* | 1 |
| | *Candida boidinii* | 14 |
| | *C. parapsilosis* | 1 |
| | *Debaryomyces hansenii* | 2 |
| | *Hyphopichia burtonii* | 1 |
| | *Kluyveromyces lactis* | 6 |
| | *Meyerozyma guillierimondii* | 1 |
| | no identification available | 35 |
| | *P. kudriavzevii* | 11 |
| | *P. manshurica* | 23 |
| | *P. membranifaciens* | 44 |
| | *Rhodotorula mucilaginosa* | 1 |
| | *Saccharomyces cerevisiae* | 21 |
| | *Schwanniomyces etchellsii* | 1 |
| | *Wickerhamiella pararugosa* | 1 |
| | *W. anomalus* | 19 |
| GYEAC Total | | 182 |
| Modified MRS agar with sodium azide (mMRS+NA) | *Lactiplantibacillus pentosus* | 227 |
| | *Lactipl.plantarum* | 6 |
| | *Lentilactobacillus buchneri* | 2 |
| | *Leuconostoc miyukkimchii* | 1 |
| | *Loigol.coryniformis* | 1 |
| | no identification available | 21 |
| | *Secundilactobacillus collinoides* | 2 |
| | *Weissella cibaria* | 1 |
| mMRS+NA Total | | 261 |
| | *Aeroc. viridans* | 1 |
| | *C. boidinii* | 1 |

| | | |
|---|---|---|
| Plate Count Agar Standard with 5% NaCl (PCA + NaCl) | *D. hansenii* | 1 |
| | *Enteroc. casseliflavus* | 1 |
| | *Geotrichum candidum* | 1 |
| | *Kocuria palustris* | 1 |
| | *Lactipl. pentosus* | 19 |
| | *Meyer. guillierimondii* | 1 |
| | no identification available | 24 |
| | *Peribacillus simplex* | 1 |
| | *P. manshurica* | 4 |
| | *P. membranifaciens* | 2 |
| | *Staphylococcus aureus* | 1 |
| | *Staph. hominis* | 1 |
| | *Staph. nepalensis* | 1 |
| | *Staph. warneri* | 3 |
| | *W. anomalus* | 1 |
| PCA + NaCl Total | | 64 |
| Grand Total | | 531 |

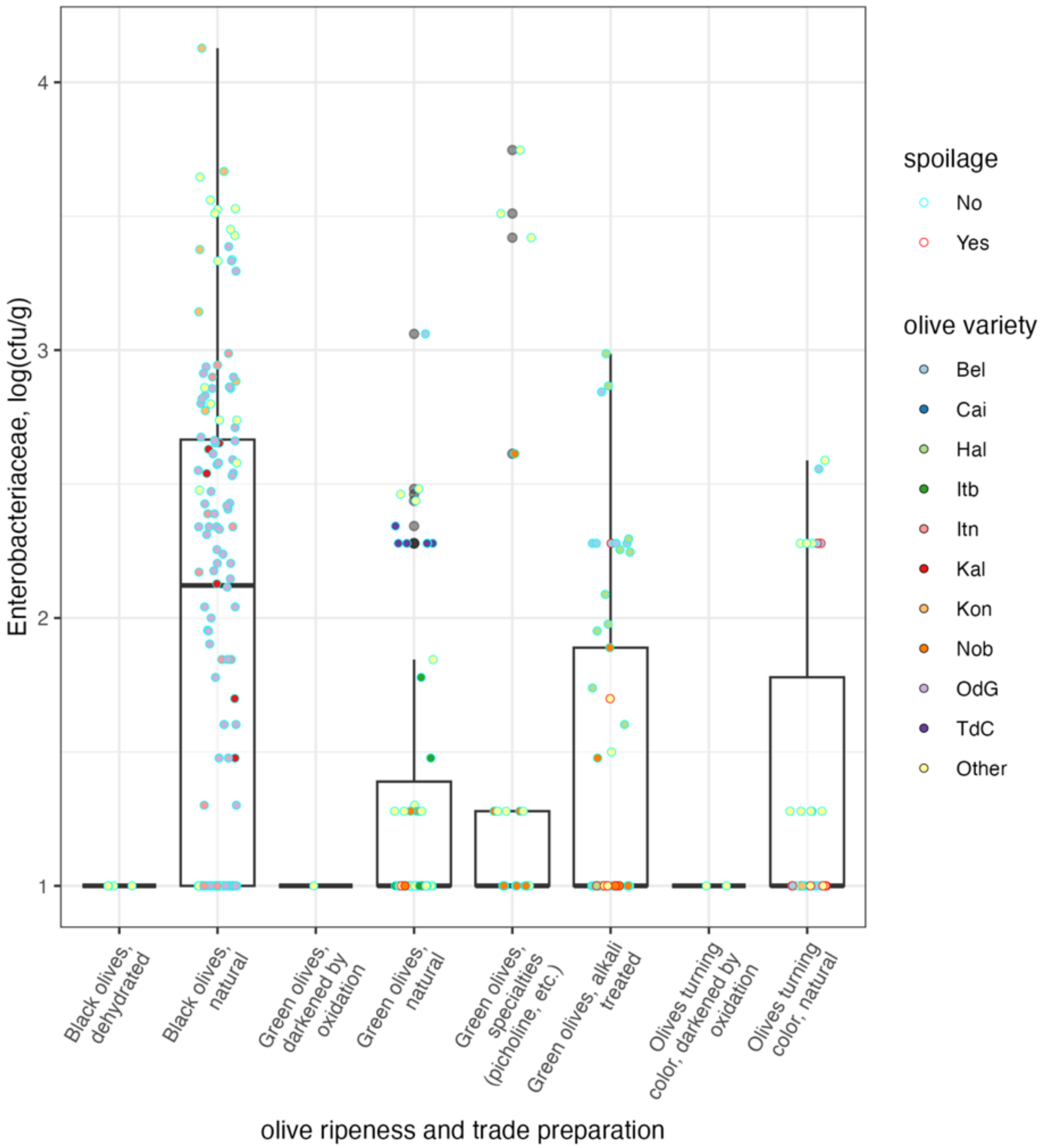


**Supplementary Figure 4**. Distribution of viable counts for *Enterobacteriaceae* for different combinations of olive ripeness and trade preparation. Different colors are used to distinguish the main varieties (those with at least 10 samples) from the others (which are pooled in the group "Other"). For olive variety abbreviations see Supplementary Table 2. The border of the symbols indicates if the samples were spoiled or not (regardless of the spoilage type).

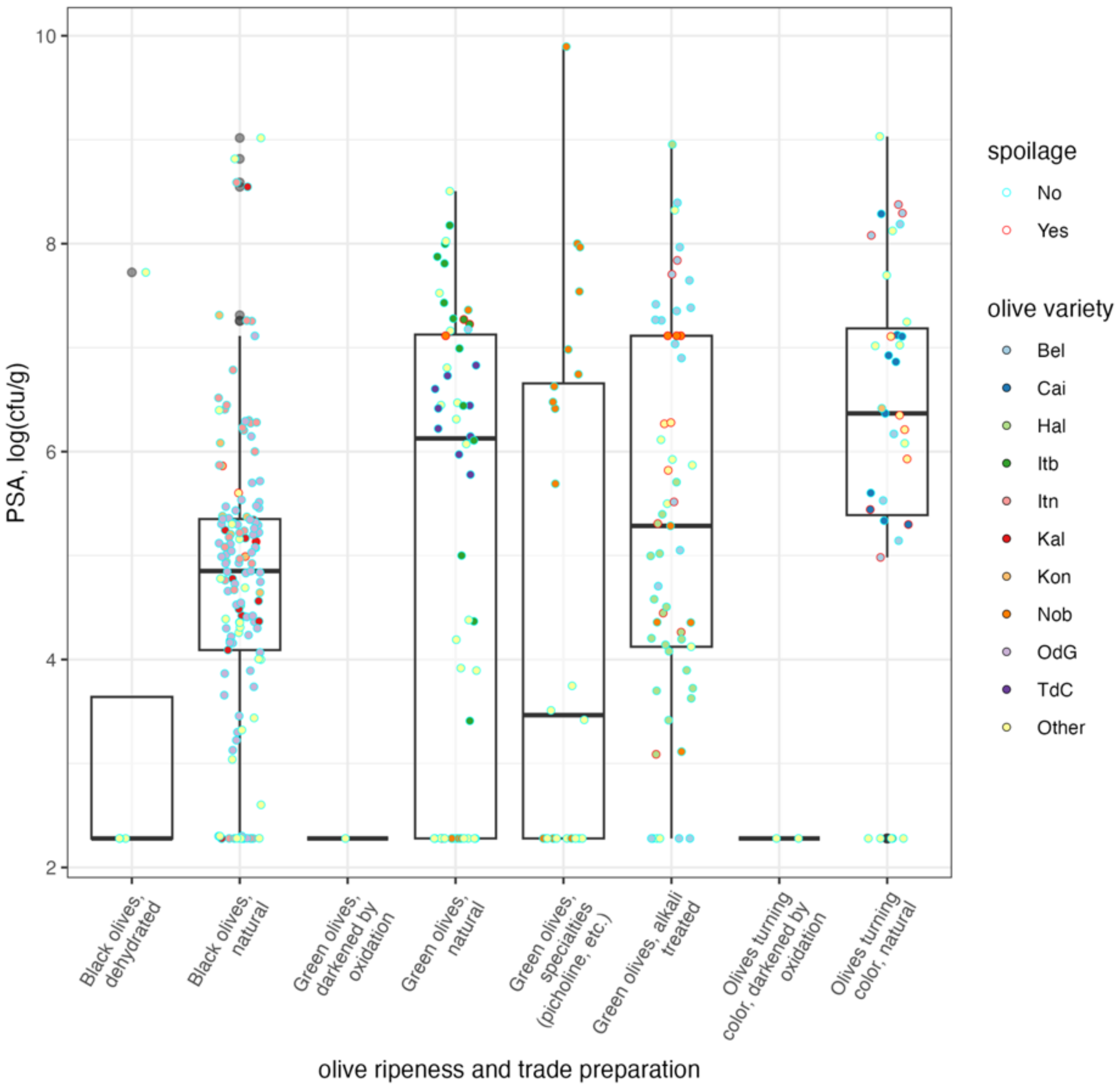


**Supplementary Figure 5**. Distribution of viable counts for contaminants on Pseudomonas Agar Base for different combinations of olive ripeness and trade preparation. Different colors are used to distinguish the main varieties (those with at least 10 samples) from the others (which are pooled in the group “Other”). For olive variety abbreviations see Supplementary Table 2. The border of the symbols indicates if the samples were spoiled or not (regardless of the spoilage type).

**Supplementary Table 10**. Descriptive statistics for viable counts of lactic acid bacteria, log(cfu/g), on mMRS+NA. na means “not available” as counts were below the detection limit of our method 2.23 log(cfu/g or ml)

| ripen. + trade prep. | median | min | max |
|---|---|---|---|
| Black olives, dehydrated | na | na | na |
| Black olives, natural | 4.82 | 2. na | 8.32 |
| Green olives, alkali treated | 5.15 | 2. na | 8.88 |
| Green olives, darkened by oxidation | 8.02 | 8.02 | 8.02 |
| Green olives, natural | 5.35 | 2. na | 8.99 |
| Green olives, specialties (picholine, etc.) | 6.20 | na | 9.02 |
| Olives turning color, darkened by oxidation | na | na | na |
| Olives turning color, natural | 6.14 | na | 9.01 |

**Supplementary Table 11.** Summary statistics on the LAB counts (log(cfu/g) brines and olives of olive varieties in this study. See supplementary Table 2 for abbreviations of table varieties. na means “not available” as counts were below the detection limit of our method 2.23 log(cfu/g or ml)

| Ripening stage + trade prep. | Olive variety | n | mean | median | min | max | sd |
|---|---|---|---|---|---|---|---|
| Black olives, dehydrated | Ofe | 4 | na | na | na | na | 0.0 |
| Black olives, natural | Alt | 4 | 4.9 | 5.1 | 3.0 | 6.3 | 1.6 |
| Black olives, natural | Amf | 4 | 4.7 | 5.5 | na | 5.6 | 1.6 |
| Black olives, natural | Itn | 33 | 5.8 | 7.2 | na | 7.8 | 2.0 |
| Black olives, natural | Kal | 15 | 5.5 | 6.0 | na | 7.5 | 1.6 |
| Black olives, natural | Klm | 7 | 4.3 | 4.4 | na | 5.4 | 1.0 |
| Black olives, natural | Kon | 10 | 6.2 | 5.8 | 5.0 | 8.3 | 1.2 |
| Black olives, natural | Lad | 3 | 4.7 | 4.7 | na | 7.1 | 3.4 |
| Black olives, natural | Lec | 1 | na | na | na | na | na |
| Black olives, natural | OdG | 65 | 3.6 | 3.4 | na | 6.1 | 1.4 |
| Black olives, natural | Per | 1 | 4.8 | 4.8 | 4.8 | 4.8 | - |
| Black olives, natural | Pic | 2 | na | na | na | na | 0.0 |
| Black olives, natural | Tag | 1 | 8.3 | 8.3 | 8.3 | 8.3 | - |
| Black olives, natural | Ter | 2 | 6.1 | 6.1 | 6.0 | 6.2 | 0.1 |
| Green olives, alkali treated | AdP | 6 | 5.7 | 5.4 | 4.7 | 8.2 | 1.3 |
| Green olives, alkali treated | Bel | 19 | 7.6 | 8.3 | na | 8.8 | 1.5 |
| Green olives, alkali treated | Hal | 23 | 4.6 | 4.5 | na | 8.9 | 1.2 |
| Green olives, alkali treated | Hoj | 4 | 8.2 | 8.2 | 8.2 | 8.2 | - |
| Green olives, alkali treated | Man | 4 | 4.6 | 4.6 | na | 7.0 | 3.3 |
| Green olives, alkali treated | Nob | 7 | 3.9 | 4.4 | na | 6.4 | 1.7 |
| Green olives, alkali treated | Plu | 2 | 4.8 | 4.8 | na | 7.2 | 3.5 |

| Ripening stage + trade prep. | Olive variety | n | mean | median | min | max | sd |
|---|---|---|---|---|---|---|---|
| Green olives, darkened by oxidation | Man | 1 | 8.0 | 8.0 | 8.0 | 8.0 | - |
| Green olives, natural | AdP | 1 | 5.5 | 5.5 | 5.5 | 5.5 | - |
| Green olives, natural | Alo | 3 | na | na | na | na | - |
| Green olives, natural | Arb | 2 | na | na | na | na | - |
| Green olives, natural | Bec | 1 | 7.4 | 7.4 | 7.4 | 7.4 | - |
| Green olives, natural | Bel | 1 | 6.3 | 6.3 | 6.3 | 6.3 | - |
| Green olives, natural | Car | 1 | 7.9 | 7.9 | 7.9 | 7.9 | - |
| Green olives, natural | Hoj | 2 | 6.1 | 6.1 | 6.1 | 6.1 | - |
| Green olives, natural | Itb | 15 | 6.3 | 7.4 | na | 9.0 | 2.5 |
| Green olives, natural | Lad | 4 | na | na | na | na | 0.0 |
| Green olives, natural | Lec | 2 | 4.8 | 4.8 | 4.8 | 4.8 | 0.0 |
| Green olives, natural | Man | 4 | 3.7 | na | na | 6.6 | 2.5 |
| Green olives, natural | Nob | 5 | 4.5 | 5.6 | na | 6.5 | 2.0 |
| Green olives, natural | Nom | 1 | 7.0 | 7.0 | 7.0 | 7.0 | - |
| Green olives, natural | Ovp | 2 | na | na | na | na | 0.0 |
| Green olives, natural | Pit | 4 | 8.5 | 8.5 | 8.2 | 8.7 | 0.3 |
| Green olives, natural | TdC | 10 | 4.2 | na | na | 8.9 | 2.9 |
| Green olives, natural | Ter | 5 | 4.4 | 4.5 | na | 5.7 | 1.3 |
| Green olives, specialties (picholine, etc.) | Cor | 3 | 2.7 | na | na | 3.4 | 0.7 |
| Green olives, specialties (picholine, etc.) | Nob | 21 | 6.5 | 7.2 | na | 9.5 | 2.5 |
| Green olives, specialties (picholine, etc.) | Oka | 3 | 3.2 | 3.5 | na | 3.7 | 0.8 |
| Green olives, specialties (picholine, etc.) | Sal | 3 | 6.1 | 6.2 | 5.3 | 6.9 | 0.8 |

| Ripening stage + trade prep. | Olive variety | n | mean | median | min | max | sd |
|---|---|---|---|---|---|---|---|
| Olives turning color, darkened by oxidation | Hoj | 3 | na | na | na | na | - |
| Olives turning color, darkened by oxidation | Man | 2 | na | na | na | na | - |
| Olives turning color, natural | Bel | 8 | 7.1 | 7.8 | na | 8.8 | 2.1 |
| Olives turning color, natural | Cai | 11 | 3.2 | na | na | 7.6 | 1.9 |
| Olives turning color, natural | Kon | 1 | na | na | na | na | - |
| Olives turning color, natural | Lad | 1 | na | na | na | na | - |
| Olives turning color, natural | Lec | 8 | 7.1 | 7.9 | na | 9.3 | 2.5 |
| Olives turning color, natural | Nob | 1 | 7.3 | 7.3 | 7.3 | 7.3 | - |
| Olives turning color, natural | Ogl | 1 | 7.9 | 7.9 | 7.9 | 7.9 | - |
| Olives turning color, natural | Per | 5 | 4.4 | na | na | 8.7 | 3.7 |
| Olives turning color, natural | Ter | 3 | 5.8 | 6.1 | na | 9.1 | 3.4 |
| Olives turning color, natural | Unk | 1 | 8.7 | 8.7 | 8.7 | 8.7 | - |

**Supplementary Table 12**. Descriptive statistics for counts of yeasts and molds, log(cfu/g) on Glucose Yeast Extract Chloramphenicol Agar. na means "not available" as counts were below the detection limit of our method, 2.23 log(cfu/g or ml)

| Ripening stage + trade prep. | median | min | max |
|---|---|---|---|
| Olives turning color, natural | 6.85 | na | 8.90 |
| Green olives, alkali treated | 5.02 | na | 8.17 |
| Black olives, natural | 5.41 | na | 9.26 |
| Green olives, specialties (picholine, etc.) | 5.46 | na | 8.18 |
| Black olives, dehydrated | 5.82 | na | 7.30 |
| Green olives, natural | 6.45 | na | 9.01 |
| Olives turning color, darkened by oxidation | 5.26 | 5.12 | 5.40 |
| Green olives, darkened by oxidation | 6.62 | 6.62 | 6.62 |

**Supplementary Table 13**. Summary statistics on the yeasts and molds counts (log(cfu/g) in brines and olives for olive varieties in this study. See Supplementary Table 2 for abbreviations of table varieties. na means “not available” as counts were below the detection limit of our method ,2.23 log(cfu/g or ml)

| Ripening stage + trade preparation | Olive variety | n | mean | median | min | max | sd |
|---|---|---|---|---|---|---|---|
| Black olives, dehydrated | Ofe | 4 | 5.3 | 5.8 | na | 7.3 | 2.2 |
| Black olives, natural | Alt | 4 | 4.5 | 4.5 | 3.5 | 5.7 | 1.1 |
| Black olives, natural | Amf | 4 | 4.8 | 4.9 | 4.4 | 5.2 | 0.3 |
| Black olives, natural | Itn | 33 | 6.5 | 6.7 | 4.4 | 9.3 | 1.1 |
| Black olives, natural | Kal | 15 | 5.0 | 4.8 | 3.5 | 8.5 | 1.2 |
| Black olives, natural | Klm | 7 | 4.5 | 4.4 | 3.7 | 5.6 | 0.6 |
| Black olives, natural | Kon | 10 | 5.3 | 5.1 | 4.7 | 6.4 | 0.6 |
| Black olives, natural | Lad | 3 | 8.9 | 8.9 | 8.8 | 9.0 | 0.1 |
| Black olives, natural | Lec | 1 | na | na | na | na | - |
| Black olives, natural | OdG | 65 | 5.2 | 5.4 | 3.0 | 6.2 | 0.7 |
| Black olives, natural | Per | 1 | 5.1 | 5.1 | 5.1 | 5.1 | - |
| Black olives, natural | Pic | 2 | 7.8 | 7.8 | 7.4 | 8.1 | 0.5 |
| Black olives, natural | Tag | 1 | 6.3 | 6.3 | 6.3 | 6.3 | - |
| Black olives, natural | Ter | 2 | na | na | na | na | 0.0 |
| Green olives, alkali treated | AdP | 6 | 6.1 | 6.0 | 5.1 | 7.2 | 0.7 |
| Green olives, alkali treated | Bel | 19 | 5.7 | 6.2 | na | 8.2 | 2.0 |
| Green olives, alkali treated | Hal | 23 | 4.5 | 4.3 | 3.4 | 5.7 | 0.6 |
| Green olives, alkali treated | Hoj | 4 | 4.0 | 4.0 | 4.0 | 4.0 | - |
| Green olives, alkali treated | Man | 4 | 5.7 | 5.7 | 4.5 | 6.9 | 1.7 |
| Green olives, alkali treated | Nob | 7 | 5.4 | 5.0 | na | 7.4 | 2.0 |
| Green olives, alkali treated | Plu | 2 | 5.0 | 5.0 | 4.4 | 5.5 | 0.8 |

| Ripening stage + trade preparation | Olive variety | n | mean | median | min | max | sd |
|---|---|---|---|---|---|---|---|
| Green olives, darkened by oxidation | Man | 1 | 6.6 | 6.6 | 6.6 | 6.6 | - |
| Green olives, natural | AdP | 1 | 6.2 | 6.2 | 6.2 | 6.2 | - |
| Green olives, natural | Alo | 3 | 7.7 | 7.7 | 7.7 | 7.7 | - |
| Green olives, natural | Arb | 2 | 6.8 | 6.8 | 6.8 | 6.8 | - |
| Green olives, natural | Bec | 1 | 7.4 | 7.4 | 7.4 | 7.4 | - |
| Green olives, natural | Bel | 1 | 5.5 | 5.5 | 5.5 | 5.5 | - |
| Green olives, natural | Car | 1 | 7.9 | 7.9 | 7.9 | 7.9 | - |
| Green olives, natural | Hoj | 2 | 5.3 | 5.3 | 5.3 | 5.3 | - |
| Green olives, natural | Itb | 15 | 7.0 | 8.0 | na | 9.0 | 2.1 |
| Green olives, natural | Lad | 4 | 3.3 | na | na | 6.4 | 2.1 |
| Green olives, natural | Lec | 2 | 2.9 | 2.9 | na | 3.6 | 0.9 |
| Green olives, natural | Man | 4 | 5.2 | 5.8 | na | 7.5 | 2.7 |
| Green olives, natural | Nob | 5 | 4.6 | 4.4 | na | 7.4 | na |
| Green olives, natural | Nom | 1 | 7.0 | 7.0 | 7.0 | 7.0 | - |
| Green olives, natural | Ovp | 2 | 5.3 | 5.3 | 4.4 | 6.2 | 1.3 |
| Green olives, natural | Pit | 4 | 3.9 | na | na | 7.1 | 2.8 |
| Green olives, natural | TdC | 10 | 7.1 | 7.1 | 6.5 | 8.0 | 0.4 |
| Green olives, natural | Ter | 5 | 3.9 | 4.0 | na | 5.3 | 1.1 |
| Green olives, specialties (picholine, etc.) | Cor | 3 | 3.8 | 4.4 | na | 4.7 | 1.3 |
| Green olives, specialties (picholine, etc.) | Nob | 21 | 5.5 | 6.2 | na | 8.2 | 2.2 |
| Green olives, specialties (picholine, etc.) | Oka | 3 | 4.8 | 4.8 | 4.4 | 5.3 | 0.5 |
| Green olives, specialties (picholine, etc.) | Sal | 3 | 6.1 | 6.2 | 5.3 | 6.9 | 0.8 |

| Ripening stage + trade preparation | Olive variety | n | mean | median | min | max | sd |
|---|---|---|---|---|---|---|---|
| Olives turning color, darkened by oxidation | Hoj | 3 | 5.1 | 5.1 | 5.1 | 5.1 | - |
| Olives turning color, darkened by oxidation | Man | 2 | 5.4 | 5.4 | 5.4 | 5.4 | - |
| Olives turning color, natural | Bel | 8 | 6.7 | 6.5 | 5.0 | 8.2 | 1.3 |
| Olives turning color, natural | Cai | 11 | 6.8 | 6.8 | 5.4 | 8.4 | 1.0 |
| Olives turning color, natural | Kon | 1 | 4.9 | 4.9 | 4.9 | 4.9 | - |
| Olives turning color, natural | Lad | 1 | 6.1 | 6.1 | 6.1 | 6.1 | - |
| Olives turning color, natural | Lec | 8 | 6.4 | 7.3 | na | 7.8 | 2.1 |
| Olives turning color, natural | Nob | 1 | 7.3 | 7.3 | 7.3 | 7.3 | - |
| Olives turning color, natural | Ogl | 1 | 7.9 | 7.9 | 7.9 | 7.9 | - |
| Olives turning color, natural | Per | 5 | 7.5 | 6.9 | 6.6 | 8.9 | 1.2 |
| Olives turning color, natural | Ter | 3 | 3.6 | na | na | 6.1 | 2.2 |
| Olives turning color, natural | Unk | 1 | 6.9 | 6.9 | 6.9 | 6.9 | - |

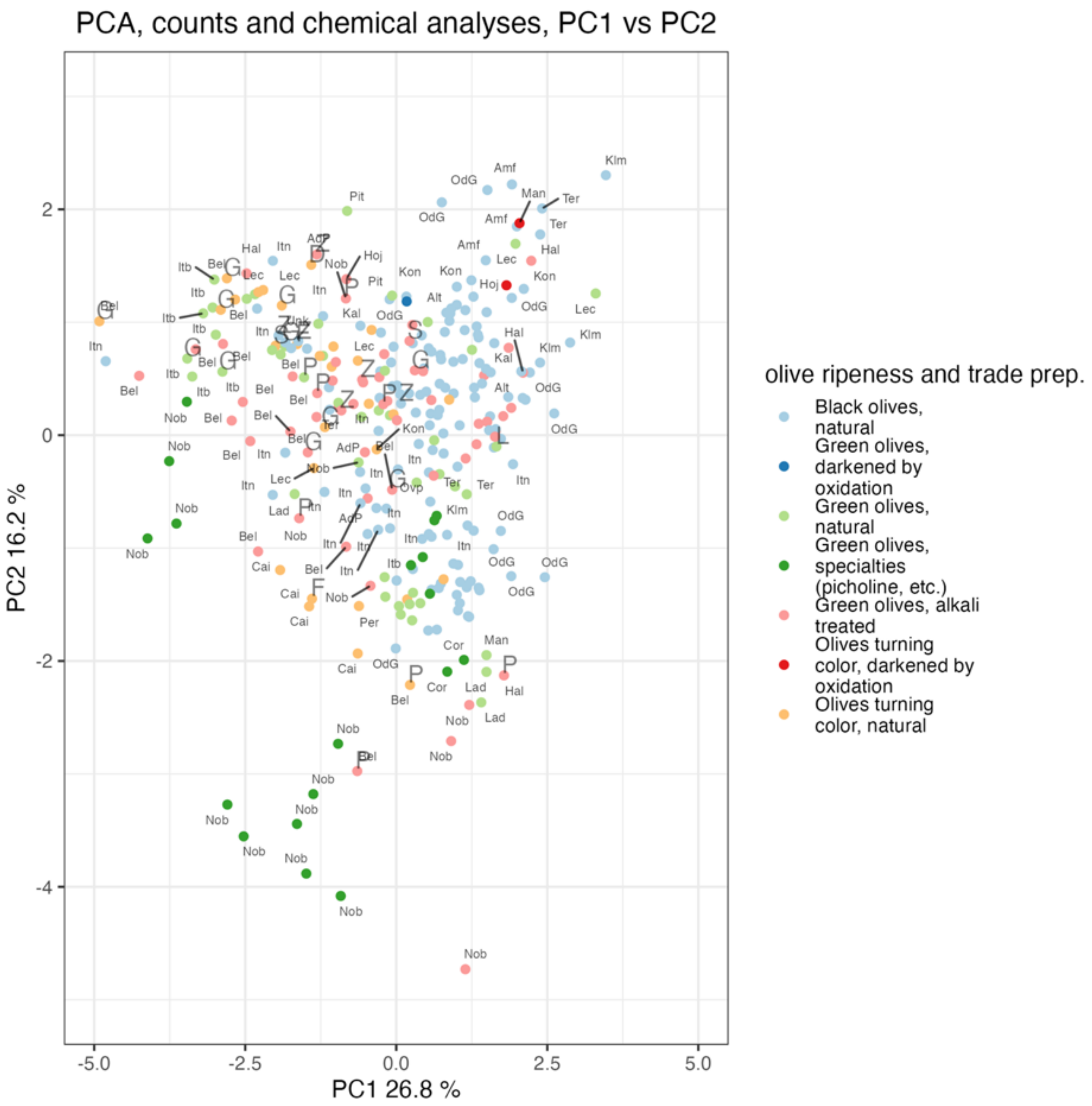


**Supplementary Figure 6**. Score plot for the first two components of a Principal Component Analysis carried out on the correlation matrix of the results of physico-chemical and chemical analysis and microbial counts for table olives used in this study. Abbreviations indicate the type of spoilage. D = Discolouration, F = Flor, G = Gas / Gas pochets, L = Larvae, P = Putrid, S = Spots, Z = Zapatera / palmiche

**Supplementary table 14**. Descriptive statistics for Chao1 by ripening stage and trade preparation for 16S rRNA metataxonomic data.

| Ripening stage + trade preparation | n | mean | median | min | max | sd |
|---|---|---|---|---|---|---|
| Black dehydrated | 1 | 38.3 | 38.3 | 38.3 | 38.3 | - |
| Black natural | 144 | 37.3 | 25.2 | 7.0 | 120.8 | 28.7 |
| Green alkali treated | 61 | 44.8 | 45.0 | 6.0 | 93.3 | 18.8 |
| Green darkened by oxidation | 3 | 25.7 | 22.0 | 13.0 | 42.0 | 14.8 |
| Green natural | 52 | 31.1 | 22.0 | 7.0 | 100.5 | 21.9 |
| Green specialties (picholine, etc.) | 27 | 17.9 | 16.2 | 3.0 | 48.0 | 12.4 |
| Turning color darkened by oxidation | 6 | 22.7 | 18.0 | 12.0 | 49.6 | 13.9 |
| Turning color natural | 35 | 43.1 | 38.5 | 10.0 | 121.5 | 31.1 |

**Supplementary Table 15**. Descriptive statistics for Chao1 by ripening stage and trade preparation and olive variety (see Supplementary Table 2 for abbreviations).

| Ripening stage + trade preparation | Olive variety | n | mean | median | min | max | sd |
|---|---|---|---|---|---|---|---|
| Black dehydrated | Ofe | 1 | 38.3 | 38.3 | 38.3 | 38.3 | - |
| Black natural | Alt | 4 | 24.1 | 19.0 | 17.0 | 41.3 | 11.6 |
| Black natural | Amf | 4 | 35.2 | 18.5 | 12.0 | 92.0 | 38.0 |
| Black natural | Cri | 4 | 30.0 | 13.5 | 7.0 | 86.0 | 37.5 |
| Black natural | Itn | 34 | 32.1 | 17.5 | 8.8 | 120.8 | 29.6 |
| Black natural | Kal | 12 | 48.6 | 40.0 | 16.0 | 96.6 | 28.5 |
| Black natural | Klm | 7 | 49.4 | 55.8 | 25.3 | 78.1 | 18.4 |
| Black natural | Kon | 9 | 34.3 | 28.0 | 21.0 | 83.5 | 19.4 |
| Black natural | Lad | 3 | 70.9 | 82.9 | 34.8 | 95.1 | 31.9 |
| Black natural | OdG | 61 | 34.1 | 20.0 | 7.0 | 117.0 | 27.0 |
| Black natural | Per | 1 | 42.5 | 42.5 | 42.5 | 42.5 | - |
| Black natural | Pic | 2 | 113.1 | 113.1 | 106.5 | 119.7 | 9.3 |
| Black natural | Tag | 1 | 52.0 | 52.0 | 52.0 | 52.0 | - |
| Black natural | Ter | 2 | 33.5 | 33.5 | 30.0 | 37.0 | 4.9 |
| Green alkali treated | AdP | 6 | 34.9 | 36.5 | 6.0 | 60.2 | 19.2 |
| Green alkali treated | Bel | 18 | 47.6 | 46.0 | 28.0 | 69.0 | 13.8 |
| Green alkali treated | Cac | 1 | 59.7 | 59.7 | 59.7 | 59.7 | - |
| Green alkali treated | Gor | 1 | 93.3 | 93.3 | 93.3 | 93.3 | - |
| Green alkali treated | Hal | 20 | 43.9 | 46.8 | 11.0 | 72.5 | 21.2 |
| Green alkali treated | Hoj | 4 | 52.7 | 43.0 | 37.0 | 87.8 | 23.9 |
| Green alkali treated | Man | 3 | 34.6 | 32.2 | 31.5 | 40.2 | 4.8 |

| Ripening stage + trade preparation | Olive variety | n | mean | median | min | max | sd |
|---|---|---|---|---|---|---|---|
| Green alkali treated | Nob | 7 | 35.6 | 31.0 | 25.0 | 57.0 | 11.5 |
| Green alkali treated | Plu | 1 | 69.1 | 69.1 | 69.1 | 69.1 | - |
| Green darkened by oxidation | Cac | 2 | 27.5 | 27.5 | 13.0 | 42.0 | 20.5 |
| Green darkened by oxidation | Man | 1 | 22.0 | 22.0 | 22.0 | 22.0 | - |
| Green natural | AdP | 1 | 43.0 | 43.0 | 43.0 | 43.0 | - |
| Green natural | Bec | 1 | 18.0 | 18.0 | 18.0 | 18.0 | - |
| Green natural | Bel | 1 | 39.3 | 39.3 | 39.3 | 39.3 | - |
| Green natural | Cac | 1 | 19.0 | 19.0 | 19.0 | 19.0 | - |
| Green natural | Car | 1 | 11.3 | 11.3 | 11.3 | 11.3 | - |
| Green natural | Hoj | 2 | 61.0 | 61.0 | 53.0 | 69.0 | 11.3 |
| Green natural | Itb | 16 | 28.9 | 18.5 | 9.0 | 100.5 | 26.7 |
| Green natural | Lad | 3 | 15.8 | 16.0 | 11.3 | 20.2 | 4.4 |
| Green natural | Lec | 2 | 18.4 | 18.4 | 17.0 | 19.8 | 1.9 |
| Green natural | Man | 2 | 35.4 | 35.4 | 34.0 | 36.8 | 1.9 |
| Green natural | Nob | 5 | 27.2 | 27.0 | 21.0 | 32.0 | 4.2 |
| Green natural | Nom | 1 | 22.0 | 22.0 | 22.0 | 22.0 | - |
| Green natural | Ovp | 2 | 85.1 | 85.1 | 72.0 | 98.1 | 18.5 |
| Green natural | Pit | 4 | 18.8 | 17.0 | 16.0 | 25.0 | 4.2 |
| Green natural | TdC | 5 | 21.4 | 28.0 | 7.0 | 31.0 | 11.5 |
| Green natural | Ter | 5 | 46.8 | 42.0 | 40.5 | 66.5 | 11.0 |
| Green specialties (picholine, etc.) | Cor | 2 | 18.8 | 18.8 | 16.5 | 21.0 | 3.2 |

| Ripening stage + trade preparation | Olive variety | n | mean | median | min | max | sd |
|---|---|---|---|---|---|---|---|
| Green specialties (picholine, etc.) | Nob | 19 | 15.8 | 11.5 | 3.0 | 48.0 | 13.4 |
| Green specialties (picholine, etc.) | Oka | 3 | 23.0 | 17.5 | 17.2 | 34.2 | 9.7 |
| Green specialties (picholine, etc.) | Sal | 3 | 25.5 | 27.1 | 15.0 | 34.5 | 9.8 |
| Turning color darkened by oxidation | Cac | 4 | 27.3 | 22.2 | 15.0 | 49.6 | 15.3 |
| Turning color darkened by oxidation | Hoj | 1 | 15.0 | 15.0 | 15.0 | 15.0 | - |
| Turning color darkened by oxidation | Man | 1 | 12.0 | 12.0 | 12.0 | 12.0 | - |
| Turning color natural | Bel | 8 | 29.2 | 25.5 | 11.0 | 54.0 | 15.9 |
| Turning color natural | Cai | 9 | 34.1 | 47.0 | 10.0 | 55.2 | 20.9 |
| Turning color natural | Kon | 1 | 89.1 | 89.1 | 89.1 | 89.1 | - |
| Turning color natural | Lad | 1 | 14.0 | 14.0 | 14.0 | 14.0 | - |
| Turning color natural | Lec | 6 | 48.0 | 36.4 | 25.2 | 121.1 | 36.5 |
| Turning color natural | Nob | 1 | 15.0 | 15.0 | 15.0 | 15.0 | - |
| Turning color natural | Ogl | 1 | 121.5 | 121.5 | 121.5 | 121.5 | - |
| Turning color natural | Per | 3 | 56.6 | 56.2 | 31.5 | 82.0 | 25.3 |
| Turning color natural | Tag | 1 | 120.8 | 120.8 | 120.8 | 120.8 | - |
| Turning color natural | Ter | 3 | 44.4 | 42.2 | 39.0 | 52.0 | 6.8 |

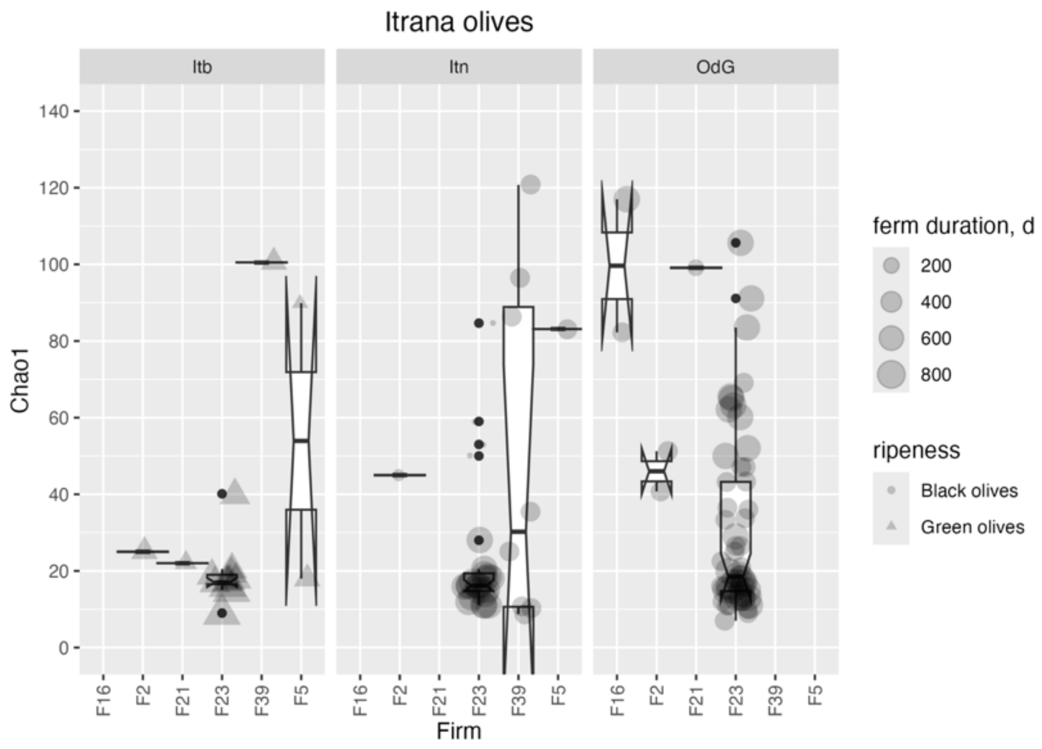


**Supplementary Figure 7**. Distribution of Chao1 values for bacterial communities of Itrana olives (Itb = Itrana bianca, Itn = Itrana nera, OdG = Oliva di Gaeta PDO). The size of the symbols is made proportional to the duration of fermentation.

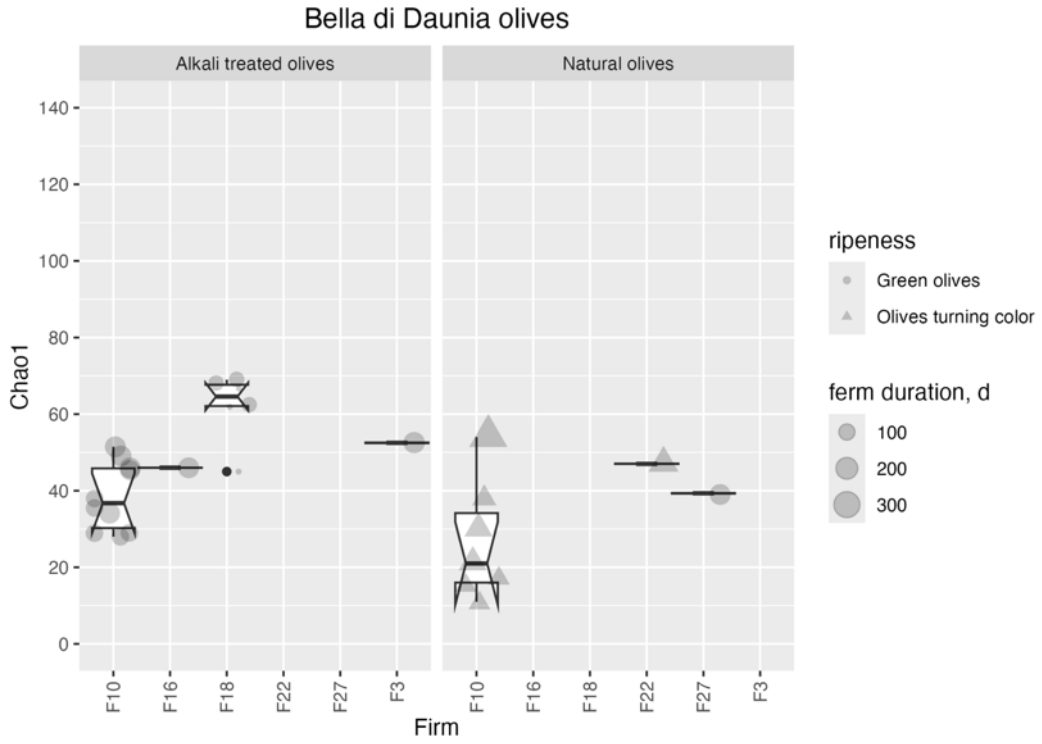


**Supplementary Figure 8.** Distribution of Chao1 values for bacterial communities of Bella di Daunia PDO olives produced by different firms. The size of the symbols is made proportional to the duration of fermentation.

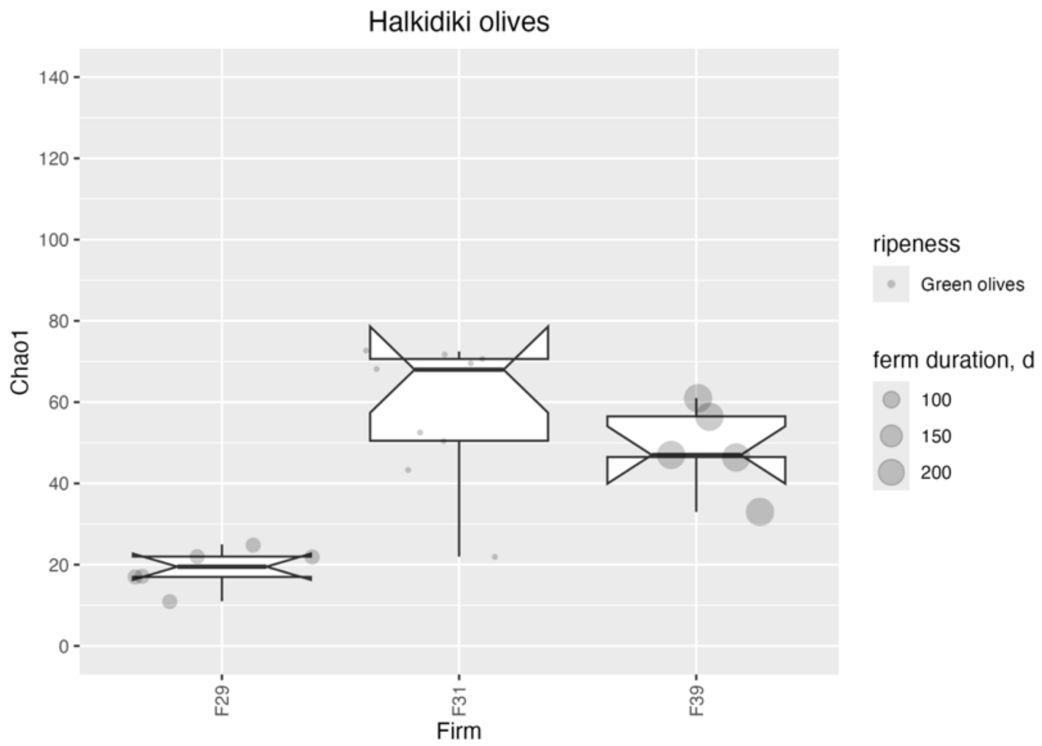


**Supplementary Figure 9.** Distribution of Chao1 values for bacterial communities of Green alkali treated Halkidiki olives. The size of the symbols is made proportional to the duration of fermentation.

**Supplementary Table 16**. Spearman ρ correlation values and p values for a two tail test with $H_0$:r=0 for alpha diversity for bacterial communities and selected variables. Only correlations with p<0.05 after Holm p-value correction for multiple testing are shown.

| Ripening stage + trade preparation | var1 | var2 | cor | statistic | p |
|---|---|---|---|---|---|
| Black natural | Chao1 | Duration of fermentation | -0.39 | 634111.708 | 2.40e-06 |
| Green natural | Chao1 | Duration of fermentation | -0.42 | 24561.344 | 3.29e-03 |
| Black natural | Chao1 | LAB | -0.35 | 616181.256 | 2.61e-05 |
| Green alkali treated | Chao1 | LAB | 0.31 | 20066.916 | 1.84e-02 |
| Green natural | Chao1 | LAB | -0.38 | 30466.621 | 6.16e-03 |
| Green alkali treated | Chao1 | YEASTMOLDS | 0.34 | 19374.946 | 1.09e-02 |
| Green natural | Chao1 | YEASTMOLDS | -0.30 | 28678.190 | 3.39e-02 |
| Black natural | Chao1 | conc_NaCl | 0.20 | 367384.862 | 1.99e-02 |
| Green alkali treated | Chao1 | conc_NaCl | -0.37 | 40133.159 | 4.80e-03 |
| Green natural | Chao1 | conc_NaCl | 0.37 | 13840.344 | 6.90e-03 |
| Green specialties (picholine, etc.) | Chao1 | conc_NaCl | -0.52 | 3081.045 | 1.06e-02 |
| Black natural | Chao1 | pH | 0.30 | 318194.148 | 2.58e-04 |
| Green specialties (picholine, etc.) | Chao1 | pH | -0.72 | 3472.147 | 1.24e-04 |
| Turning color natural | Chao1 | pH | 0.41 | 2941.261 | 2.31e-02 |

**Supplementary Table 17.** Descriptive statistics for Chao1 for fungal communities of table olives by ripening stage and trade preparation. NA not available.

| Olive ripeness + trade prep. | n | mean | median | min | max | sd |
|---|---|---|---|---|---|---|
| Black dehydrated | 1 | 12.0 | 12.0 | 12 | 12.0 | - |
| Black natural | 145 | 11.7 | 9.0 | 3 | 51.1 | 9.6 |
| Green alkali treated | 63 | 18.2 | 15.0 | 5 | 56.9 | 11.0 |
| Green darkened by oxidation | 3 | 9.0 | 10.0 | 5 | 12.0 | 3.6 |
| Green natural | 54 | 11.3 | 11.0 | 1 | 25.8 | 5.8 |
| Green specialties (picholine, etc.) | 27 | 54.8 | 37.8 | 4 | 209.0 | 53.4 |
| Turning color darkened by oxidation | 5 | 13.5 | 13.3 | 4 | 21.0 | 7.4 |
| Turning color natural | 32 | 15.6 | 12.5 | 2 | 44.2 | 12.1 |

**Supplementary Table 18**. Descriptive statistics for Chao1 for fungal communities by ripening stage and trade preparation and olive variety (see Supplementary table 2 for abbreviations).

| Olive ripeness + trade prep. | Olive variety | n | mean | median | min | max | sd |
|---|---|---|---|---|---|---|---|
| Black dehydrated | Ofe | 1 | 12.0 | 12.0 | 12.0 | 12.0 | - |
| Black natural | Alt | 4 | 10.4 | 10.2 | 9.0 | 12.0 | 1.6 |
| Black natural | Amf | 4 | 20.2 | 20.5 | 13.0 | 27.0 | 7.3 |
| Black natural | Cri | 4 | 7.2 | 5.5 | 5.0 | 13.0 | 3.9 |
| Black natural | Itn | 35 | 7.5 | 6.0 | 3.0 | 34.0 | 6.0 |
| Black natural | Kal | 13 | 18.2 | 13.0 | 6.0 | 50.7 | 12.7 |
| Black natural | Klm | 7 | 37.9 | 39.0 | 23.0 | 51.1 | 8.7 |
| Black natural | Kon | 9 | 11.8 | 12.3 | 7.0 | 16.8 | 3.1 |
| Black natural | Lad | 3 | 14.7 | 11.0 | 8.0 | 25.2 | 9.2 |
| Black natural | OdG | 61 | 8.3 | 8.0 | 3.0 | 26.6 | 4.1 |
| Black natural | Per | 1 | 15.0 | 15.0 | 15.0 | 15.0 | - |
| Black natural | Pic | 2 | 24.3 | 24.3 | 20.2 | 28.4 | 5.8 |
| Black natural | Ter | 2 | 31.9 | 31.9 | 31.8 | 32.1 | 0.3 |
| Green alkali treated | AdP | 6 | 18.4 | 14.0 | 5.0 | 49.5 | 16.6 |
| Green alkali treated | Bel | 19 | 15.7 | 12.0 | 7.0 | 56.9 | 11.6 |
| Green alkali treated | Cac | 1 | 18.0 | 18.0 | 18.0 | 18.0 | - |
| Green alkali treated | Gor | 1 | 20.5 | 20.5 | 20.5 | 20.5 | - |
| Green alkali treated | Hal | 21 | 19.8 | 20.0 | 5.0 | 41.7 | 11.1 |
| Green alkali treated | Hoj | 4 | 18.5 | 15.5 | 5.0 | 38.0 | 13.9 |
| Green alkali treated | Man | 3 | 14.4 | 13.0 | 11.0 | 19.3 | 4.4 |
| Green alkali treated | Nob | 7 | 21.9 | 23.0 | 9.0 | 29.0 | 6.6 |
| Green alkali treated | Plu | 1 | 14.0 | 14.0 | 14.0 | 14.0 | - |

| Olive ripeness + trade prep. | Olive variety | n | mean | median | min | max | sd |
|---|---|---|---|---|---|---|---|
| Green darkened by oxidation | Cac | 2 | 7.5 | 7.5 | 5.0 | 10.0 | 3.5 |
| Green darkened by oxidation | Man | 1 | 12.0 | 12.0 | 12.0 | 12.0 | - |
| Green natural | AdP | 1 | 4.0 | 4.0 | 4.0 | 4.0 | - |
| Green natural | Bec | 1 | 8.0 | 8.0 | 8.0 | 8.0 | - |
| Green natural | Bel | 1 | 11.0 | 11.0 | 11.0 | 11.0 | - |
| Green natural | Car | 1 | 6.0 | 6.0 | 6.0 | 6.0 | - |
| Green natural | Hoj | 2 | 10.8 | 10.8 | 6.0 | 15.5 | 6.7 |
| Green natural | Itb | 16 | 7.7 | 6.0 | 3.0 | 20.0 | 4.9 |
| Green natural | Lad | 3 | 11.9 | 9.0 | 6.3 | 20.5 | 7.5 |
| Green natural | Lec | 2 | 14.0 | 14.0 | 7.0 | 21.0 | 9.9 |
| Green natural | Man | 2 | 10.0 | 10.0 | 7.0 | 13.0 | 4.2 |
| Green natural | Nob | 5 | 14.6 | 15.0 | 9.0 | 18.0 | 3.5 |
| Green natural | Nom | 1 | 12.0 | 12.0 | 12.0 | 12.0 | - |
| Green natural | Ovp | 2 | 14.6 | 14.6 | 12.2 | 17.0 | 3.4 |
| Green natural | Pit | 4 | 18.9 | 19.5 | 11.0 | 25.8 | 6.4 |
| Green natural | TdC | 9 | 12.2 | 14.0 | 1.0 | 18.0 | 5.3 |
| Green natural | Ter | 4 | 13.7 | 12.9 | 8.0 | 21.0 | 5.5 |
| Green specialties (picholine, etc.) | Cor | 3 | 19.1 | 15.5 | 4.0 | 37.8 | 17.2 |
| Green specialties (picholine, etc.) | Nob | 18 | 62.8 | 56.7 | 4.0 | 209.0 | 58.4 |
| Green specialties (picholine, etc.) | Oka | 3 | 18.0 | 17.3 | 15.0 | 21.6 | 3.3 |
| Green specialties (picholine, etc.) | Sal | 3 | 79.5 | 66.5 | 38.3 | 133.6 | 48.9 |

| Olive ripeness + trade prep. | Olive variety | n | mean | median | min | max | sd |
|---|---|---|---|---|---|---|---|
| Turning color darkened by oxidation | Cac | 4 | 14.7 | 16.9 | 4.0 | 21.0 | 8.0 |
| Turning color darkened by oxidation | Man | 1 | 8.5 | 8.5 | 8.5 | 8.5 | - |
| Turning color natural | Bel | 8 | 7.6 | 5.0 | 3.0 | 17.0 | 4.7 |
| Turning color natural | Cai | 7 | 21.6 | 26.0 | 2.0 | 43.3 | 17.2 |
| Turning color natural | Kon | 1 | 26.2 | 26.2 | 26.2 | 26.2 | - |
| Turning color natural | Lad | 1 | 6.0 | 6.0 | 6.0 | 6.0 | - |
| Turning color natural | Lec | 6 | 16.2 | 14.8 | 11.0 | 25.0 | 5.0 |
| Turning color natural | Nob | 1 | 5.0 | 5.0 | 5.0 | 5.0 | - |
| Turning color natural | Ogl | 1 | 13.0 | 13.0 | 13.0 | 13.0 | - |
| Turning color natural | Per | 2 | 12.8 | 12.8 | 10.0 | 15.5 | 3.9 |
| Turning color natural | Tag | 1 | 44.2 | 44.2 | 44.2 | 44.2 | - |
| Turning color natural | Ter | 3 | 19.7 | 19.5 | 4.0 | 35.6 | 15.8 |
| Turning color natural | Unk | 1 | 11.0 | 11.0 | 11.0 | 11.0 | - |
| Black dehydrated | Ofe | 1 | 12.0 | 12.0 | 12.0 | 12.0 | - |
| Black natural | Alt | 4 | 10.4 | 10.2 | 9.0 | 12.0 | 1.6 |

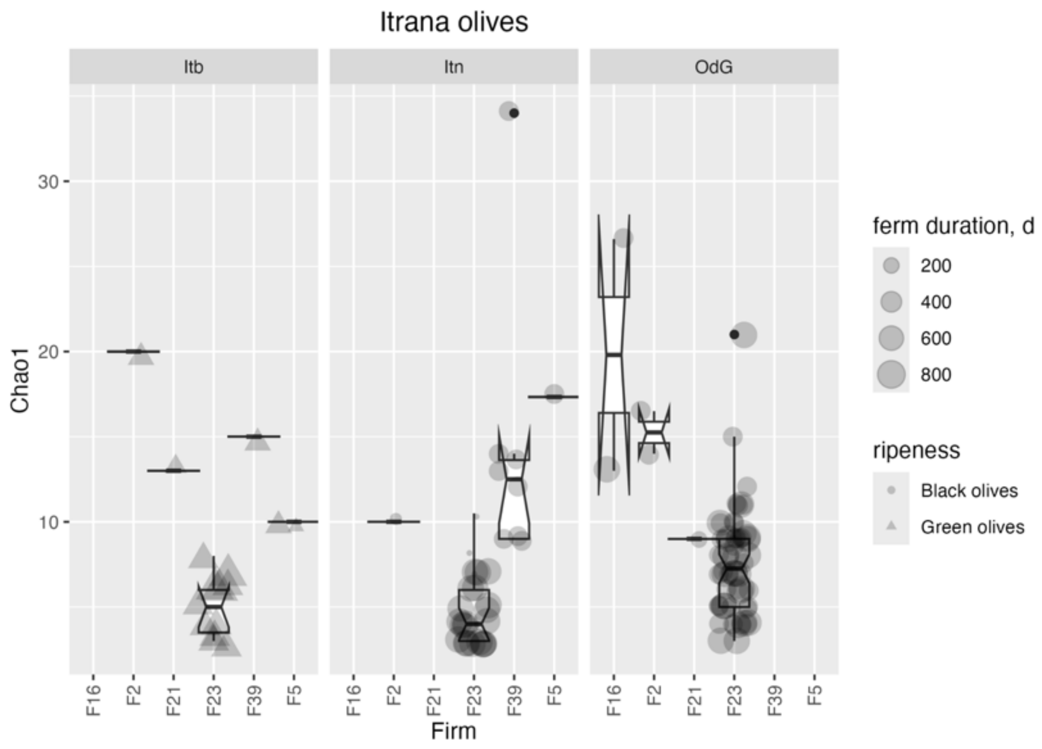


**Supplementary Figure 10.** Fungal diversity for Itrana olives (Itb = Itrana bianca; Itn = Itrana nera; OdG = Oliva di Gaeta PDO)

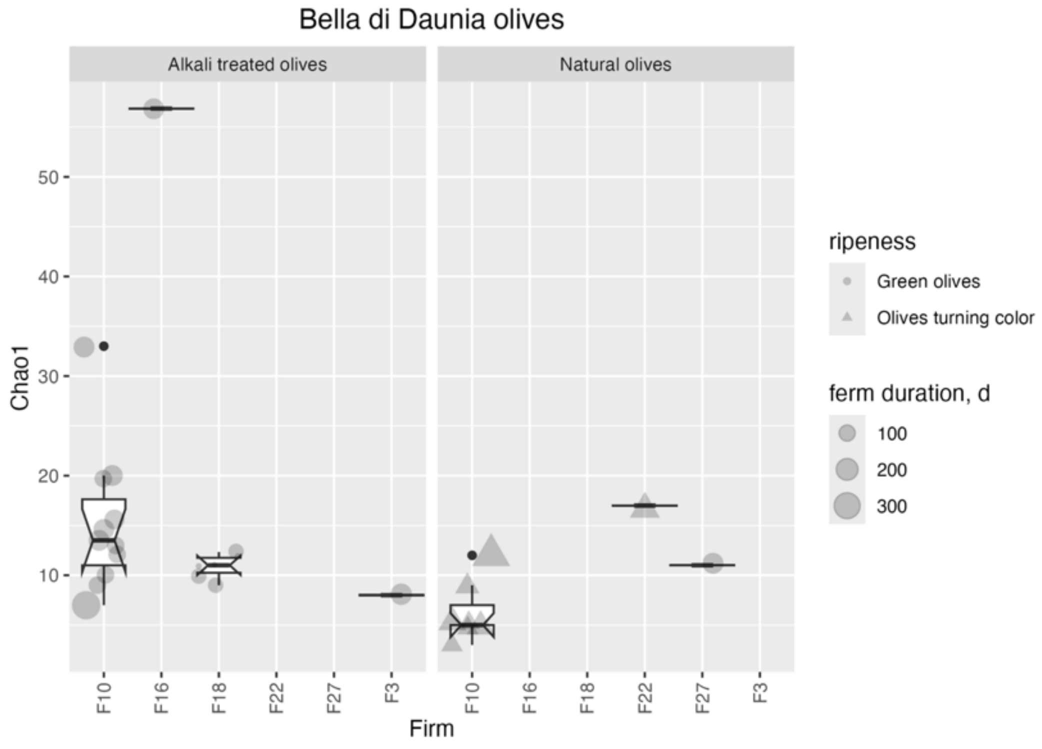


**Supplementary Figure 11**. Fungal diversity for Bella di Daunia PDO olives

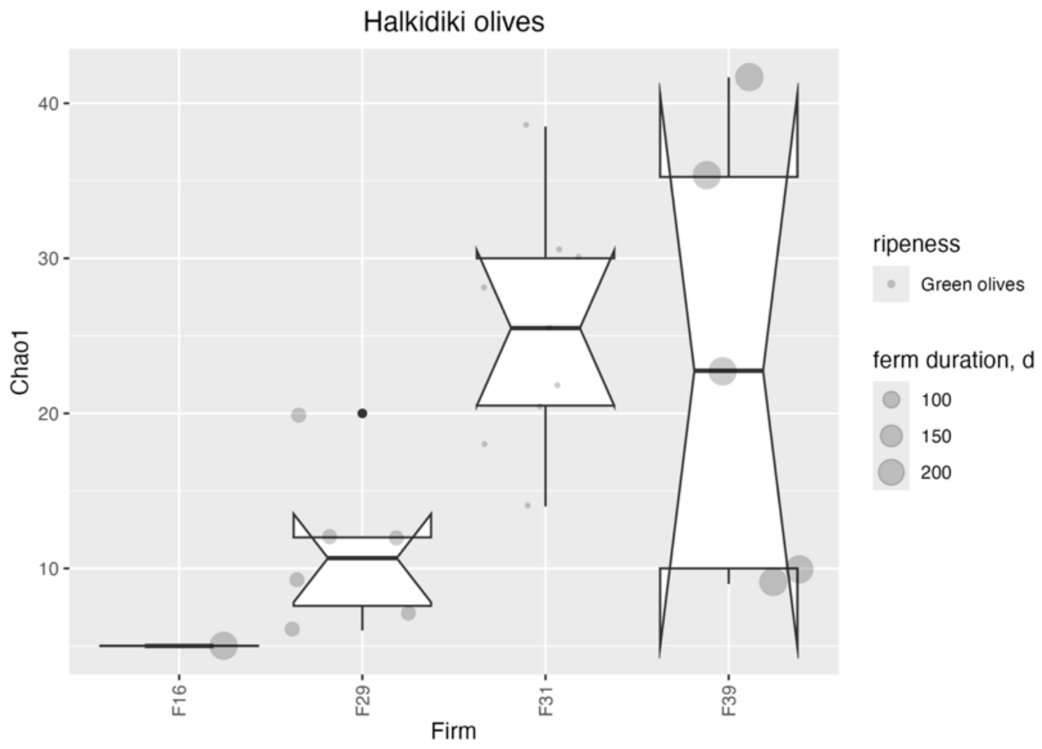


**Supplementary Figure 12**. Fungal diversity for Halkidiki olives.

**Supplementary Table 19**. Spearman ρ correlation values and p values for a two-tailed test with $H_0$:r=0 for alpha diversity for fungal communities and selected variables. Only correlations with p<0.05 after Holm p-value correction for multiple testing are shown.

| Olive ripeness + trade prep. | var1 | var2 | cor | statistic | p |
|---|---|---|---|---|---|
| Black natural | Chao1 | Duration of fermentation | -0.62 | 757949.9442 | 0.00e+00 |
| Green natural | Chao1 | Duration of fermentation | -0.53 | 31795.7468 | 8.49e-05 |
| Turning color natural | Chao1 | Duration of fermentation | 0.38 | 3082.7494 | 3.58e-02 |
| Green natural | Chao1 | LAB | -0.29 | 33858.7105 | 3.30e-02 |
| Black natural | Chao1 | YEASTMOLDS | -0.43 | 668785.7189 | 1.00e-07 |
| Green alkali treated | Chao1 | YEASTMOLDS | -0.32 | 42962.3266 | 1.38e-02 |
| Green natural | Chao1 | YEASTMOLDS | -0.60 | 42036.8438 | 1.50e-06 |
| Green specialties (picholine, etc.) | Chao1 | YEASTMOLDS | -0.46 | 3365.3340 | 2.26e-02 |
| Black natural | Chao1 | conc_NaCl | 0.34 | 308580.9520 | 3.82e-05 |
| Green alkali treated | Chao1 | conc_NaCl | -0.26 | 41035.6902 | 4.67e-02 |
| Green natural | Chao1 | conc_NaCl | 0.38 | 16362.8239 | 5.04e-03 |
| Green specialties (picholine, etc.) | Chao1 | pH | 0.59 | 941.5234 | 2.38e-03 |

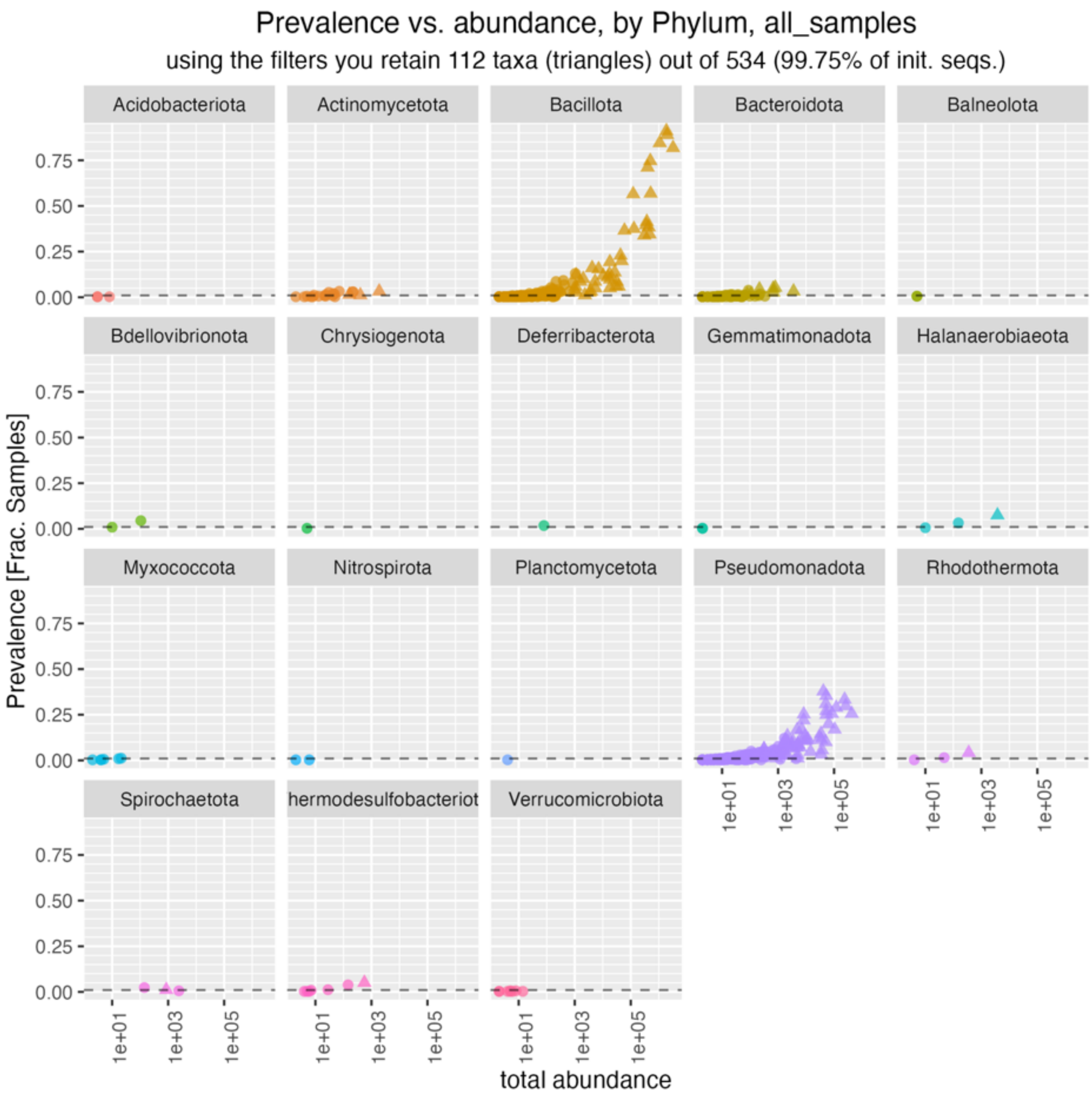


**Supplementary Figure 13**. Prevalence and abundance of bacterial genera in table olives used in this study; only taxa which had a maximum abundance >1% and a prevalence >1% were retained.

**Supplementary Table 20**. Top 24 bacterial genera in table olives in terms of prevalence and/or abundance. Sort_var is the score for the first principal component.

| label | Phylum | Class | Mean relative abundance | Prevalence | sort_var |
|---|---|---|---|---|---|
| *Lentilactobacillus* | Bacillota | Bacilli | 0.229 | 0.818 | 16.500 |
| *Pediococcus* | Bacillota | Bacilli | 0.140 | 0.890 | 12.395 |
| *Lactiplantibacillus* | Bacillota | Bacilli | 0.132 | 0.911 | 12.128 |
| *Secundilactobacillus* | Bacillota | Bacilli | 0.076 | 0.845 | 8.889 |
| *Loigolactobacillus* | Bacillota | Bacilli | 0.035 | 0.747 | 6.160 |
| *Paucilactobacillus* | Bacillota | Bacilli | 0.028 | 0.711 | 5.582 |
| *Levilactobacillus* | Bacillota | Bacilli | 0.036 | 0.568 | 5.087 |
| *Lacticaseibacillus* | Bacillota | Bacilli | 0.009 | 0.565 | 3.655 |
| *Alkalibacterium* | Bacillota | Bacilli | 0.031 | 0.384 | 3.624 |
| *Marinilactibacillus* | Bacillota | Bacilli | 0.026 | 0.414 | 3.600 |
| *Weissella* | Bacillota | Bacilli | 0.033 | 0.345 | 3.500 |
| *Lactobacillus* | Bacillota | Bacilli | 0.025 | 0.396 | 3.409 |
| *Leuconostoc* | Bacillota | Bacilli | 0.021 | 0.339 | 2.871 |
| *Celerinatantimonas* | Pseudomonadota | Gammaproteobacteria | 0.030 | 0.256 | 2.790 |
| *Enterobacter* | Pseudomonadota | Gammaproteobacteria | 0.017 | 0.333 | 2.599 |
| *Enterococcus* | Bacillota | Bacilli | 0.009 | 0.375 | 2.473 |
| *Lelliottia* | Pseudomonadota | Gammaproteobacteria | 0.018 | 0.298 | 2.455 |
| *Acinetobacter* | Pseudomonadota | Gammaproteobacteria | 0.004 | 0.387 | na2 |
| *Pseudomonas* | Pseudomonadota | Gammaproteobacteria | 0.003 | 0.381 | 2.199 |
| *Ligilactobacillus* | Bacillota | Bacilli | 0.004 | 0.366 | 2.164 |
| *Suttonella* | Pseudomonadota | Gammaproteobacteria | 0.008 | 0.289 | 1.886 |

**Supplementary Table 20**. Top 24 bacterial genera in table olives in terms of prevalence and/or abundance. Sort_var is the score for the first principal component.

| label | Phylum | Class | Mean relative abundance | Prevalence | sort_var |
|---|---|---|---|---|---|
| *Halomonas* | Pseudomonadota | Gammaproteobacteria | 0.004 | 0.310 | 1.771 |
| *Pantoea* | Pseudomonadota | Gammaproteobacteria | 0.006 | 0.256 | 1.551 |
| *Salinicola* | Pseudomonadota | Gammaproteobacteria | 0.004 | 0.271 | 1.532 |

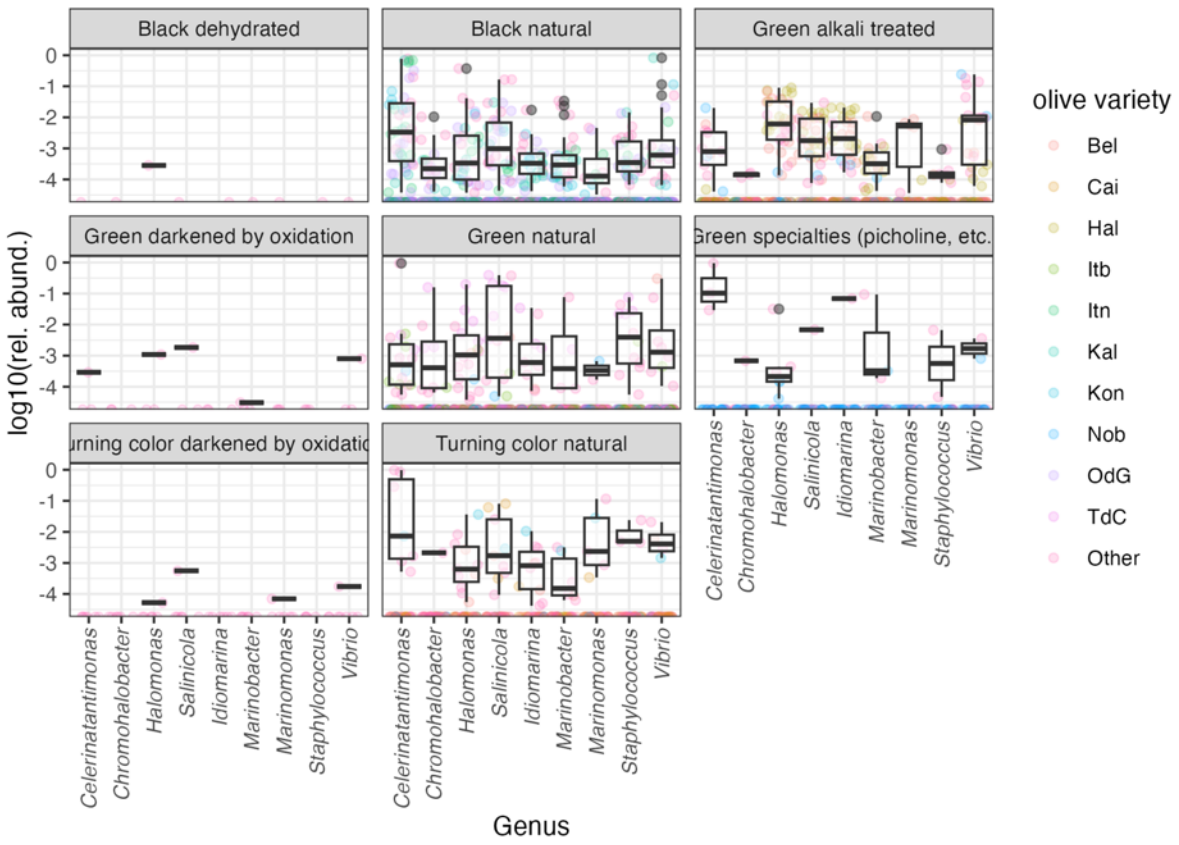


**Supplementary Figure 14**. Distribution of the relative abundance of halophilic genera (other than LAB and HALAB) in table olives.

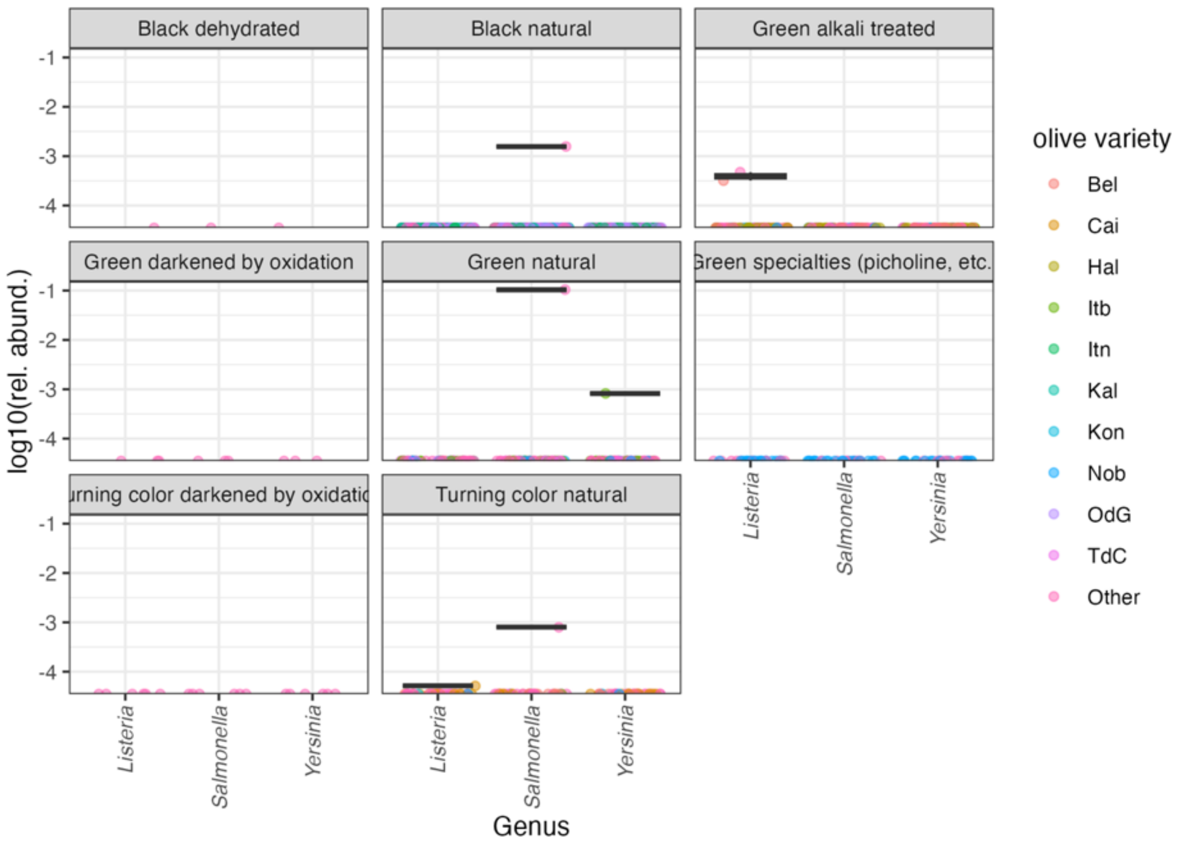


**Supplementary Figure 15**. Distribution of the relative abundance of DNA sequences assigned to bacterial genera which include pathogenic species in table olives.

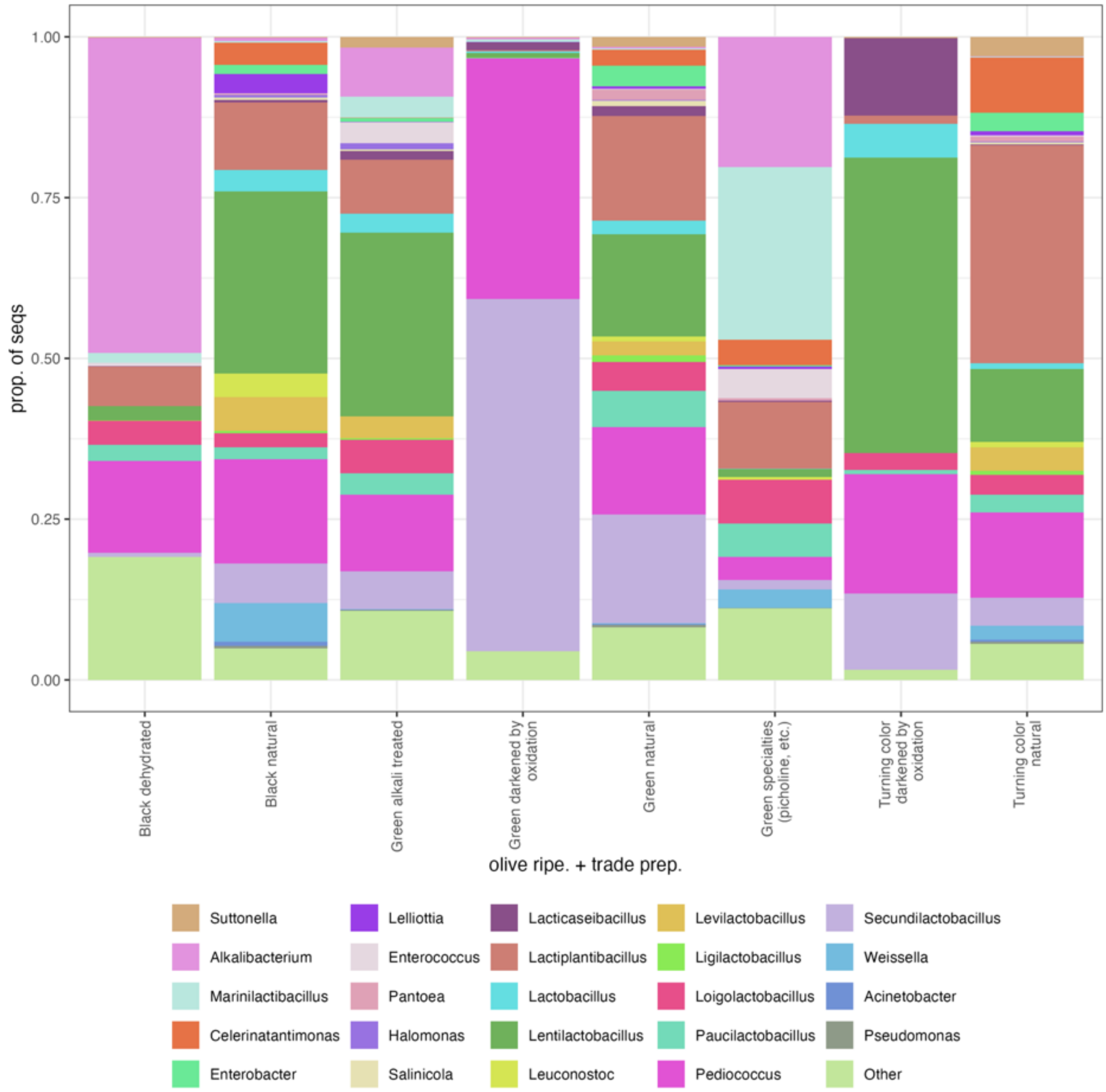


**Supplementary Figure 16**. Average abundance of the most prevalent and abundant bacterial genera in table olives, by combinations of ripeness and trade preparation. “Other” represent the combined relative abundance of all other genera.

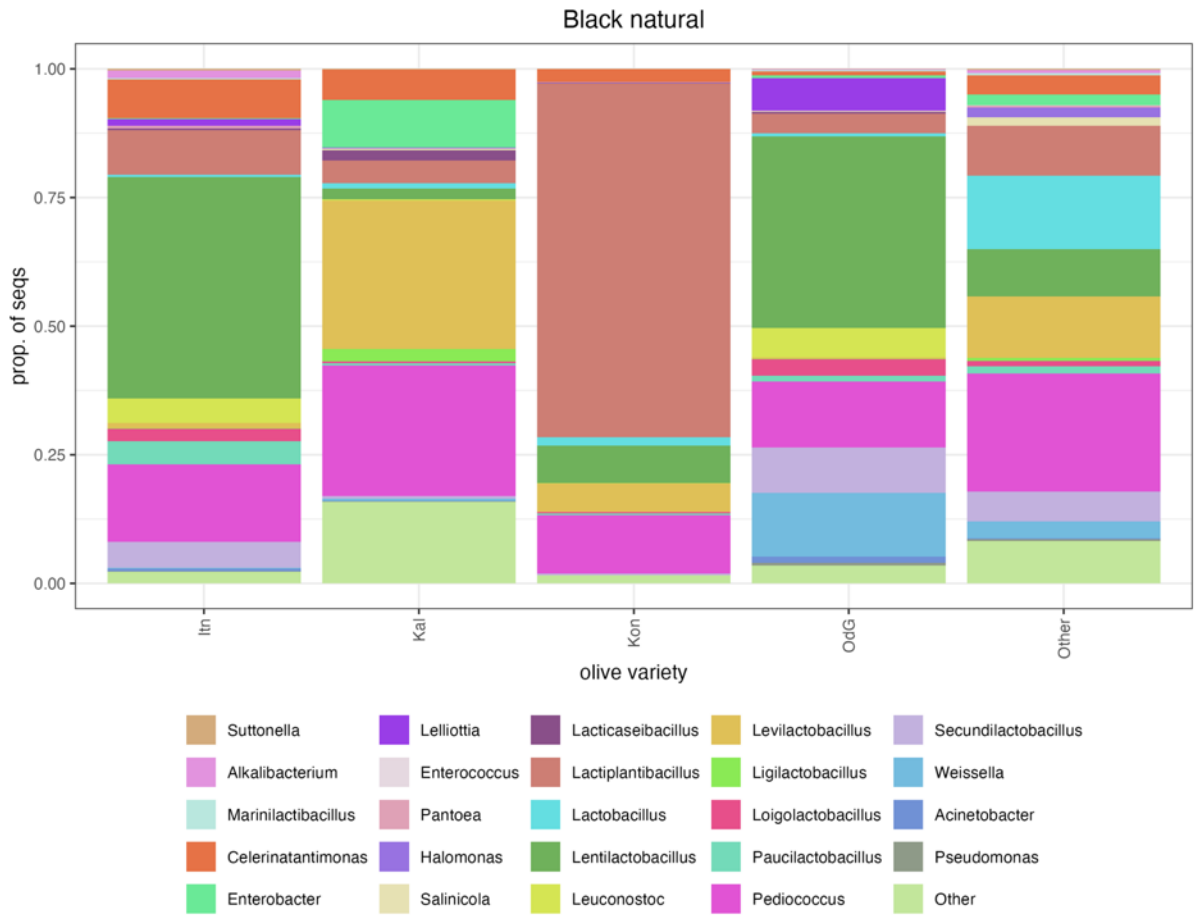


**Supplementary Figure 17.** Average abundance of the most prevalent and abundant bacterial genera in black natural olives. The main varieties (those with >9 samples) are shown in individual columns, while the others are pooled.

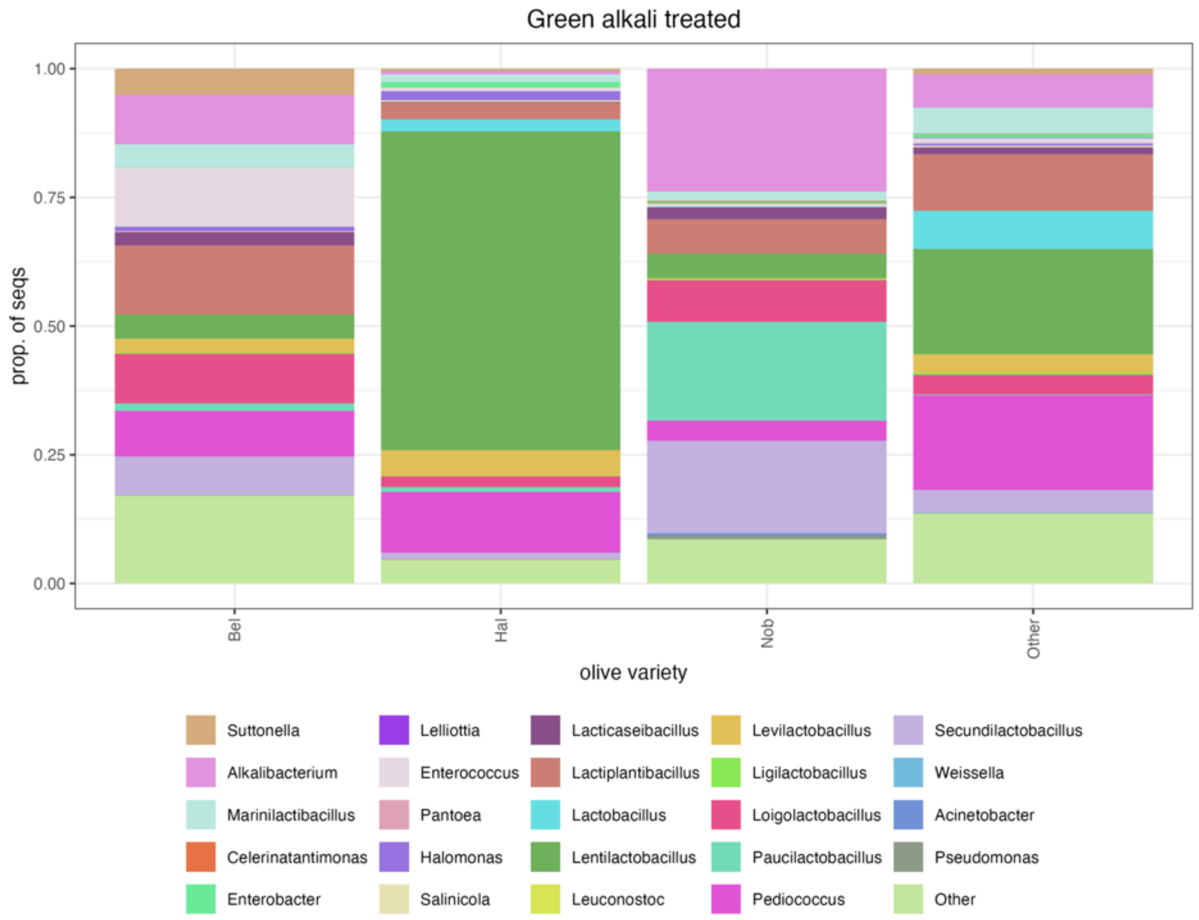


**Supplementary Figure 18.** Average abundance of the most prevalent and abundant bacterial genera in green alkali treated olives. The main varieties (those with >9 samples) are shown in individual columns, while the others are pooled.

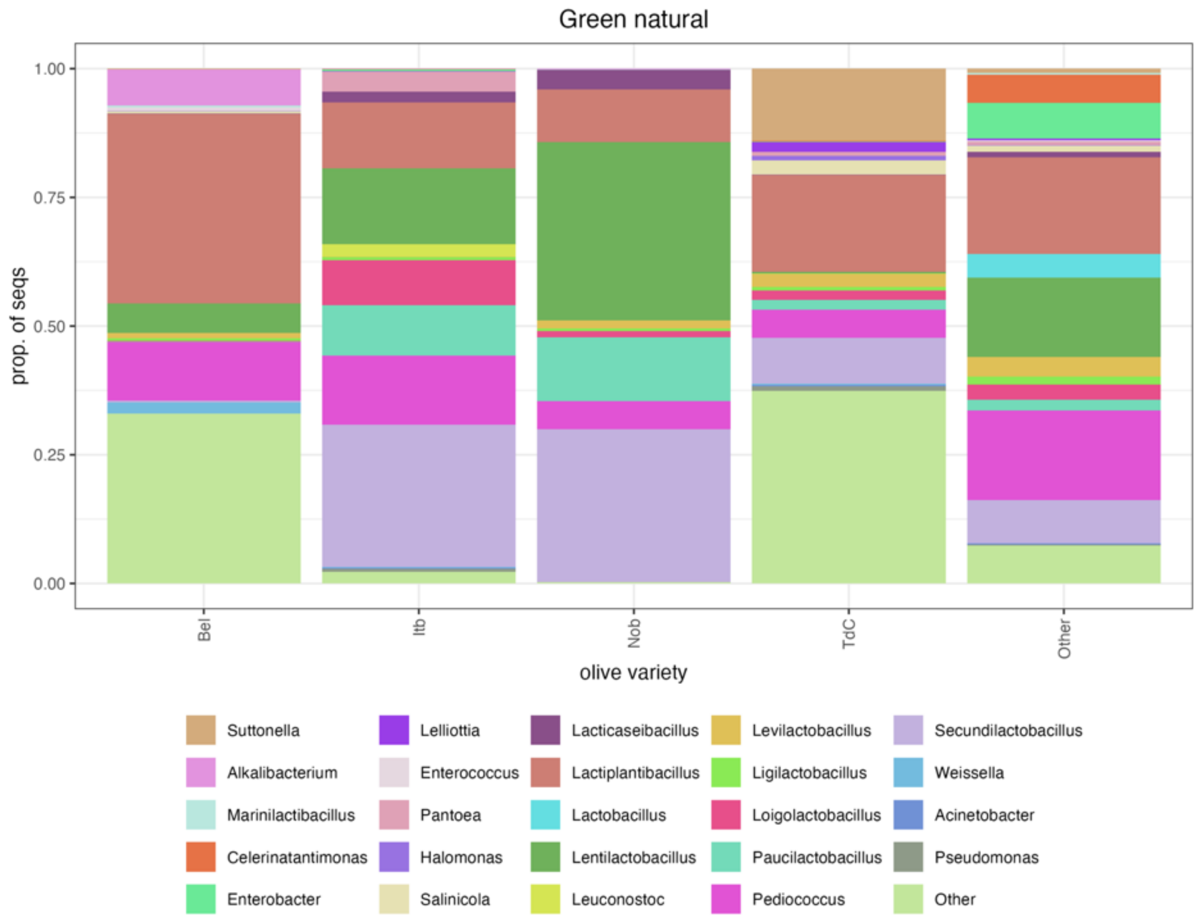


**Supplementary Figure 19.** Average abundance of the most prevalent and abundant bacterial genera in green naturally fermented olives. The main varieties (those with >9 samples) are shown in individual columns, while the others are pooled.

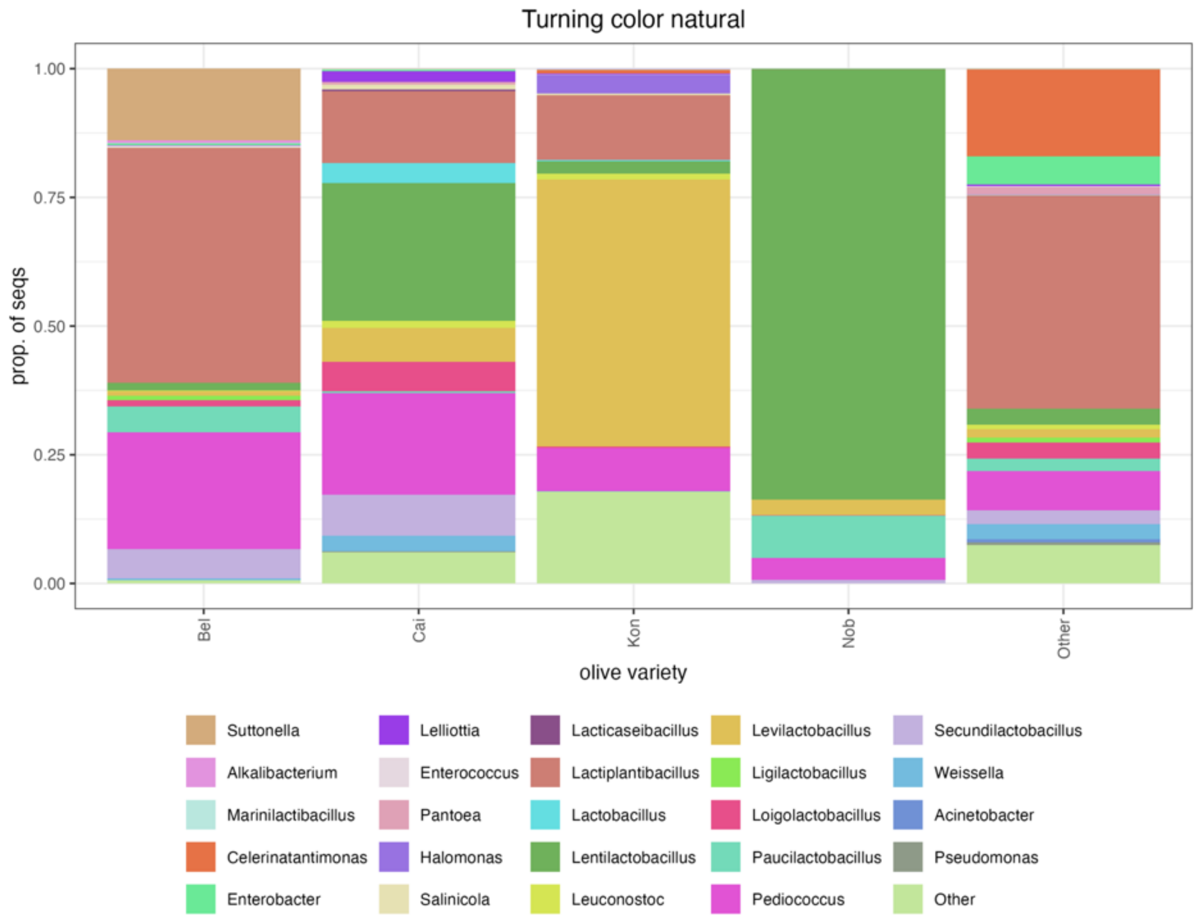


**Supplementary Figure 20**. Average abundance of the most prevalent and abundant bacterial genera in turning color naturally fermented olives. The main varieties (those with >9 samples) are shown in individual columns, while the others are pooled.

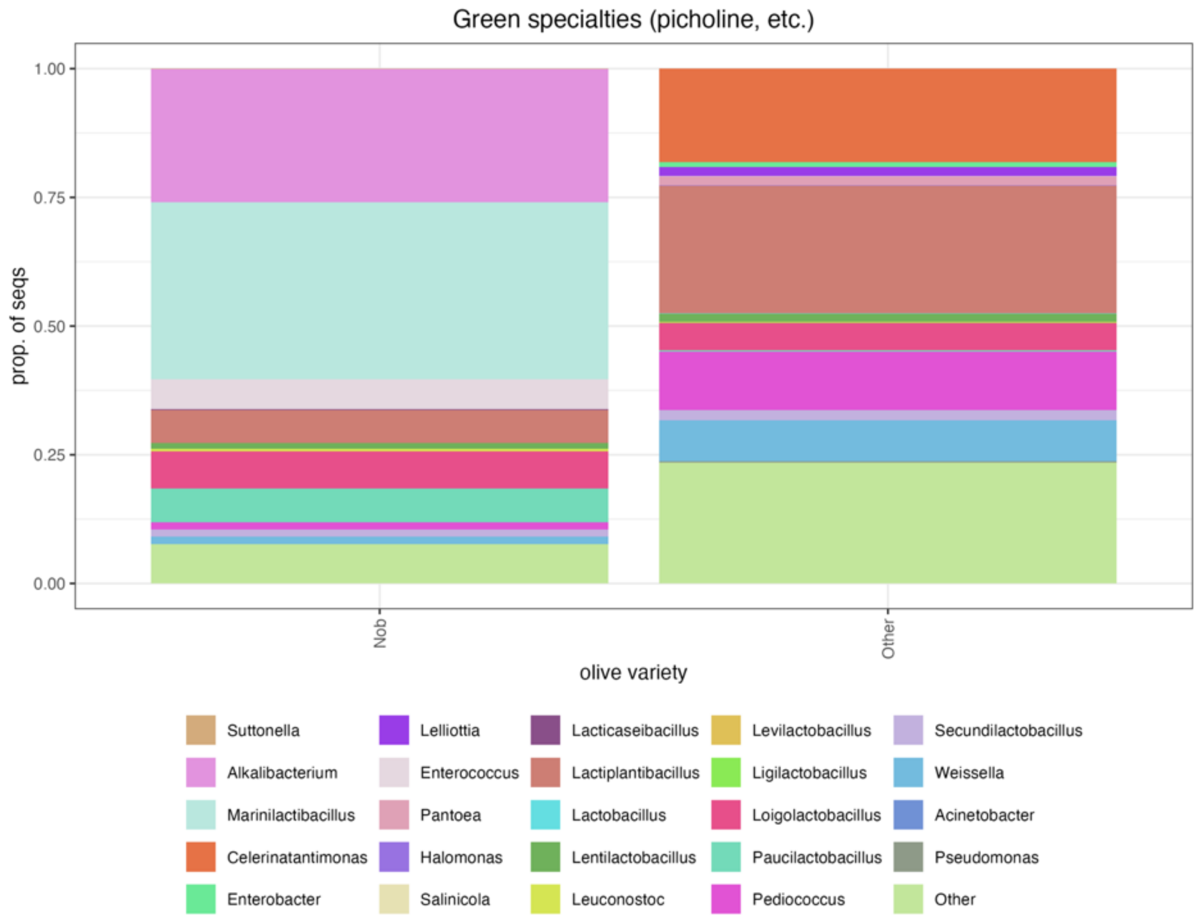


**Supplementary Figure 21**. Average abundance of the most prevalent and abundant bacterial genera in green specialties olives. The main varieties (those with >9 samples) are shown in individual columns, while the others are pooled.

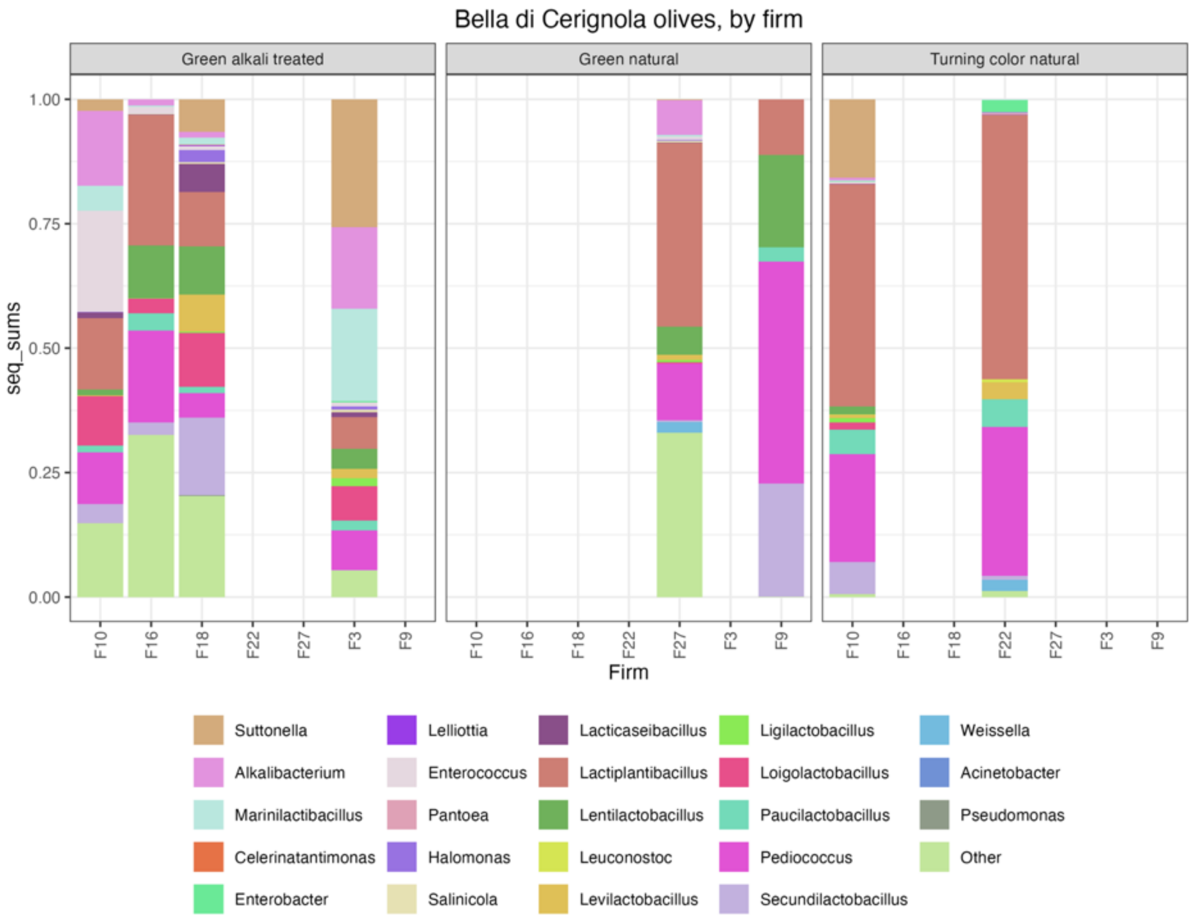


**Supplementary Figure 22**. Average abundance of the most prevalent and abundant bacterial genera in table olives produced with the Bella di Cerignola variety, by producing firm and type.

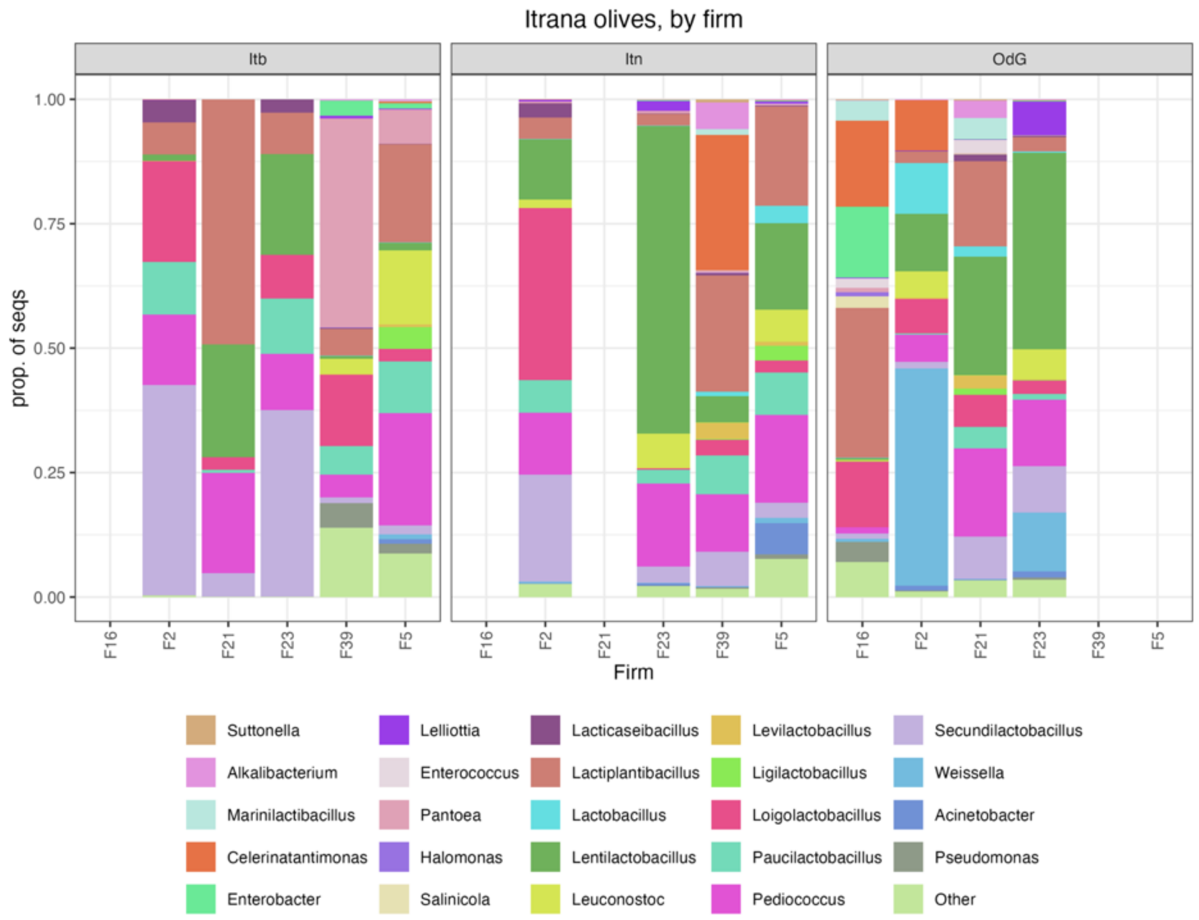


**Supplementary Figure 23**. Average abundance of the most prevalent and abundant bacterial genera in table olives produced with the Itrana variety, by producing firm and type (Itb Itrana bianca, green natural olives; Itn Itrana nera non-PDO black natural olives; OdG Oliva di Gaeta, PDO black natural olives).

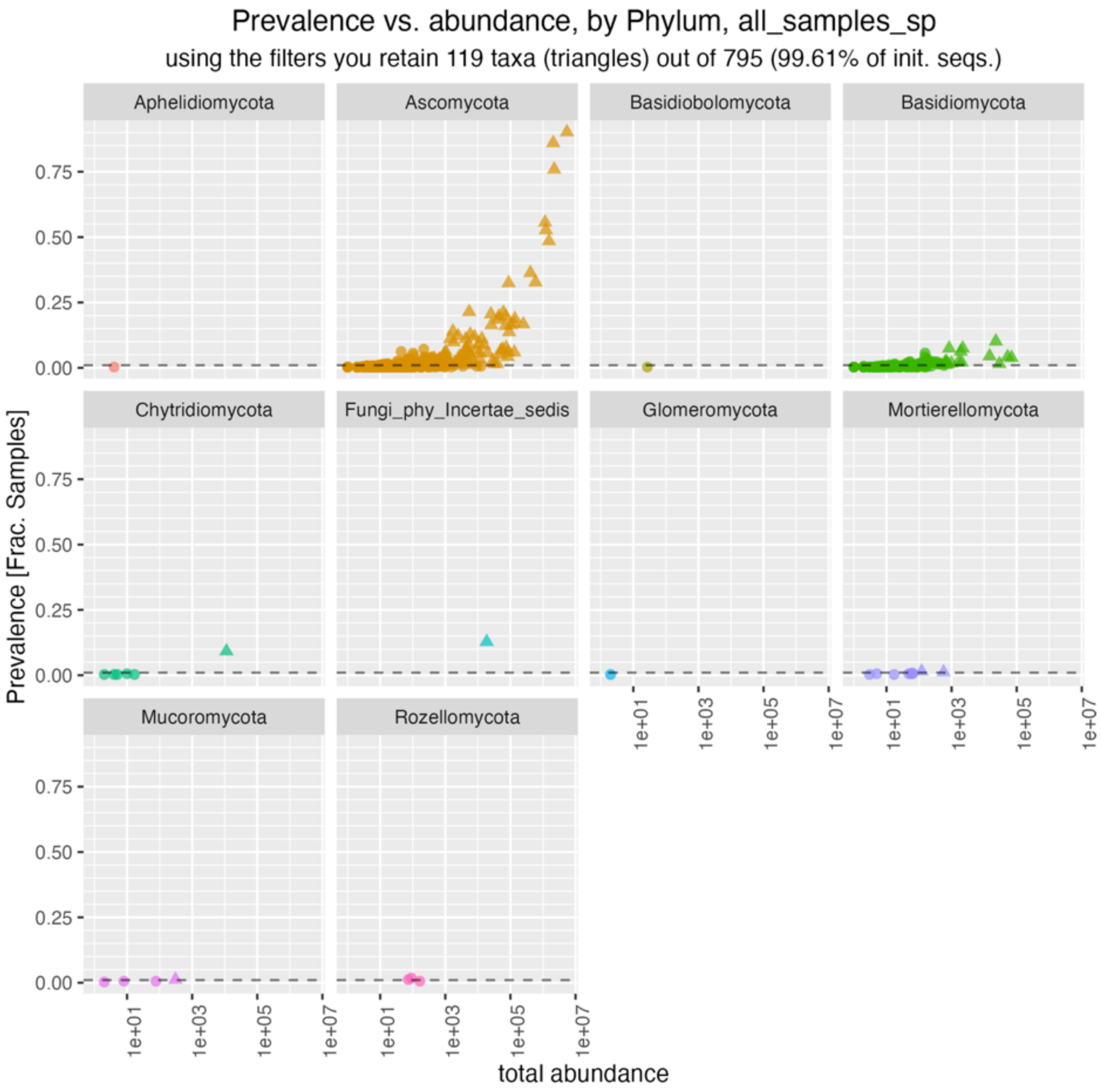


**Supplementary Figure 24**. Prevalence and abundance of fungal taxa in this study; only taxa which had a maximum abundance >1% and a prevalence >1% were retained.

**Supplementary Table 21**. Core fungal taxa in table olives analyzed in this study.

| label | Maximum relative abundance | Median relative abundance | Relative prevalence | Order |
|---|---|---|---|---|
| *Pichia membranifaciens* | 1.00 | 0.17 | 0.90 | Saccharomycetales |
| *Pichia manshurica* | 0.93 | 0.02 | 0.86 | Saccharomycetales |
| *Candida boidinii* | 0.98 | 0.00 | 0.76 | Saccharomycetales |
| *Dekkera custersiana* | 0.99 | 0.00 | 0.49 | Saccharomycetales |
| *Wickerhamomyces anomalus* | 1.00 | 0.00 | 0.56 | Saccharomycetales |
| *Saccharomyces bayanus* | 0.96 | 0.00 | 0.53 | Saccharomycetales |
| *Pichia kudriavzevii* | 0.99 | 0.00 | 0.33 | Saccharomycetales |
| *Zygotorulaspora mrakii* | 0.98 | 0.00 | 0.36 | Saccharomycetales |
| *Penicillium roqueforti* | 0.50 | 0.00 | 0.32 | Eurotiales |
| *Kluyveromyces lactis* | 1.00 | 0.00 | 0.17 | Saccharomycetales |
| *Schwanniomyces etchellsii* | 0.81 | 0.00 | 0.21 | Saccharomycetales |
| *Candida diddensiae* | 0.92 | 0.00 | 0.19 | Saccharomycetales |
| *Aureobasidium pullulans* | 0.38 | 0.00 | 0.20 | Dothideales |
| *Yamadazyma paraaseri* | 0.02 | 0.00 | 0.21 | Saccharomycetales |
| *Citeromyces nyonsensis* | 0.21 | 0.00 | 0.21 | Saccharomycetales |
| *Debaryomyces prosopidis* | 0.16 | 0.00 | 0.20 | Saccharomycetales |
| *Starmerella* | 0.56 | 0.00 | 0.17 | Saccharomycetales |
| *Cladosporium herbarum* | 0.24 | 0.00 | 0.18 | Cladosporiales |
| *Geotrichum candidum* | 0.83 | 0.00 | 0.16 | Saccharomycetales |
| *Starmerella apicola* | 0.44 | 0.00 | 0.16 | Saccharomycetales |
| *Nakazawaea molendinolei* | 0.13 | 0.00 | 0.16 | Saccharomycetales |

| label | Maximum relative abundance | Median relative abundance | Relative prevalence | Order |
|---|---|---|---|---|
| *Candida norvegica* | 0.99 | 0.00 | 0.14 | Saccharomycetales |
| *Penicillium psychrosexuale* | 0.01 | 0.00 | 0.14 | Eurotiales |
| *Fungi_gen_Incertae_sedis* | 0.34 | 0.00 | 0.13 | Fungi_ord_Incertae_sedis |
| *Meyerozyma smithsonii* | 0.06 | 0.00 | 0.13 | Saccharomycetales |
| *Wickerhamiella pararugosa* | 0.04 | 0.00 | 0.12 | Saccharomycetales |
| *Pichia* | 0.09 | 0.00 | 0.11 | Saccharomycetales |
| *Penicillium* | 0.05 | 0.00 | 0.11 | Eurotiales |
| *Vishniacozyma victoriae* | 0.24 | 0.00 | 0.10 | Tremellales |

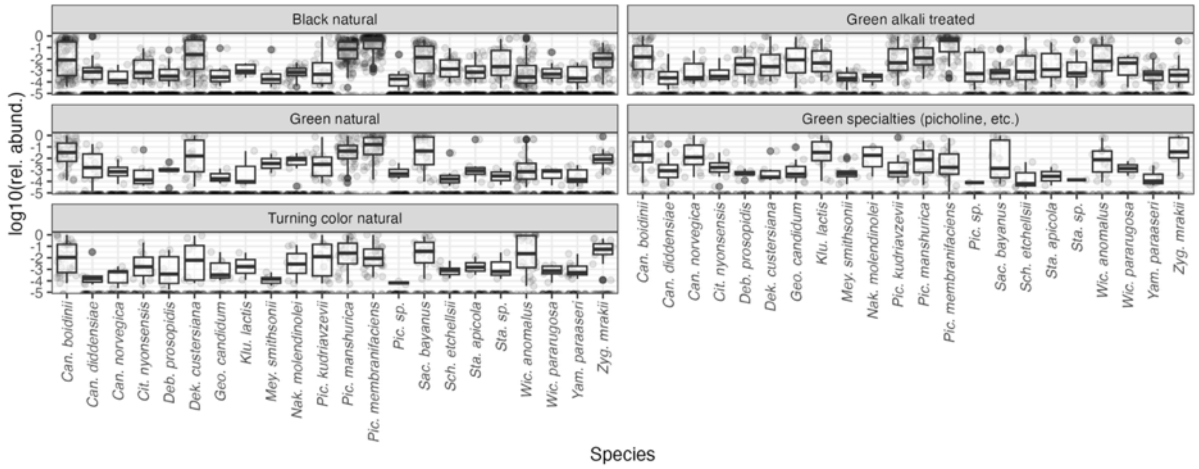


**Supplementary Figure 25**. Distribution of the abundance of the most prevalent fungal species in the main groups of table olives.

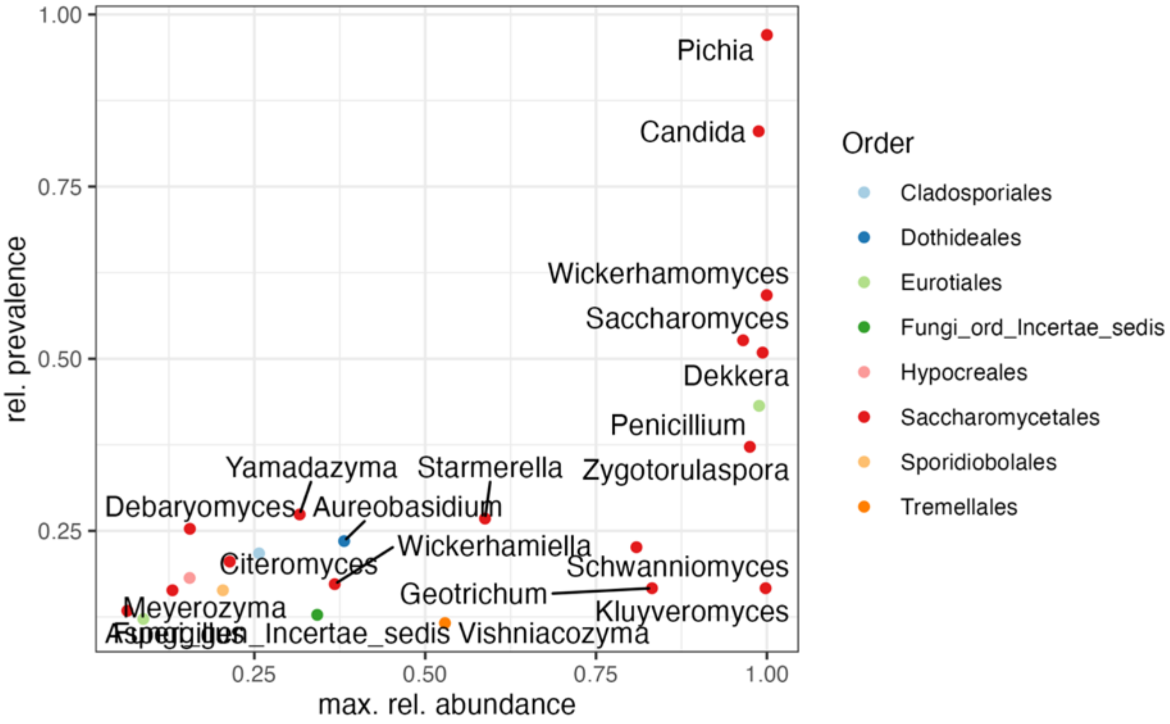


**Supplementary Figure 26**. Average prevalence and maximum relative abundance of core fungal genera in table olives.

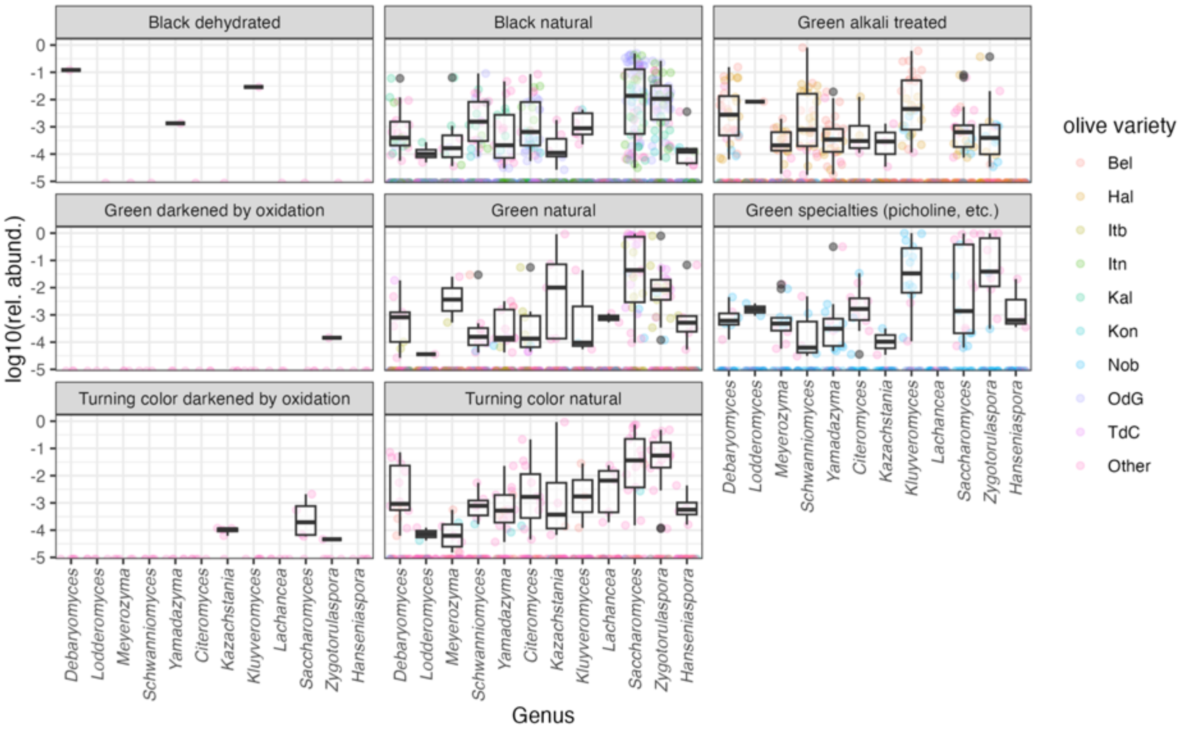


**Supplementary Figure 27.** Distribution of genera belonging to families *Saccharomycetaceae, "Saccharomycodaceae,* and *Debaryomycetaceae* in table olives used in this study. The main varieties are shown (see Supplementary Table 2 for abbreviations).

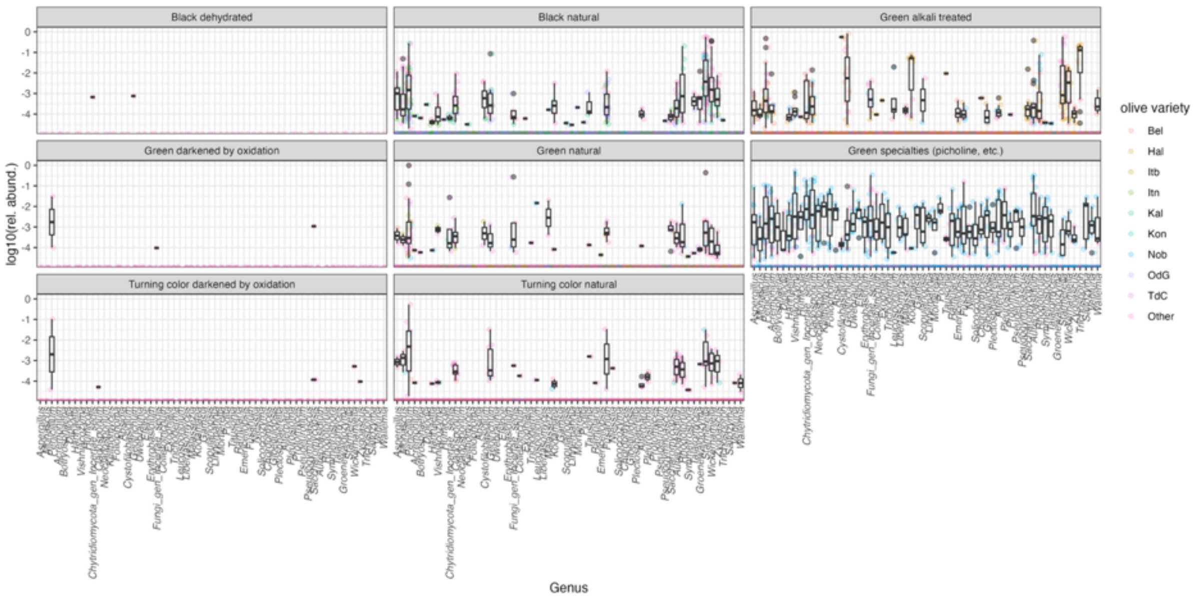


**Supplementary Figure 28.** Distribution other fungal genera in table olives used in this study. The main varieties are shown (see Supplementary Table 2 for abbreviations).

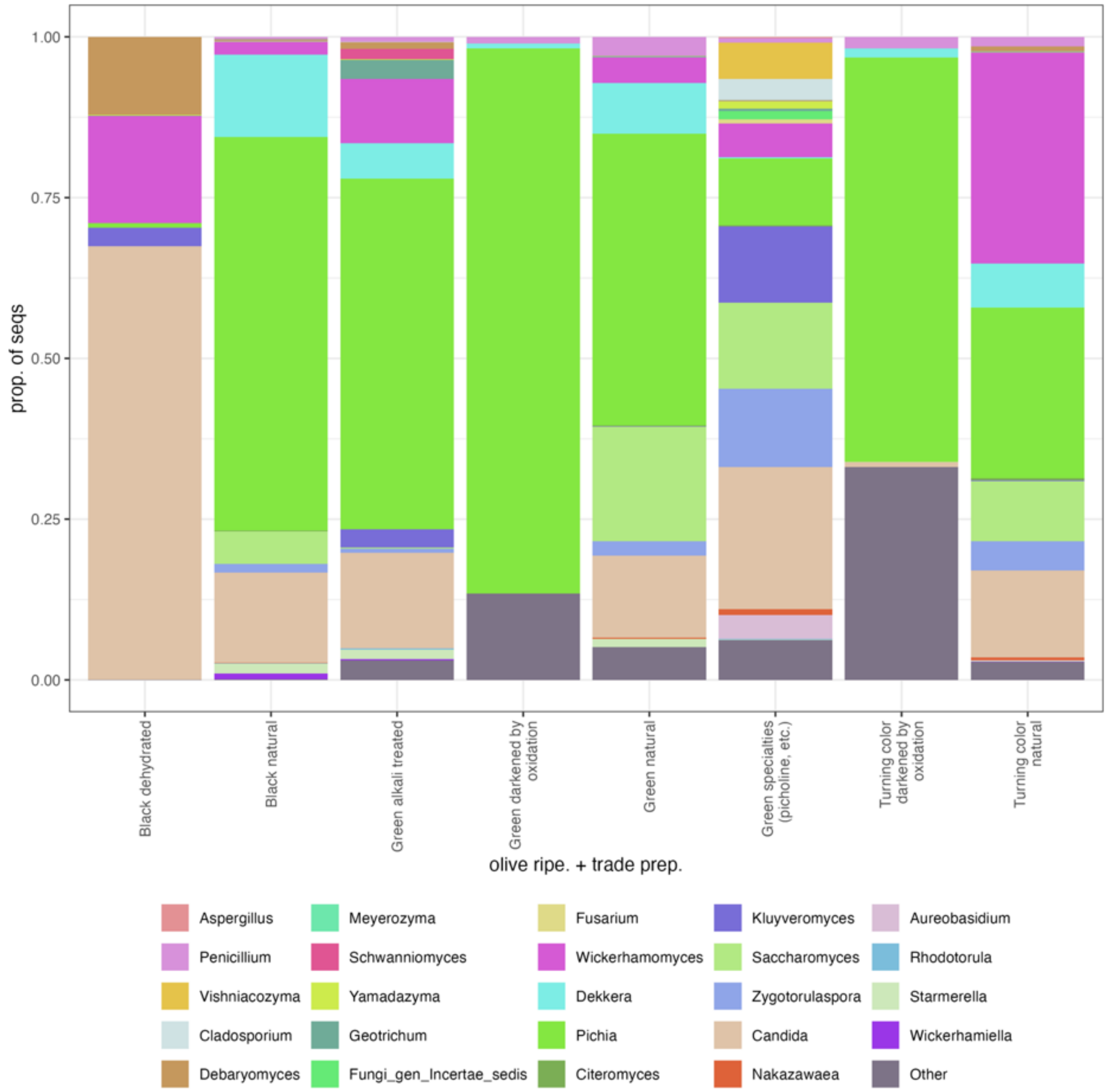


**Supplementary Figure 29**. Relative abundance of the top fungal genera in table olives used in this study. All other genera are pooled in the category “Other”.

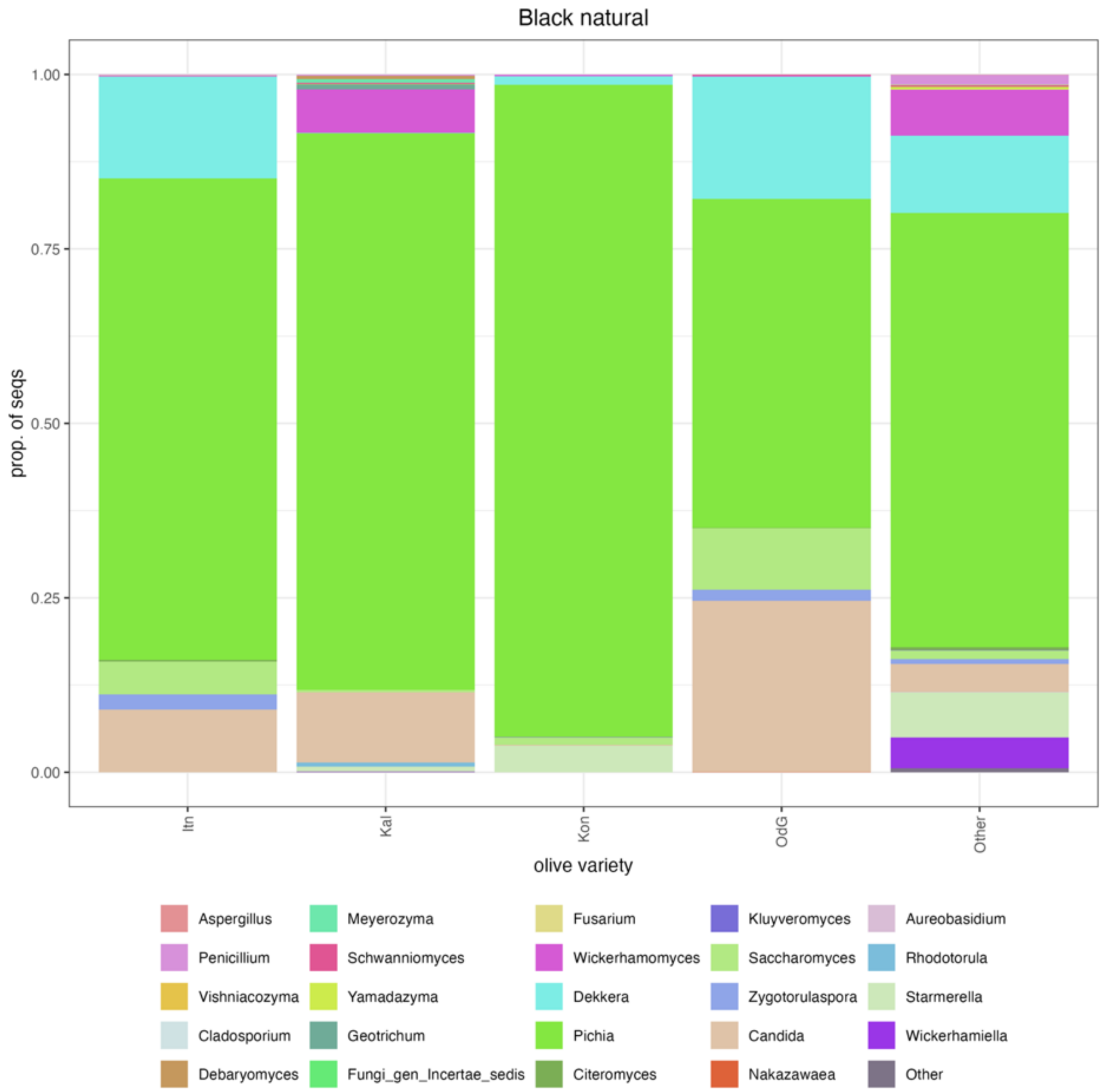


**Supplementary Figure 30.** Relative abundance of the top fungal genera in black, naturally fermented table olives used in this study. All other genera are pooled in the category "Other".

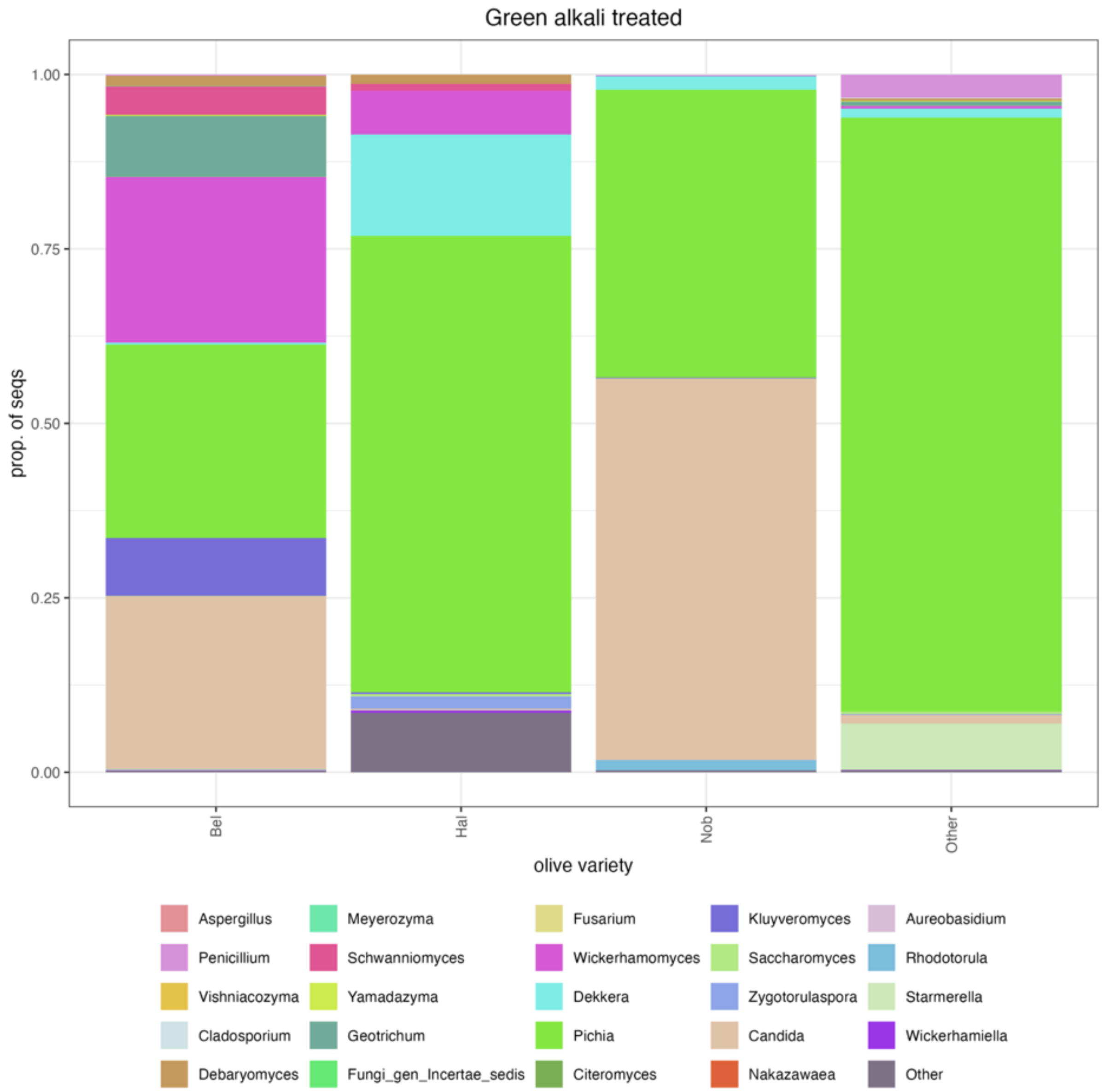


**Supplementary Figure 31.** Relative abundance of the top fungal genera in green, alkali treated table olives used in this study. All other genera are pooled in the category "Other".

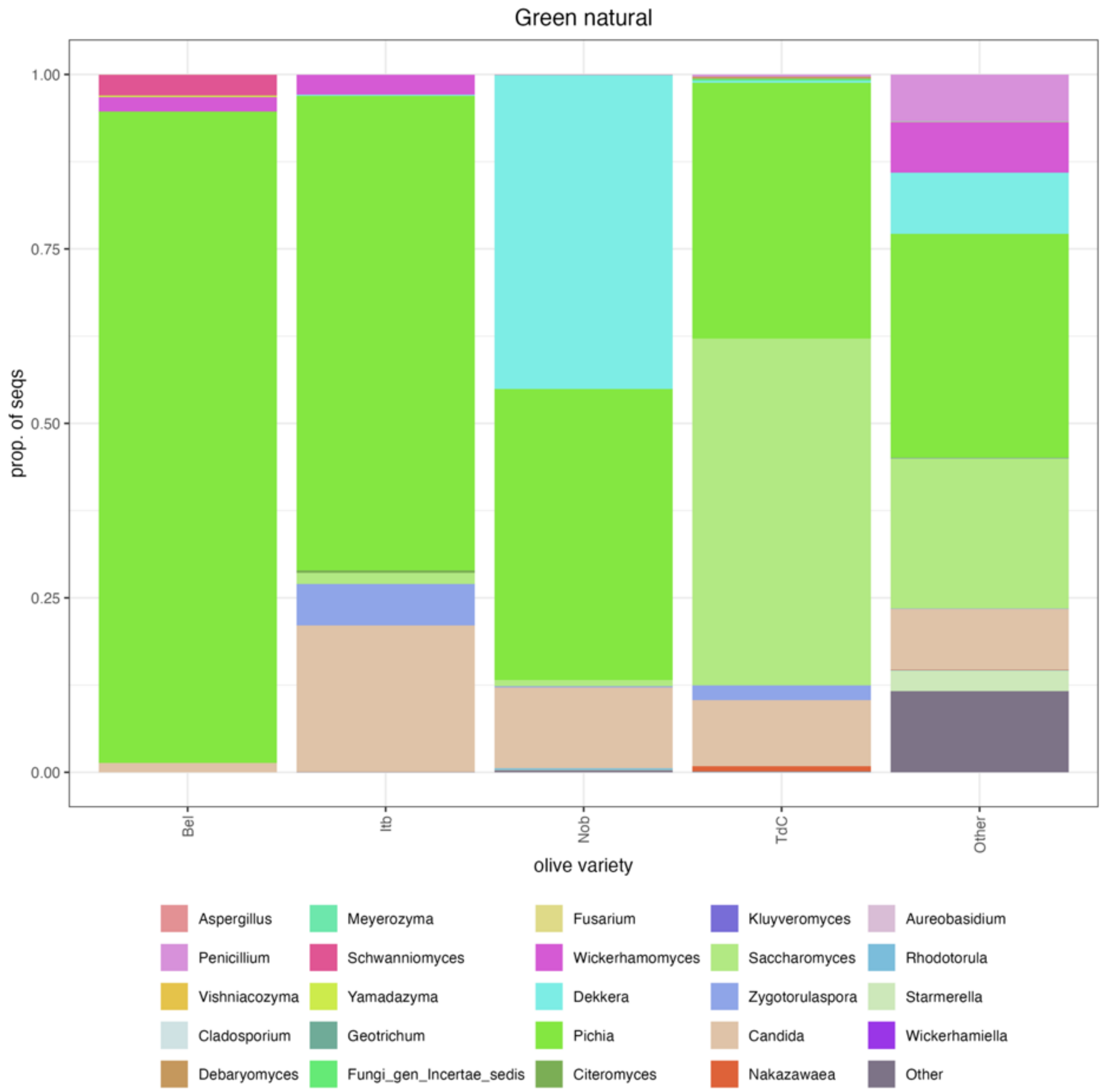


**Supplementary Figure 32.** Relative abundance of the top fungal genera in green, naturally fermented table olives used in this study. All other genera are pooled in the category "Other".

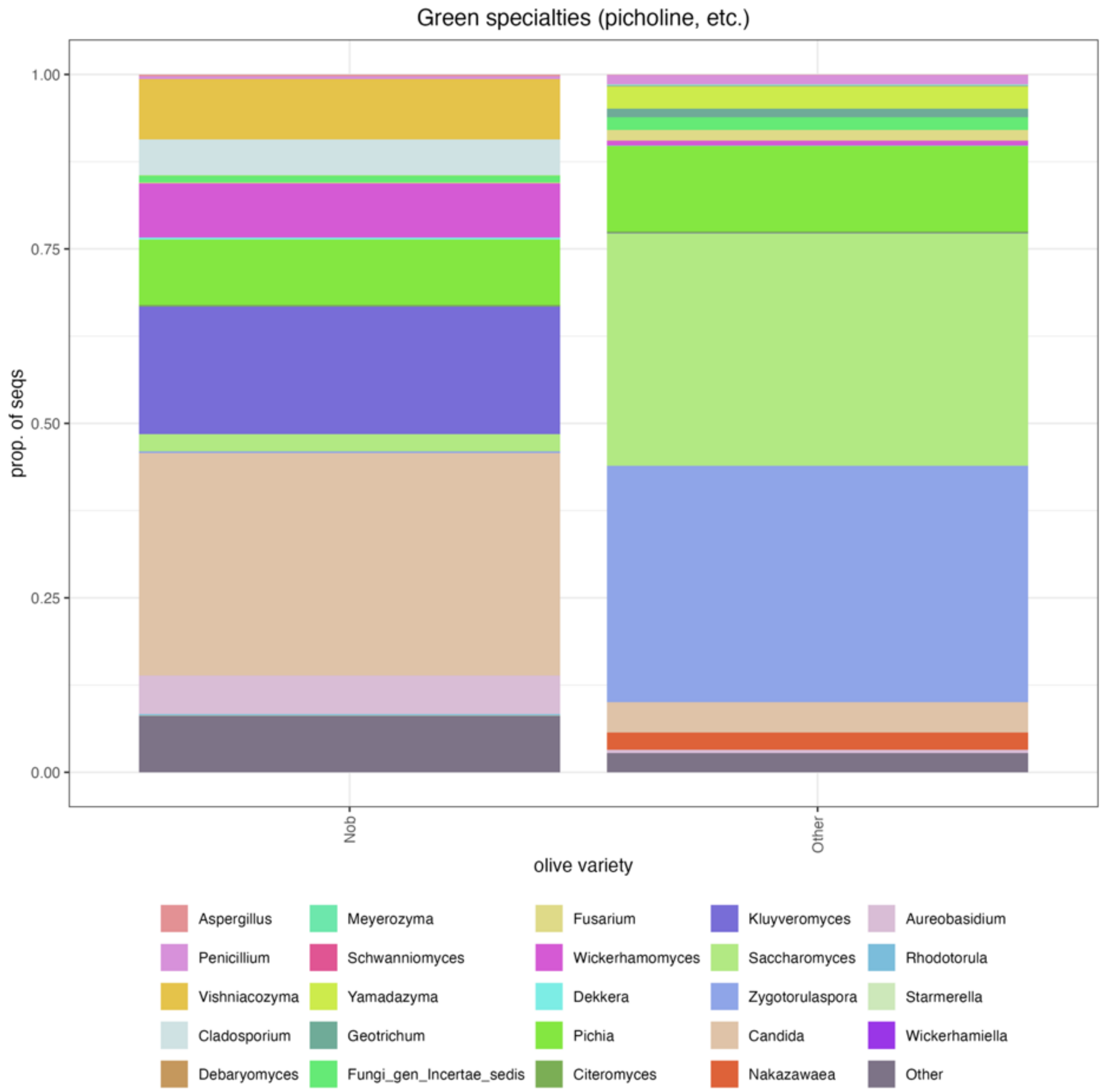


**Supplementary Figure 33.** Relative abundance of the top fungal genera in green specialties table olives used in this study. All other genera are pooled in the category “Other”.

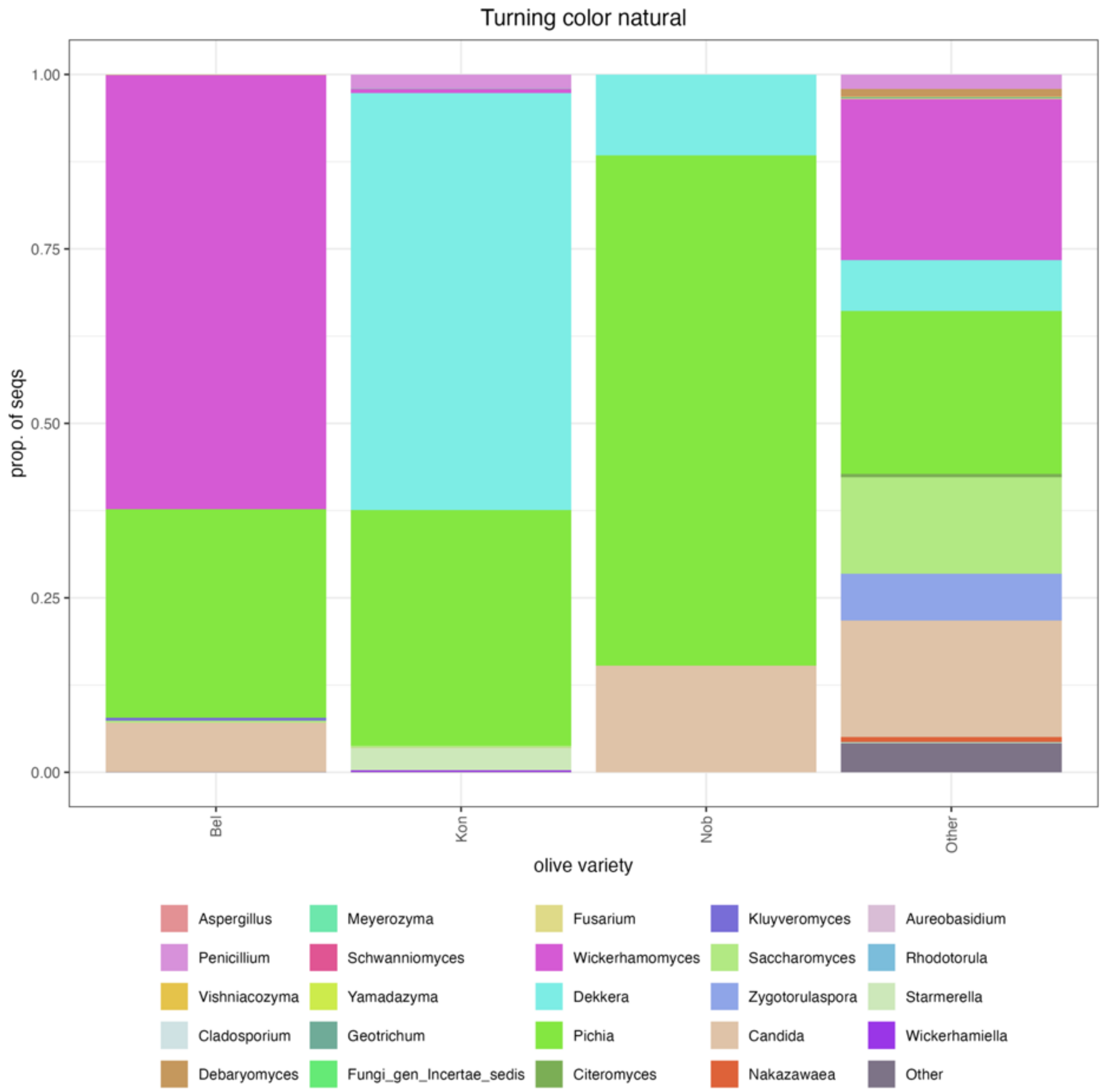


**Supplementary Figure 34.** Relative abundance of the top fungal genera in turning color, naturally fermented table olives used in this study. All other genera are pooled in the category “Other”.

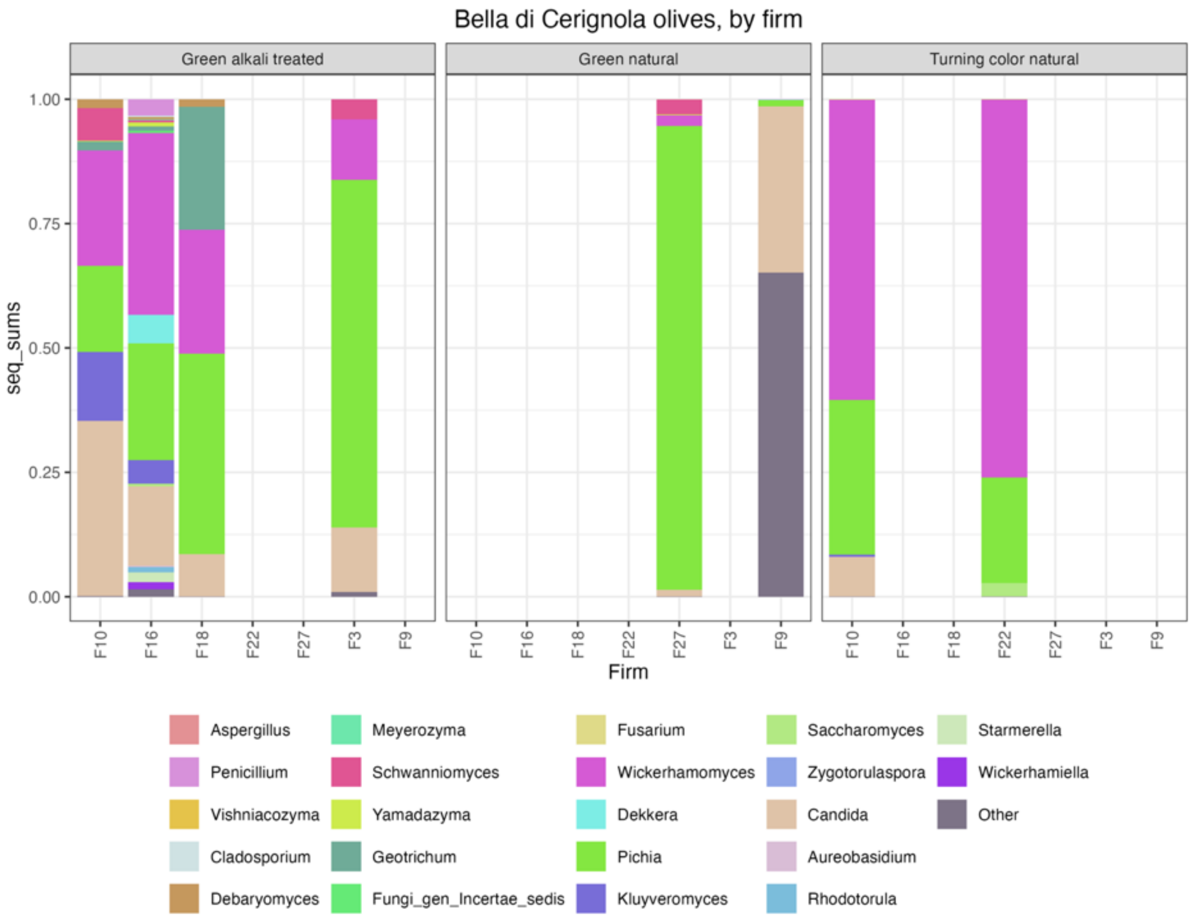


**Supplementary Figure 35.** Relative abundance of the top fungal genera in Bella di Cerignola table olives used in this study by different firms. All other genera are pooled in the category "Other".

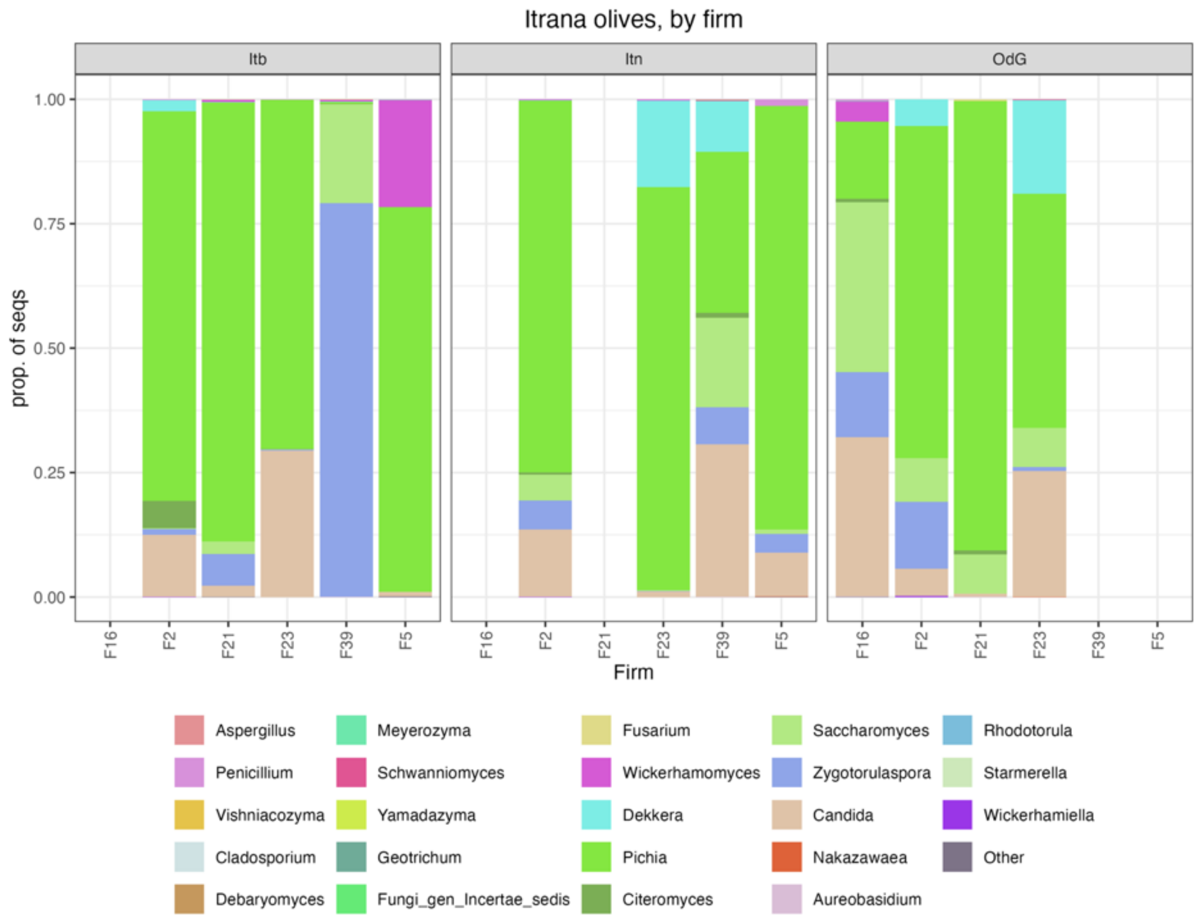


**Supplementary Figure 36.** Relative abundance of the top fungal genera in Itrana table olives used in this study by different firms. All other genera are pooled in the category “Other”.

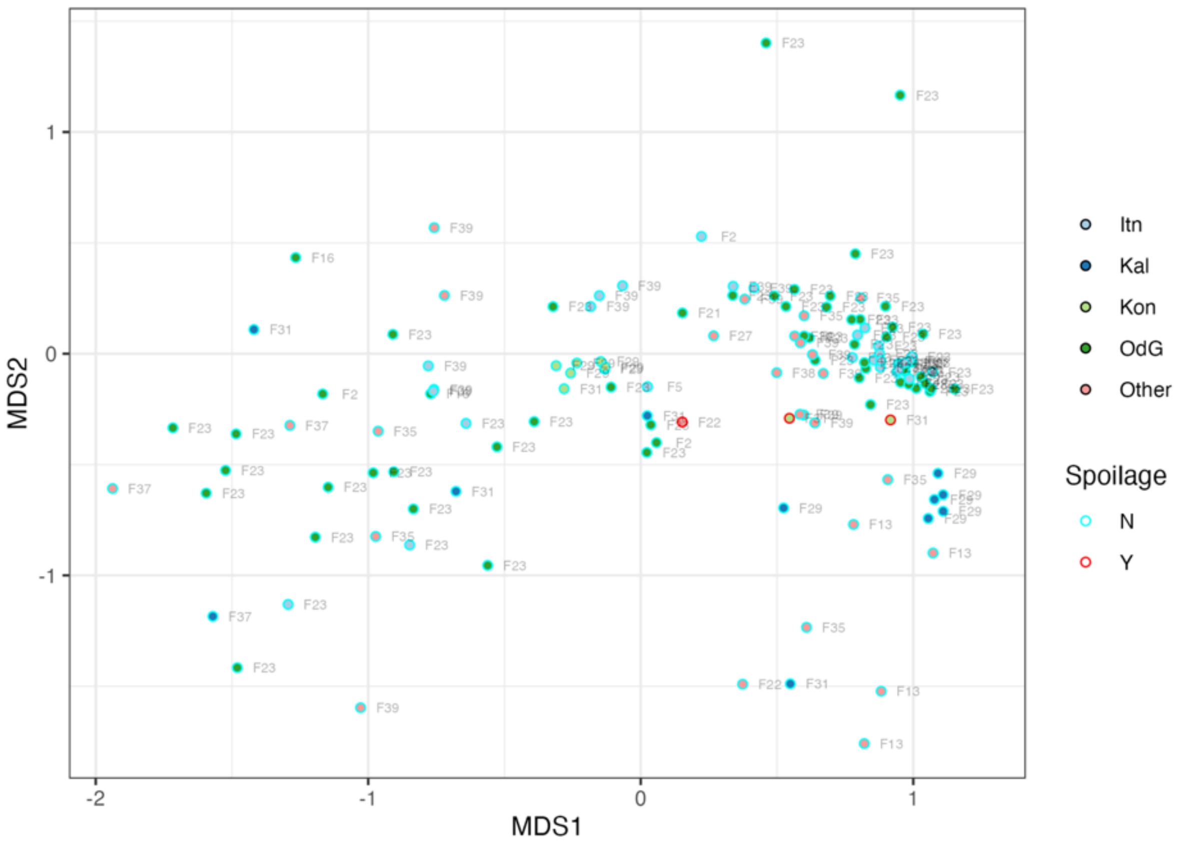


**Supplementary Figure 37.** Configuration plot for the first two dimensions of a non-monotonic Multidimensional Scaling of the Bray-Curtis distance matrix of the composition of bacterial communities of black natural table olives. Fx indicates the producing firm.

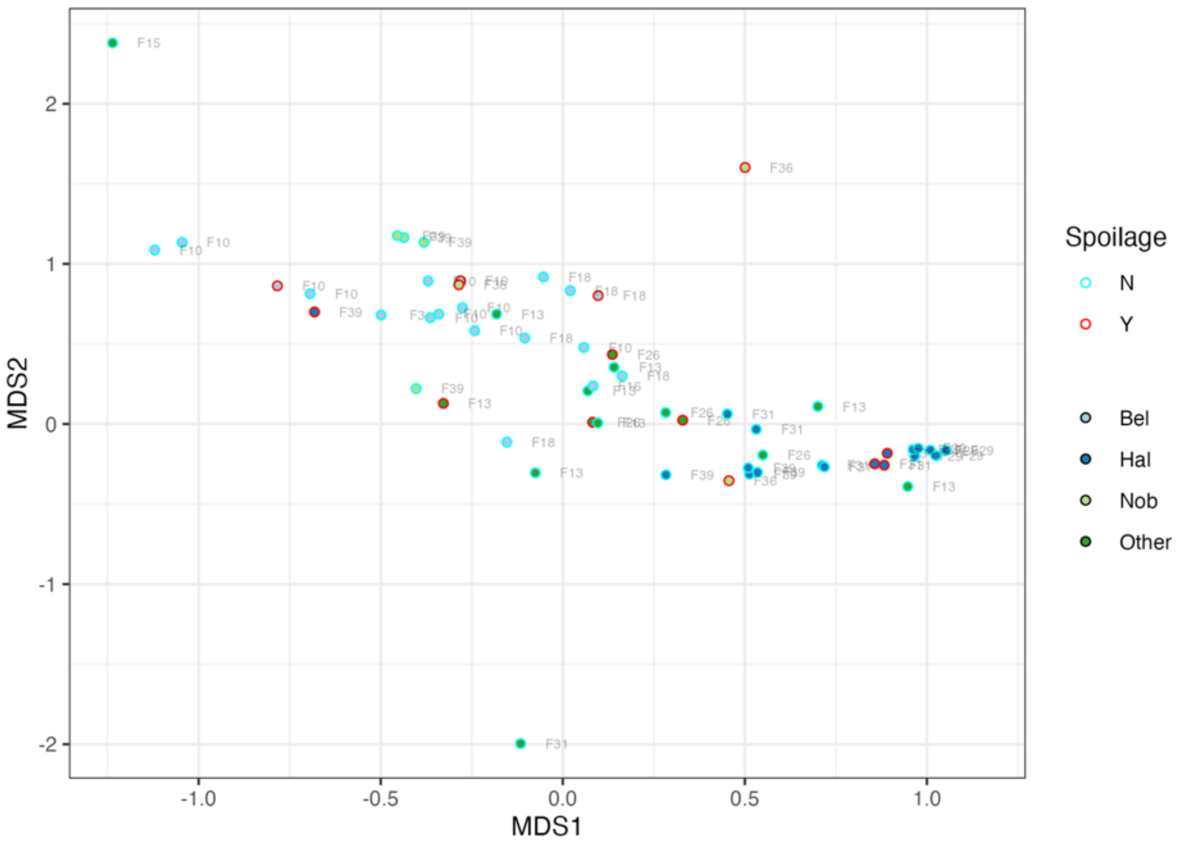


**Supplementary Figure 38**. Configuration plot for the first two dimensions of a non-monotonic Multidimensional Scaling of the Bray-Curtis distance matrix of the composition of bacterial communities of green alkali treated table olives. Fx indicates the producing firm.

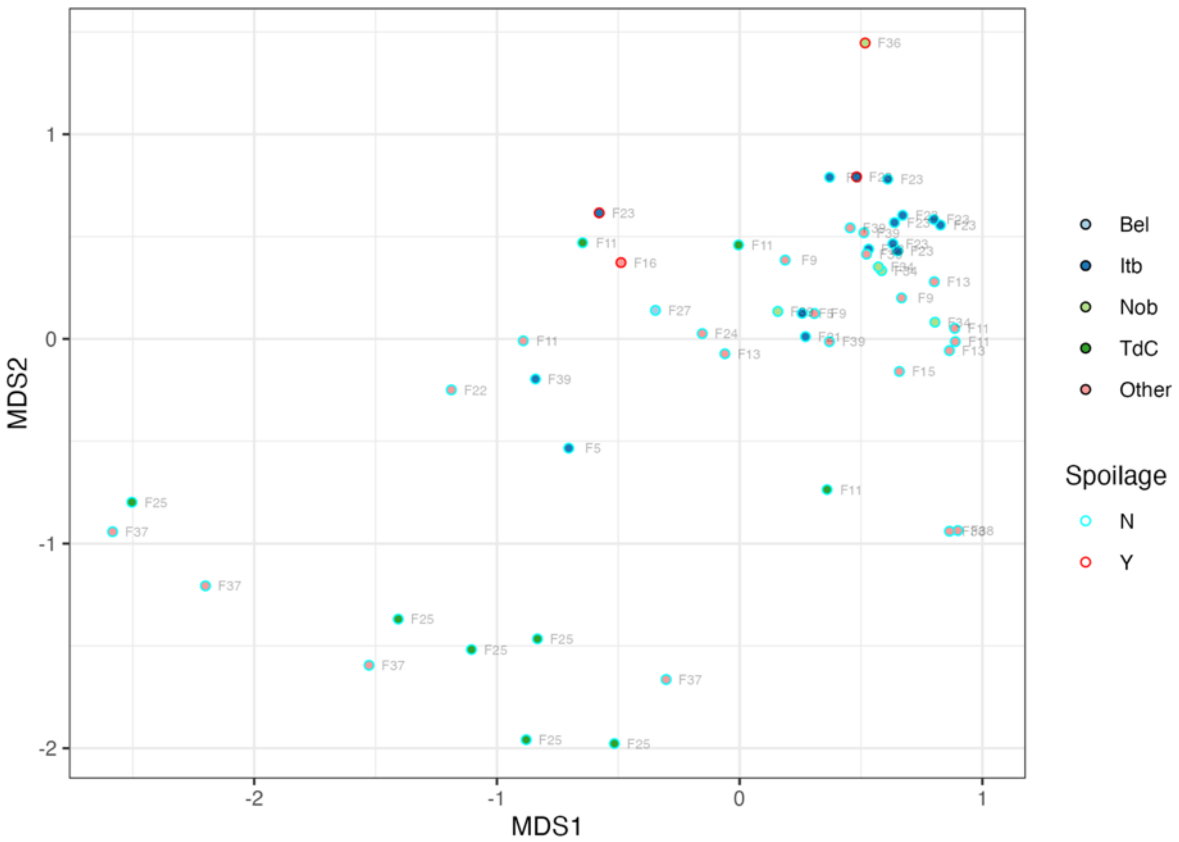


**Supplementary Figure 39**. Configuration plot for the first two dimensions of a non-monotonic Multidimensional Scaling of the Bray-Curtis distance matrix of the composition of bacterial communities of green natural table olives. Fx indicates the producing firm.

**Supplementary Table 22.** Results of PERMANOVA used to test the hypothesis that variety did not affect the composition of bacterial communities of table olives within a given ripeness and trade preparation group.

| Ripening stage + trade preparation | samples | varieties | F | $R^2$ | Prob |
|---|---|---|---|---|---|
| Black natural | 122 | 5 | 5.758 | 0.163 | 0.001 |
| Green alkali treated | 52 | 4 | 8.148 | 0.337 | 0.001 |
| Green natural* | 31 | 3 | 4.064 | 0.225 | 0.002 |
| Turning color natural | 23 | 3 | 1.178 | 0.105 | 0.307 |

* results of the betadispers() test were significant.

**Supplementary Table 23.** Results of PERMANOVA used to test the hypothesis that firm affected the composition of bacterial communities of table olives specific sets of olive varieties.

| Ripening stage + trade preparation | Olive variety | samples | Firms | F | $R^2$ | Prob |
|---|---|---|---|---|---|---|
| Black natural | Itrana nera | 32 | 2 | 13.128 | 0.304 | 0.001 |
| Black natural | Kalamata* | 11 | 2 | 7.807 | 0.464 | 0.005 |
| Green alkali treated | Bella di Daunia | 17 | 2 | 4.341 | 0.165 | 0.001 |
| Green alkali treated | Halkidiki | 20 | 3 | 3.445 | 0.288 | 0.001 |

* results of the betadispers() test were significant.

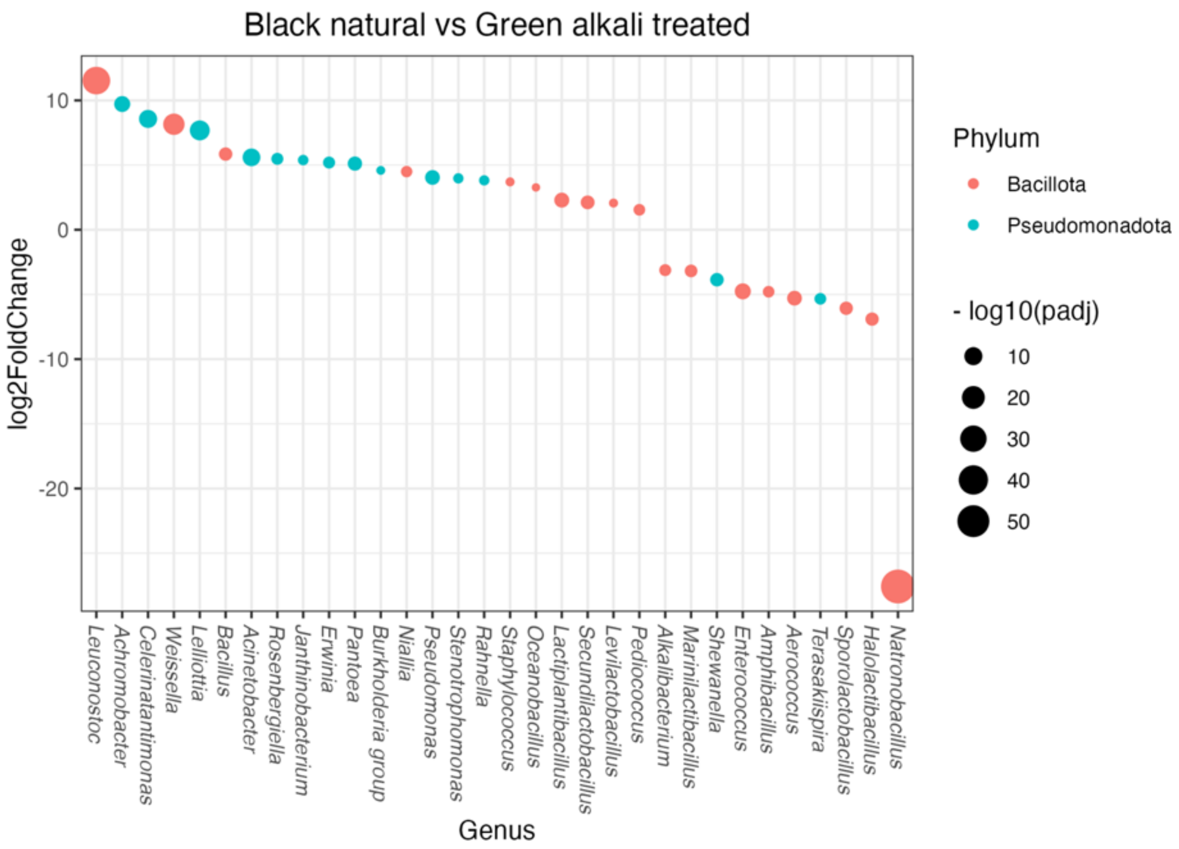


**Supplementary Figure 40**. Differentially abundant bacterial genera for the contrast Black natural vs. Green alkali treated olives, DeSeq2

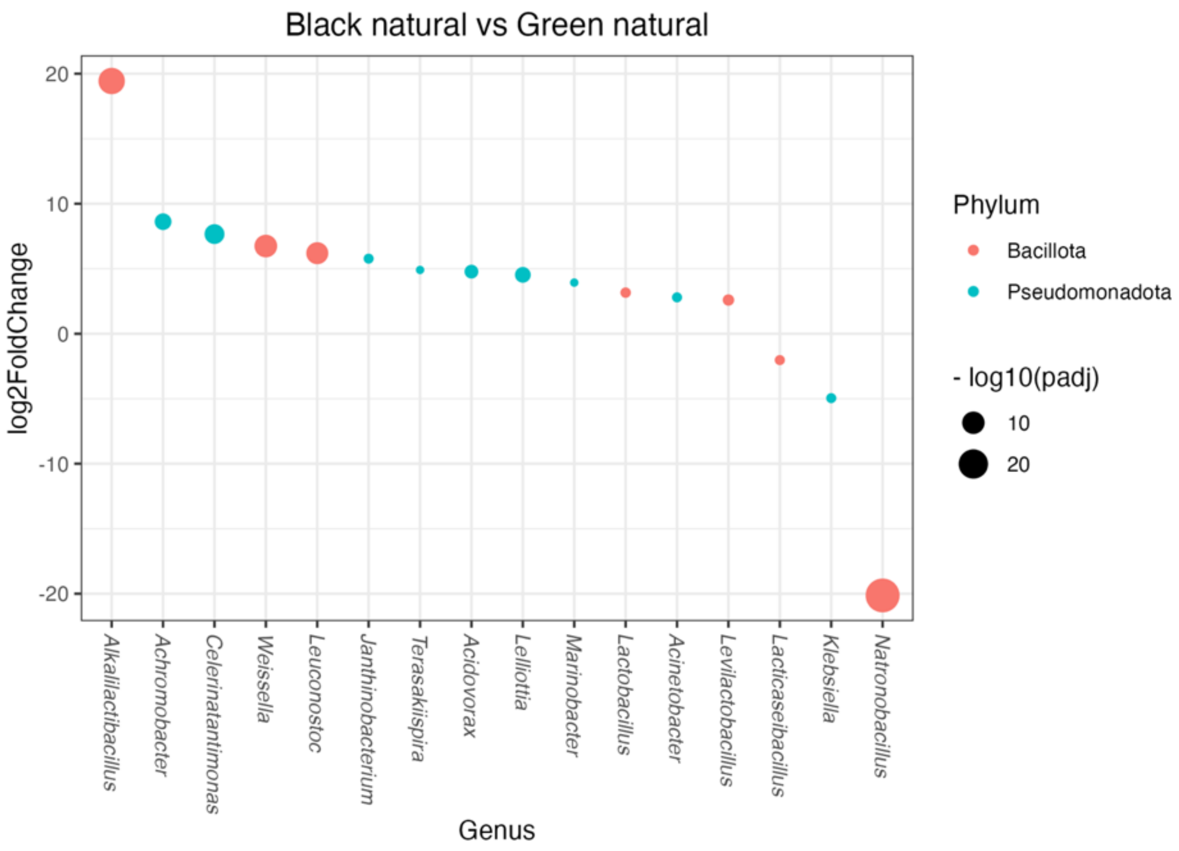


**Supplementary Figure 41.** Differentially abundant bacterial genera for the contrast Black natural vs. Green natural olives, DeSeq2

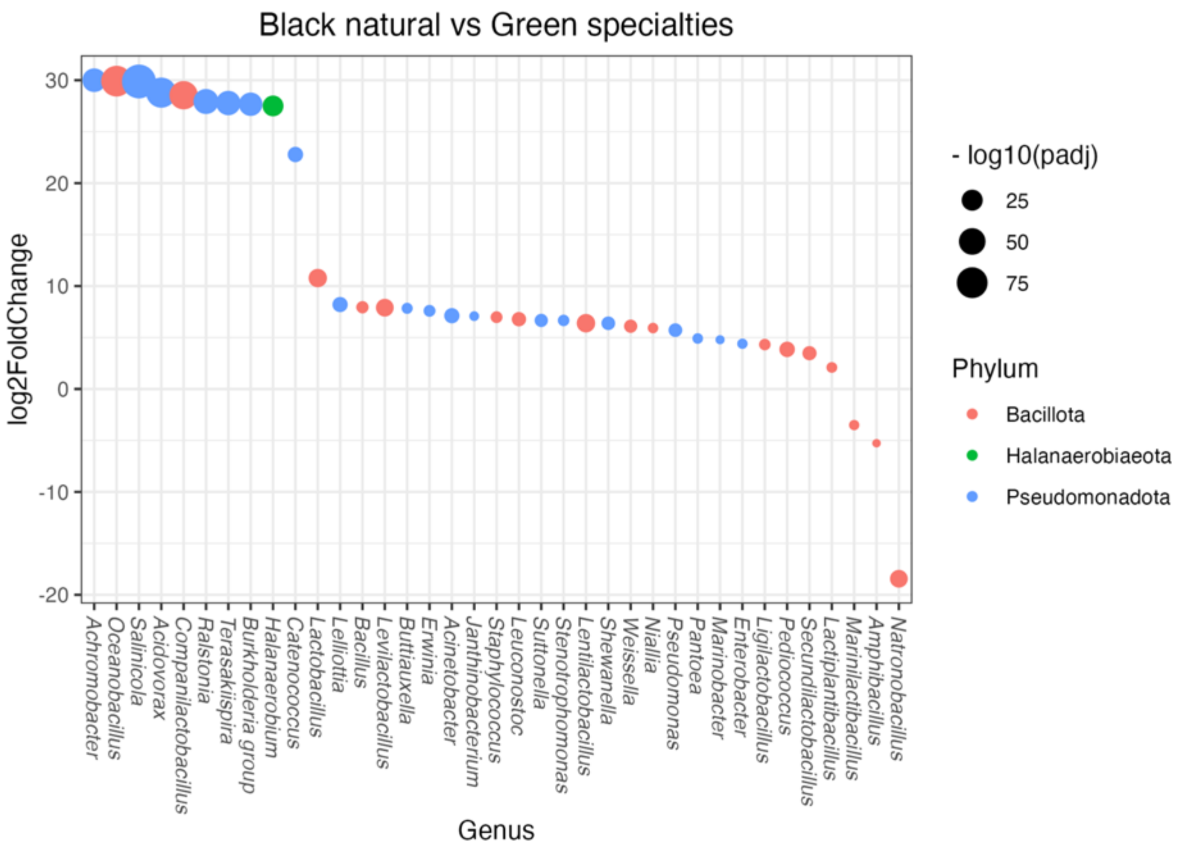


**Supplementary Figure 42.** Differentially abundant bacterial genera for the contrast Black natural vs. Green specialties, DeSeq2

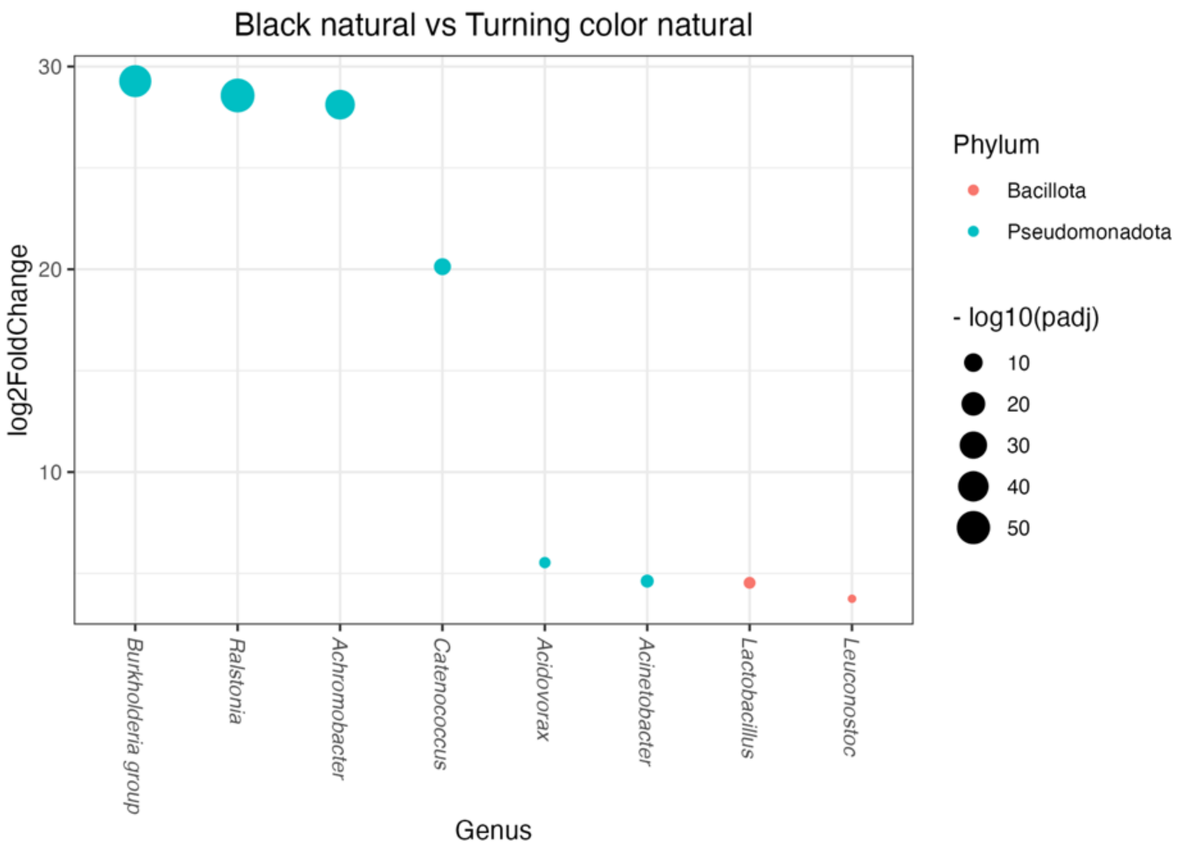


**Supplementary Figure 43.** Differentially abundant bacterial genera for the contrast Black natural vs. Turning color natural, DeSeq2

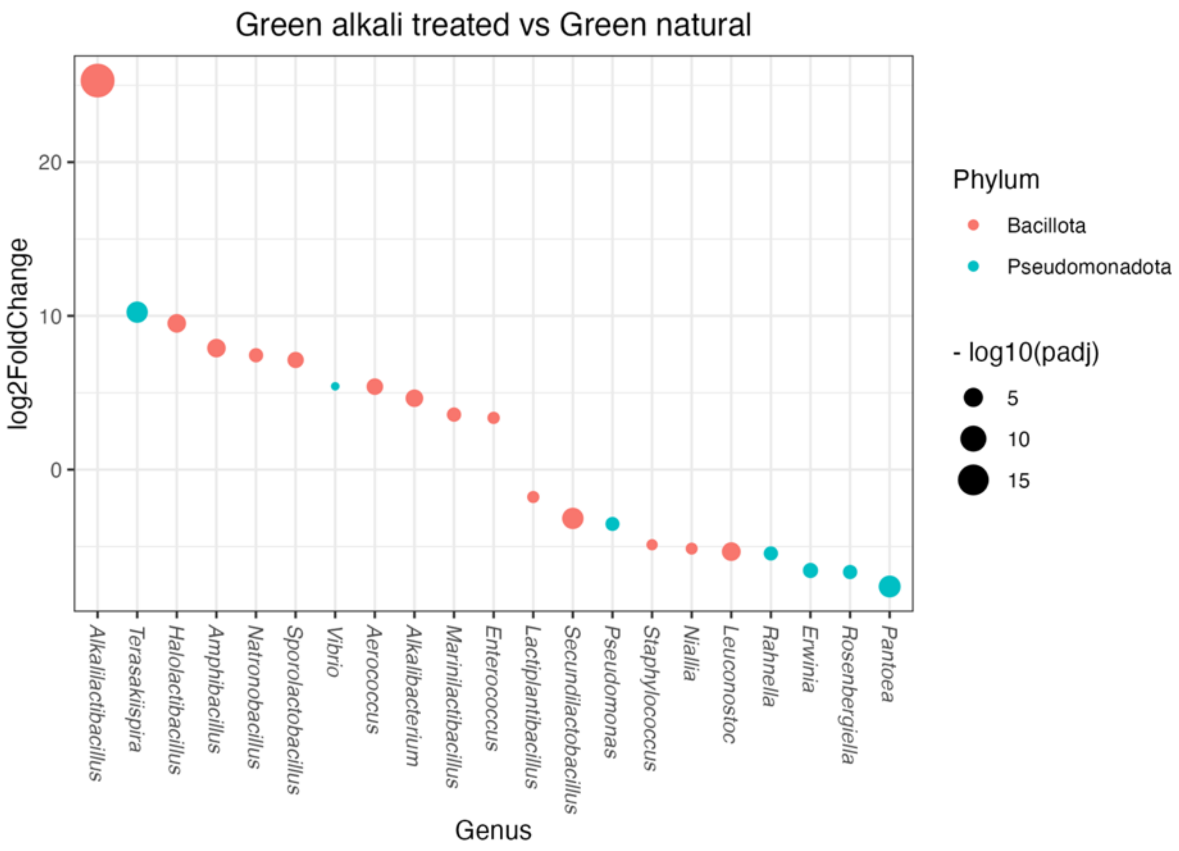


**Supplementary Figure 44.** Differentially abundant bacterial genera for the contrast Green alkali treated vs. Green natural, DeSeq2

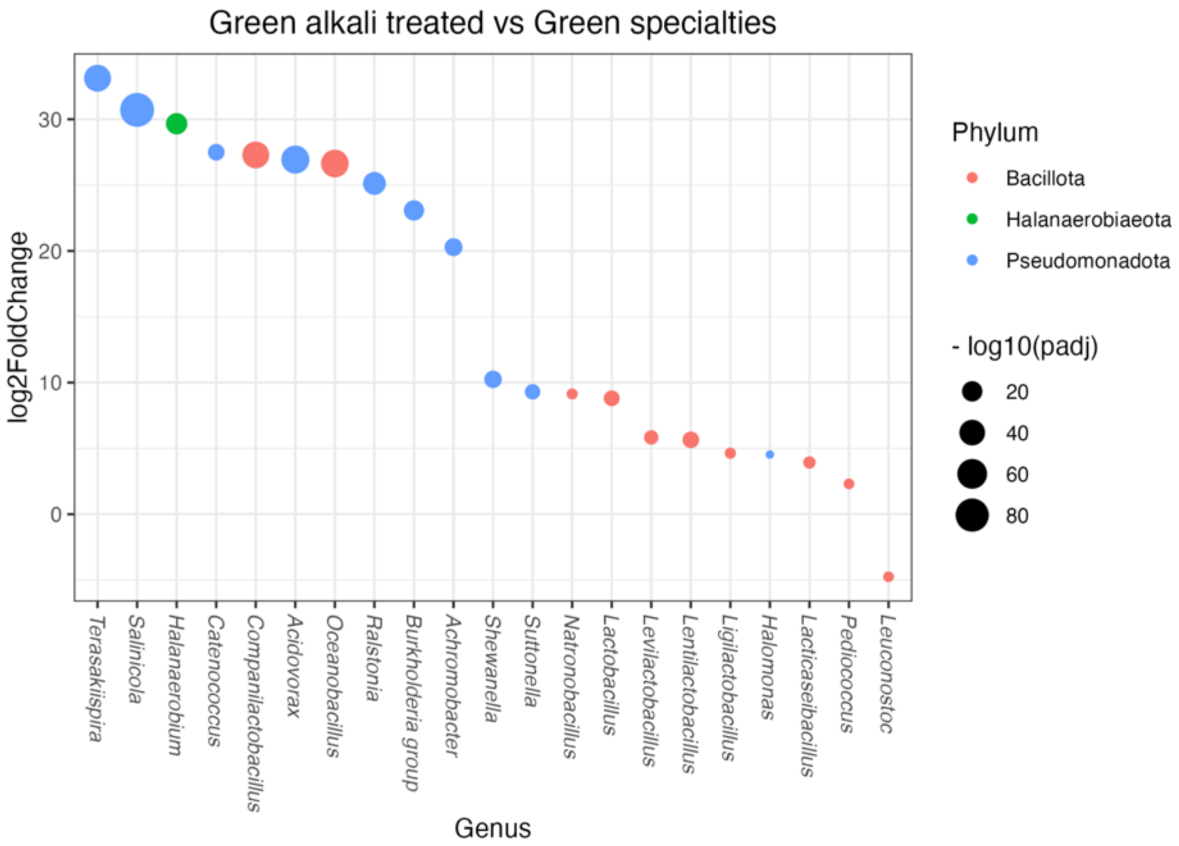


**Supplementary Figure 45**. Differentially abundant bacterial genera for the contrast Green alkali treated vs. Green specialties, DeSeq2

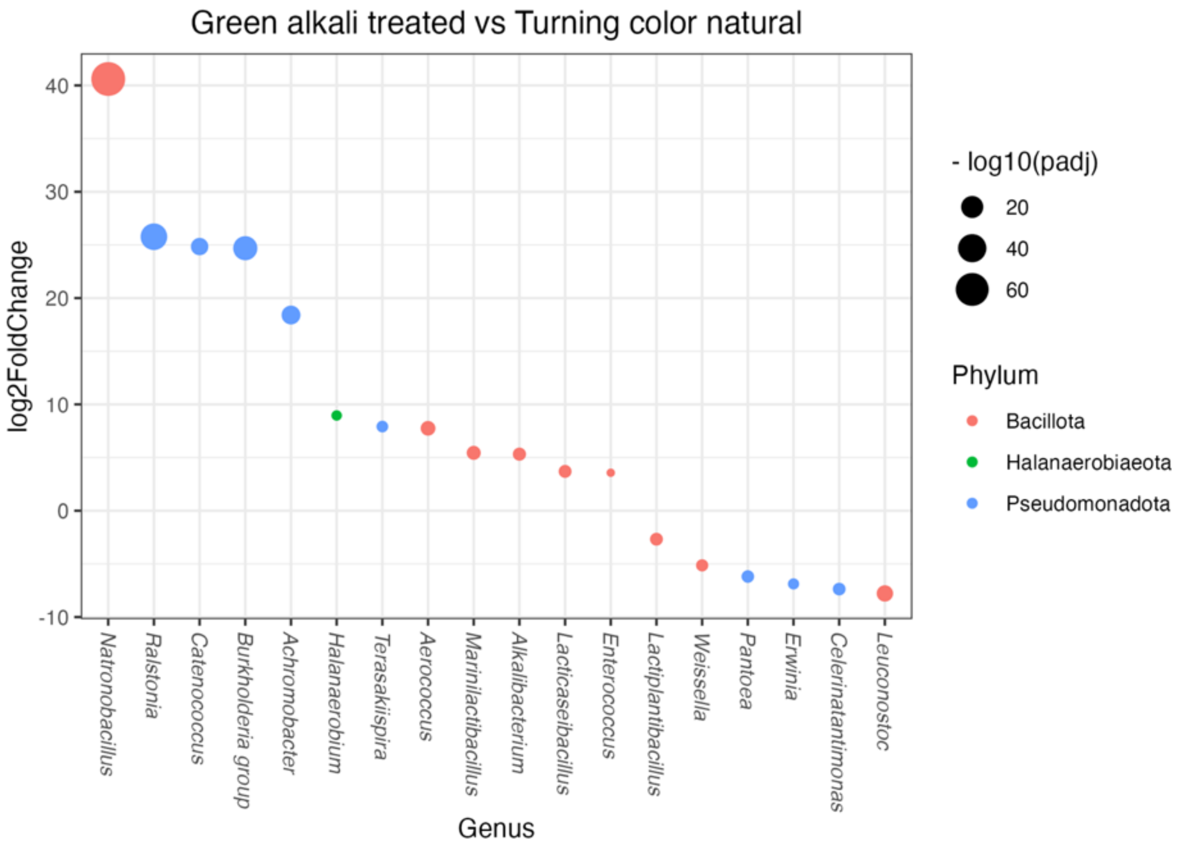


**Supplementary Figure 46.** Differentially abundant bacterial genera for the contrast Green alkali treated vs. Turning color natural, DeSeq2

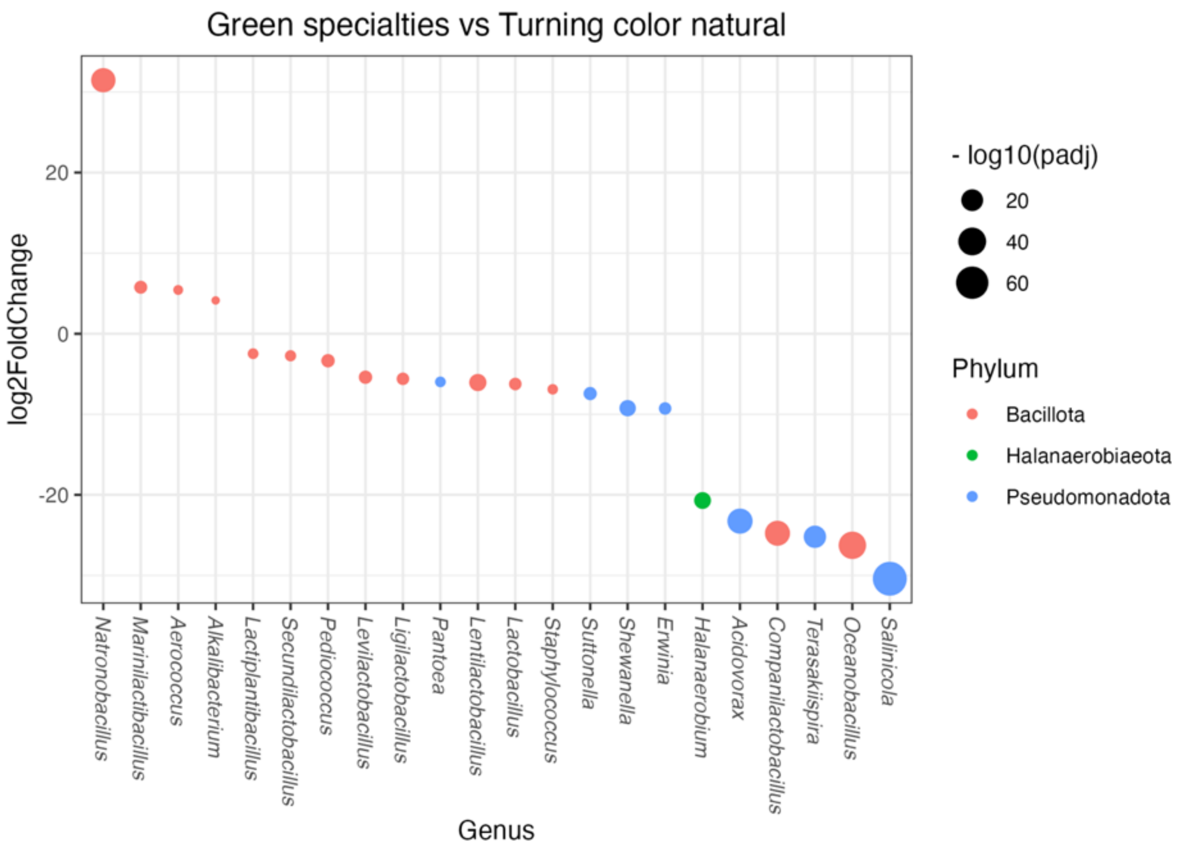


**Supplementary Figure 47.** Differentially abundant bacterial genera for the contrast Green specialties vs. Turning color natural, DeSeq2

**Supplementary table 24.** Summary results for DeSeq2 differential abundance analysis for bacterial genera.

| contrast | Genus | Family | baseMean | log2FoldChange | padj |
|---|---|---|---|---|---|
| Black_natural vs. Green_alkali_treated | *Aerococcus* | *Aerococcaceae* | 89.27 | -5.29 | 0.00 |
| Green_alkali_treated vs. Green_natural | *Aerococcus* | *Aerococcaceae* | 89.27 | 5.40 | 0.00 |
| Green_alkali_treated vs. Turning_color_natural | *Aerococcus* | *Aerococcaceae* | 89.27 | 7.75 | 0.00 |
| Green_specialties vs. Turning_color_natural | *Aerococcus* | *Aerococcaceae* | 89.27 | 5.43 | 0.01 |
| Black_natural vs. Green_alkali_treated | *Achromobacter* | *Alcaligenaceae* | 132.48 | 9.72 | 0.00 |
| Black_natural vs. Green_natural | *Achromobacter* | *Alcaligenaceae* | 132.48 | 8.62 | 0.00 |
| Black_natural vs. Green_specialties | *Achromobacter* | *Alcaligenaceae* | 132.48 | 30.00 | 0.00 |
| Black_natural vs. Turning_color_natural | *Achromobacter* | *Alcaligenaceae* | 132.48 | 28.12 | 0.00 |
| Green_alkali_treated vs. Green_specialties | *Achromobacter* | *Alcaligenaceae* | 132.48 | 20.28 | 0.00 |
| Green_alkali_treated vs. Turning_color_natural | *Achromobacter* | *Alcaligenaceae* | 132.48 | 18.41 | 0.00 |
| Black_natural vs. Green_natural | *Alkalilactibacillus* | *Bacillaceae* | 4.84 | 19.44 | 0.00 |
| Green_alkali_treated vs. Green_natural | *Alkalilactibacillus* | *Bacillaceae* | 4.84 | 25.30 | 0.00 |
| Black_natural vs. Green_alkali_treated | *Amphibacillus* | *Bacillaceae* | 19.91 | -4.79 | 0.00 |
| Black_natural vs. Green_specialties | *Amphibacillus* | *Bacillaceae* | 19.91 | -5.27 | 0.01 |
| Green_alkali_treated vs. Green_natural | *Amphibacillus* | *Bacillaceae* | 19.91 | 7.90 | 0.00 |
| Black_natural vs. Green_alkali_treated | *Bacillus* | *Bacillaceae* | 5.38 | 5.85 | 0.00 |
| Black_natural vs. Green_specialties | *Bacillus* | *Bacillaceae* | 5.38 | 7.94 | 0.00 |
| Black_natural vs. Green_alkali_treated | *Halolactibacillus* | *Bacillaceae* | 29.81 | -6.91 | 0.00 |
| Green_alkali_treated vs. Green_natural | *Halolactibacillus* | *Bacillaceae* | 29.81 | 9.51 | 0.00 |
| Black_natural vs. Green_alkali_treated | *Natronobacillus* | *Bacillaceae* | 43.73 | -27.57 | 0.00 |
| Black_natural vs. Green_natural | *Natronobacillus* | *Bacillaceae* | 43.73 | -20.13 | 0.00 |
| Black_natural vs. Green_specialties | *Natronobacillus* | *Bacillaceae* | 43.73 | -18.44 | 0.00 |
| Green_alkali_treated vs. Green_natural | *Natronobacillus* | *Bacillaceae* | 43.73 | 7.44 | 0.00 |
| Green_alkali_treated vs. Green_specialties | *Natronobacillus* | *Bacillaceae* | 43.73 | 9.14 | 0.00 |

| contrast | Genus | Family | baseMean | log2FoldChange | padj |
|---|---|---|---|---|---|
| Green_alkali_treated vs. Turning_color_natural | *Natronobacillus* | *Bacillaceae* | 43.73 | 40.60 | 0.00 |
| Green_specialties vs. Turning_color_natural | *Natronobacillus* | *Bacillaceae* | 43.73 | 31.47 | 0.00 |
| Black_natural vs. Green_alkali_treated | *Niallia* | *Bacillaceae* | 2.53 | 4.50 | 0.00 |
| Black_natural vs. Green_specialties | *Niallia* | *Bacillaceae* | 2.53 | 5.92 | 0.00 |
| Green_alkali_treated vs. Green_natural | *Niallia* | *Bacillaceae* | 2.53 | -5.14 | 0.00 |
| Black_natural vs. Green_alkali_treated | *Oceanobacillus* | *Bacillaceae* | 24.76 | 3.27 | 0.01 |
| Black_natural vs. Green_specialties | *Oceanobacillus* | *Bacillaceae* | 24.76 | 29.91 | 0.00 |
| Green_alkali_treated vs. Green_specialties | *Oceanobacillus* | *Bacillaceae* | 24.76 | 26.64 | 0.00 |
| Green_specialties vs. Turning_color_natural | *Oceanobacillus* | *Bacillaceae* | 24.76 | -26.26 | 0.00 |
| Black_natural vs. Green_alkali_treated | *Burkholderia group* | *Burkholderiaceae* | 8.91 | 4.59 | 0.01 |
| Black_natural vs. Green_specialties | *Burkholderia group* | *Burkholderiaceae* | 8.91 | 27.66 | 0.00 |
| Black_natural vs. Turning_color_natural | *Burkholderia group* | *Burkholderiaceae* | 8.91 | 29.28 | 0.00 |
| Green_alkali_treated vs. Green_specialties | *Burkholderia group* | *Burkholderiaceae* | 8.91 | 23.08 | 0.00 |
| Green_alkali_treated vs. Turning_color_natural | *Burkholderia group* | *Burkholderiaceae* | 8.91 | 24.69 | 0.00 |
| Black_natural vs. Green_specialties | *Ralstonia* | *Burkholderiaceae* | 14.63 | 27.93 | 0.00 |
| Black_natural vs. Turning_color_natural | *Ralstonia* | *Burkholderiaceae* | 14.63 | 28.57 | 0.00 |
| Green_alkali_treated vs. Green_specialties | *Ralstonia* | *Burkholderiaceae* | 14.63 | 25.13 | 0.00 |
| Green_alkali_treated vs. Turning_color_natural | *Ralstonia* | *Burkholderiaceae* | 14.63 | 25.77 | 0.00 |
| Black_natural vs. Green_specialties | *Suttonella* | *Cardiobacteriaceae* | 109.17 | 6.66 | 0.00 |
| Green_alkali_treated vs. Green_specialties | *Suttonella* | *Cardiobacteriaceae* | 109.17 | 9.30 | 0.00 |
| Green_specialties vs. Turning_color_natural | *Suttonella* | *Cardiobacteriaceae* | 109.17 | -7.43 | 0.00 |
| Black_natural vs. Green_alkali_treated | *Alkalibacterium* | *Carnobacteriaceae* | 408.71 | -3.12 | 0.00 |
| Green_alkali_treated vs. Green_natural | *Alkalibacterium* | *Carnobacteriaceae* | 408.71 | 4.64 | 0.00 |
| Green_alkali_treated vs. Turning_color_natural | *Alkalibacterium* | *Carnobacteriaceae* | 408.71 | 5.32 | 0.00 |

| contrast | Genus | Family | baseMean | log2FoldChange | padj |
|---|---|---|---|---|---|
| Green_specialties vs. Turning_color_natural | *Alkalibacterium* | *Carnobacteriaceae* | 408.71 | 4.13 | 0.01 |
| Black_natural vs. Green_alkali_treated | *Marinilactibacillus* | *Carnobacteriaceae* | 349.90 | -3.18 | 0.00 |
| Black_natural vs. Green_specialties | *Marinilactibacillus* | *Carnobacteriaceae* | 349.90 | -3.51 | 0.00 |
| Green_alkali_treated vs. Green_natural | *Marinilactibacillus* | *Carnobacteriaceae* | 349.90 | 3.58 | 0.00 |
| Green_alkali_treated vs. Turning_color_natural | *Marinilactibacillus* | *Carnobacteriaceae* | 349.90 | 5.44 | 0.00 |
| Green_specialties vs. Turning_color_natural | *Marinilactibacillus* | *Carnobacteriaceae* | 349.90 | 5.77 | 0.00 |
| Black_natural vs. Green_alkali_treated | *Celerinatantimonas* | *Celerinatantimonadaceae* | 1708.91 | 8.57 | 0.00 |
| Black_natural vs. Green_natural | *Celerinatantimonas* | *Celerinatantimonadaceae* | 1708.91 | 7.66 | 0.00 |
| Green_alkali_treated vs. Turning_color_natural | *Celerinatantimonas* | *Celerinatantimonadaceae* | 1708.91 | -7.37 | 0.00 |
| Black_natural vs. Green_natural | *Acidovorax* | *Comamonadaceae* | 11.49 | 4.78 | 0.00 |
| Black_natural vs. Green_specialties | *Acidovorax* | *Comamonadaceae* | 11.49 | 28.80 | 0.00 |
| Black_natural vs. Turning_color_natural | *Acidovorax* | *Comamonadaceae* | 11.49 | 5.53 | 0.00 |
| Green_alkali_treated vs. Green_specialties | *Acidovorax* | *Comamonadaceae* | 11.49 | 26.94 | 0.00 |
| Green_specialties vs. Turning_color_natural | *Acidovorax* | *Comamonadaceae* | 11.49 | -23.26 | 0.00 |
| Black_natural vs. Green_specialties | *Buttiauxella* | *Enterobacteriaceae* | 5.40 | 7.84 | 0.00 |
| Black_natural vs. Green_specialties | *Enterobacter* | *Enterobacteriaceae* | 344.90 | 4.40 | 0.00 |
| Black_natural vs. Green_natural | *Klebsiella* | *Enterobacteriaceae* | 4.49 | -4.96 | 0.01 |
| Black_natural vs. Green_alkali_treated | *Lelliottia* | *Enterobacteriaceae* | 321.49 | 7.68 | 0.00 |
| Black_natural vs. Green_natural | *Lelliottia* | *Enterobacteriaceae* | 321.49 | 4.53 | 0.00 |
| Black_natural vs. Green_specialties | *Lelliottia* | *Enterobacteriaceae* | 321.49 | 8.20 | 0.00 |
| Black_natural vs. Green_alkali_treated | *Enterococcus* | *Enterococcaceae* | 192.35 | -4.76 | 0.00 |
| Green_alkali_treated vs. Green_natural | *Enterococcus* | *Enterococcaceae* | 192.35 | 3.37 | 0.00 |
| Green_alkali_treated vs. Turning_color_natural | *Enterococcus* | *Enterococcaceae* | 192.35 | 3.57 | 0.01 |
| Black_natural vs. Green_alkali_treated | *Erwinia* | *Erwiniaceae* | 18.18 | 5.20 | 0.00 |

| contrast | Genus | Family | baseMean | log2FoldChange | padj |
|---|---|---|---|---|---|
| Black_natural vs. Green_specialties | *Erwinia* | *Erwiniaceae* | 18.18 | 7.59 | 0.00 |
| Green_alkali_treated vs. Green_natural | *Erwinia* | *Erwiniaceae* | 18.18 | -6.56 | 0.00 |
| Green_alkali_treated vs. Turning_color_natural | *Erwinia* | *Erwiniaceae* | 18.18 | -6.88 | 0.00 |
| Green_specialties vs. Turning_color_natural | *Erwinia* | *Erwiniaceae* | 18.18 | -9.28 | 0.00 |
| Black_natural vs. Green_alkali_treated | *Pantoea* | *Erwiniaceae* | 195.48 | 5.12 | 0.00 |
| Black_natural vs. Green_specialties | *Pantoea* | *Erwiniaceae* | 195.48 | 4.91 | 0.00 |
| Green_alkali_treated vs. Green_natural | *Pantoea* | *Erwiniaceae* | 195.48 | -7.60 | 0.00 |
| Green_alkali_treated vs. Turning_color_natural | *Pantoea* | *Erwiniaceae* | 195.48 | -6.19 | 0.00 |
| Green_specialties vs. Turning_color_natural | *Pantoea* | *Erwiniaceae* | 195.48 | -5.98 | 0.00 |
| Black_natural vs. Green_alkali_treated | *Rosenbergiella* | *Erwiniaceae* | 6.41 | 5.48 | 0.00 |
| Green_alkali_treated vs. Green_natural | *Rosenbergiella* | *Erwiniaceae* | 6.41 | -6.66 | 0.00 |
| Black_natural vs. Green_specialties | *Halanaerobium* | *Halanaerobiaceae* | 11.38 | 27.50 | 0.00 |
| Green_alkali_treated vs. Green_specialties | *Halanaerobium* | *Halanaerobiaceae* | 11.38 | 29.66 | 0.00 |
| Green_alkali_treated vs. Turning_color_natural | *Halanaerobium* | *Halanaerobiaceae* | 11.38 | 8.96 | 0.00 |
| Green_specialties vs. Turning_color_natural | *Halanaerobium* | *Halanaerobiaceae* | 11.38 | -20.70 | 0.00 |
| Green_alkali_treated vs. Green_specialties | *Halomonas* | *Halomonadaceae* | 123.65 | 4.53 | 0.01 |
| Black_natural vs. Green_specialties | *Salinicola* | *Halomonadaceae* | 57.64 | 29.88 | 0.00 |
| Green_alkali_treated vs. Green_specialties | *Salinicola* | *Halomonadaceae* | 57.64 | 30.71 | 0.00 |
| Green_specialties vs. Turning_color_natural | *Salinicola* | *Halomonadaceae* | 57.64 | -30.41 | 0.00 |
| Black_natural vs. Green_alkali_treated | *Terasakiispira* | *Halomonadaceae* | 83.35 | -5.34 | 0.00 |
| Black_natural vs. Green_natural | *Terasakiispira* | *Halomonadaceae* | 83.35 | 4.91 | 0.01 |
| Black_natural vs. Green_specialties | *Terasakiispira* | *Halomonadaceae* | 83.35 | 27.78 | 0.00 |
| Green_alkali_treated vs. Green_natural | *Terasakiispira* | *Halomonadaceae* | 83.35 | 10.24 | 0.00 |
| Green_alkali_treated vs. Green_specialties | *Terasakiispira* | *Halomonadaceae* | 83.35 | 33.12 | 0.00 |

| contrast | Genus | Family | baseMean | log2FoldChange | padj |
|---|---|---|---|---|---|
| Green_alkali_treated vs. Turning_color_natural | *Terasakiispira* | *Halomonadaceae* | 83.35 | 7.92 | 0.00 |
| Green_specialties vs. Turning_color_natural | *Terasakiispira* | *Halomonadaceae* | 83.35 | -25.20 | 0.00 |
| Black_natural vs. Green_specialties | *Companilactobacillus* | *Lactobacillaceae* | 10.32 | 28.54 | 0.00 |
| Green_alkali_treated vs. Green_specialties | *Companilactobacillus* | *Lactobacillaceae* | 10.32 | 27.29 | 0.00 |
| Green_specialties vs. Turning_color_natural | *Companilactobacillus* | *Lactobacillaceae* | 10.32 | -24.77 | 0.00 |
| Black_natural vs. Green_natural | *Lacticaseibacillus* | *Lactobacillaceae* | 141.68 | -2.03 | 0.01 |
| Green_alkali_treated vs. Green_specialties | *Lacticaseibacillus* | *Lactobacillaceae* | 141.68 | 3.93 | 0.00 |
| Green_alkali_treated vs. Turning_color_natural | *Lacticaseibacillus* | *Lactobacillaceae* | 141.68 | 3.70 | 0.00 |
| Black_natural vs. Green_alkali_treated | *Lactiplantibacillus* | *Lactobacillaceae* | 9265.76 | 2.29 | 0.00 |
| Black_natural vs. Green_specialties | *Lactiplantibacillus* | *Lactobacillaceae* | 9265.76 | 2.09 | 0.00 |
| Green_alkali_treated vs. Green_natural | *Lactiplantibacillus* | *Lactobacillaceae* | 9265.76 | -1.78 | 0.00 |
| Green_alkali_treated vs. Turning_color_natural | *Lactiplantibacillus* | *Lactobacillaceae* | 9265.76 | -2.68 | 0.00 |
| Green_specialties vs. Turning_color_natural | *Lactiplantibacillus* | *Lactobacillaceae* | 9265.76 | -2.48 | 0.00 |
| Black_natural vs. Green_natural | *Lactobacillus* | *Lactobacillaceae* | 697.09 | 3.16 | 0.01 |
| Black_natural vs. Green_specialties | *Lactobacillus* | *Lactobacillaceae* | 697.09 | 10.78 | 0.00 |
| Black_natural vs. Turning_color_natural | *Lactobacillus* | *Lactobacillaceae* | 697.09 | 4.54 | 0.00 |
| Green_alkali_treated vs. Green_specialties | *Lactobacillus* | *Lactobacillaceae* | 697.09 | 8.81 | 0.00 |
| Green_specialties vs. Turning_color_natural | *Lactobacillus* | *Lactobacillaceae* | 697.09 | -6.24 | 0.00 |
| Black_natural vs. Green_specialties | *Lentilactobacillus* | *Lactobacillaceae* | 21431.85 | 6.39 | 0.00 |
| Green_alkali_treated vs. Green_specialties | *Lentilactobacillus* | *Lactobacillaceae* | 21431.85 | 5.65 | 0.00 |
| Green_specialties vs. Turning_color_natural | *Lentilactobacillus* | *Lactobacillaceae* | 21431.85 | -6.06 | 0.00 |
| Black_natural vs. Green_alkali_treated | *Leuconostoc* | *Lactobacillaceae* | 798.95 | 11.53 | 0.00 |
| Black_natural vs. Green_natural | *Leuconostoc* | *Lactobacillaceae* | 798.95 | 6.20 | 0.00 |

| contrast | Genus | Family | baseMean | log2FoldChange | padj |
|---|---|---|---|---|---|
| Black_natural vs. Green_specialties | *Leuconostoc* | *Lactobacillaceae* | 798.95 | 6.78 | 0.00 |
| Black_natural vs. Turning_color_natural | *Leuconostoc* | *Lactobacillaceae* | 798.95 | 3.75 | 0.01 |
| Green_alkali_treated vs. Green_natural | *Leuconostoc* | *Lactobacillaceae* | 798.95 | -5.33 | 0.00 |
| Green_alkali_treated vs. Green_specialties | *Leuconostoc* | *Lactobacillaceae* | 798.95 | -4.75 | 0.00 |
| Green_alkali_treated vs. Turning_color_natural | *Leuconostoc* | *Lactobacillaceae* | 798.95 | -7.78 | 0.00 |
| Black_natural vs. Green_alkali_treated | *Levilactobacillus* | *Lactobacillaceae* | 2017.05 | 2.07 | 0.01 |
| Black_natural vs. Green_natural | *Levilactobacillus* | *Lactobacillaceae* | 2017.05 | 2.59 | 0.00 |
| Black_natural vs. Green_specialties | *Levilactobacillus* | *Lactobacillaceae* | 2017.05 | 7.90 | 0.00 |
| Green_alkali_treated vs. Green_specialties | *Levilactobacillus* | *Lactobacillaceae* | 2017.05 | 5.84 | 0.00 |
| Green_specialties vs. Turning_color_natural | *Levilactobacillus* | *Lactobacillaceae* | 2017.05 | -5.40 | 0.00 |
| Black_natural vs. Green_specialties | *Ligilactobacillus* | *Lactobacillaceae* | 58.66 | 4.31 | 0.00 |
| Green_alkali_treated vs. Green_specialties | *Ligilactobacillus* | *Lactobacillaceae* | 58.66 | 4.64 | 0.00 |
| Green_specialties vs. Turning_color_natural | *Ligilactobacillus* | *Lactobacillaceae* | 58.66 | -5.60 | 0.00 |
| Black_natural vs. Green_alkali_treated | *Pediococcus* | *Lactobacillaceae* | 12039.11 | 1.54 | 0.00 |
| Black_natural vs. Green_specialties | *Pediococcus* | *Lactobacillaceae* | 12039.11 | 3.85 | 0.00 |
| Green_alkali_treated vs. Green_specialties | *Pediococcus* | *Lactobacillaceae* | 12039.11 | 2.31 | 0.00 |
| Green_specialties vs. Turning_color_natural | *Pediococcus* | *Lactobacillaceae* | 12039.11 | -3.35 | 0.00 |
| Black_natural vs. Green_alkali_treated | *Secundilactobacillus* | *Lactobacillaceae* | 3969.62 | 2.12 | 0.00 |
| Black_natural vs. Green_specialties | *Secundilactobacillus* | *Lactobacillaceae* | 3969.62 | 3.48 | 0.00 |
| Green_alkali_treated vs. Green_natural | *Secundilactobacillus* | *Lactobacillaceae* | 3969.62 | -3.17 | 0.00 |
| Green_specialties vs. Turning_color_natural | *Secundilactobacillus* | *Lactobacillaceae* | 3969.62 | -2.74 | 0.00 |
| Black_natural vs. Green_alkali_treated | *Weissella* | *Lactobacillaceae* | 581.65 | 8.15 | 0.00 |
| Black_natural vs. Green_natural | *Weissella* | *Lactobacillaceae* | 581.65 | 6.74 | 0.00 |
| Black_natural vs. Green_specialties | *Weissella* | *Lactobacillaceae* | 581.65 | 6.10 | 0.00 |

| contrast | Genus | Family | baseMean | log2FoldChange | padj |
|---|---|---|---|---|---|
| Green_alkali_treated vs. Turning_color_natural | *Weissella* | *Lactobacillaceae* | 581.65 | -5.14 | 0.00 |
| Black_natural vs. Green_alkali_treated | *Stenotrophomonas* | *Lysobacteraceae* | 3.50 | 3.97 | 0.00 |
| Black_natural vs. Green_specialties | *Stenotrophomonas* | *Lysobacteraceae* | 3.50 | 6.66 | 0.00 |
| Black_natural vs. Green_natural | *Marinobacter* | *Marinobacteraceae* | 3.87 | 3.93 | 0.01 |
| Black_natural vs. Green_specialties | *Marinobacter* | *Marinobacteraceae* | 3.87 | 4.78 | 0.01 |
| Black_natural vs. Green_alkali_treated | *Acinetobacter* | *Moraxellaceae* | 157.05 | 5.59 | 0.00 |
| Black_natural vs. Green_natural | *Acinetobacter* | *Moraxellaceae* | 157.05 | 2.80 | 0.01 |
| Black_natural vs. Green_specialties | *Acinetobacter* | *Moraxellaceae* | 157.05 | 7.12 | 0.00 |
| Black_natural vs. Turning_color_natural | *Acinetobacter* | *Moraxellaceae* | 157.05 | 4.62 | 0.00 |
| Black_natural vs. Green_alkali_treated | *Janthinobacterium* | *Oxalobacteraceae* | 5.12 | 5.39 | 0.00 |
| Black_natural vs. Green_natural | *Janthinobacterium* | *Oxalobacteraceae* | 5.12 | 5.78 | 0.01 |
| Black_natural vs. Green_specialties | *Janthinobacterium* | *Oxalobacteraceae* | 5.12 | 7.08 | 0.01 |
| Black_natural vs. Green_alkali_treated | *Pseudomonas* | *Pseudomonadaceae* | 101.46 | 4.04 | 0.00 |
| Black_natural vs. Green_specialties | *Pseudomonas* | *Pseudomonadaceae* | 101.46 | 5.72 | 0.00 |
| Green_alkali_treated vs. Green_natural | *Pseudomonas* | *Pseudomonadaceae* | 101.46 | -3.53 | 0.00 |
| Black_natural vs. Green_alkali_treated | *Shewanella* | *Shewanellaceae* | 16.74 | -3.86 | 0.00 |
| Black_natural vs. Green_specialties | *Shewanella* | *Shewanellaceae* | 16.74 | 6.38 | 0.00 |
| Green_alkali_treated vs. Green_specialties | *Shewanella* | *Shewanellaceae* | 16.74 | 10.24 | 0.00 |
| Green_specialties vs. Turning_color_natural | *Shewanella* | *Shewanellaceae* | 16.74 | -9.24 | 0.00 |
| Black_natural vs. Green_alkali_treated | *Sporolactobacillus* | *Sporolactobacillaceae* | 27.43 | -6.07 | 0.00 |
| Green_alkali_treated vs. Green_natural | *Sporolactobacillus* | *Sporolactobacillaceae* | 27.43 | 7.13 | 0.00 |
| Black_natural vs. Green_alkali_treated | *Staphylococcus* | *Staphylococcaceae* | 6.74 | 3.70 | 0.01 |
| Black_natural vs. Green_specialties | *Staphylococcus* | *Staphylococcaceae* | 6.74 | 6.98 | 0.00 |
| Green_alkali_treated vs. Green_natural | *Staphylococcus* | *Staphylococcaceae* | 6.74 | -4.89 | 0.01 |
| Green_specialties vs. Turning_color_natural | *Staphylococcus* | *Staphylococcaceae* | 6.74 | -6.90 | 0.00 |

| contrast | Genus | Family | baseMean | log2FoldChange | padj |
|---|---|---|---|---|---|
| Black_natural vs. Green_specialties | *Catenococcus* | *Vibrionaceae* | 3.76 | 22.78 | 0.00 |
| Black_natural vs. Turning_color_natural | *Catenococcus* | *Vibrionaceae* | 3.76 | 20.13 | 0.00 |
| Green_alkali_treated vs. Green_specialties | *Catenococcus* | *Vibrionaceae* | 3.76 | 27.50 | 0.00 |
| Green_alkali_treated vs. Turning_color_natural | *Catenococcus* | *Vibrionaceae* | 3.76 | 24.84 | 0.00 |
| Green_alkali_treated vs. Green_natural | *Vibrio* | *Vibrionaceae* | 51.99 | 5.42 | 0.01 |
| Black_natural vs. Green_alkali_treated | *Rahnella* | *Yersiniaceae* | 19.65 | 3.82 | 0.00 |
| Green_alkali_treated vs. Green_natural | *Rahnella* | *Yersiniaceae* | 19.65 | -5.45 | 0.00 |

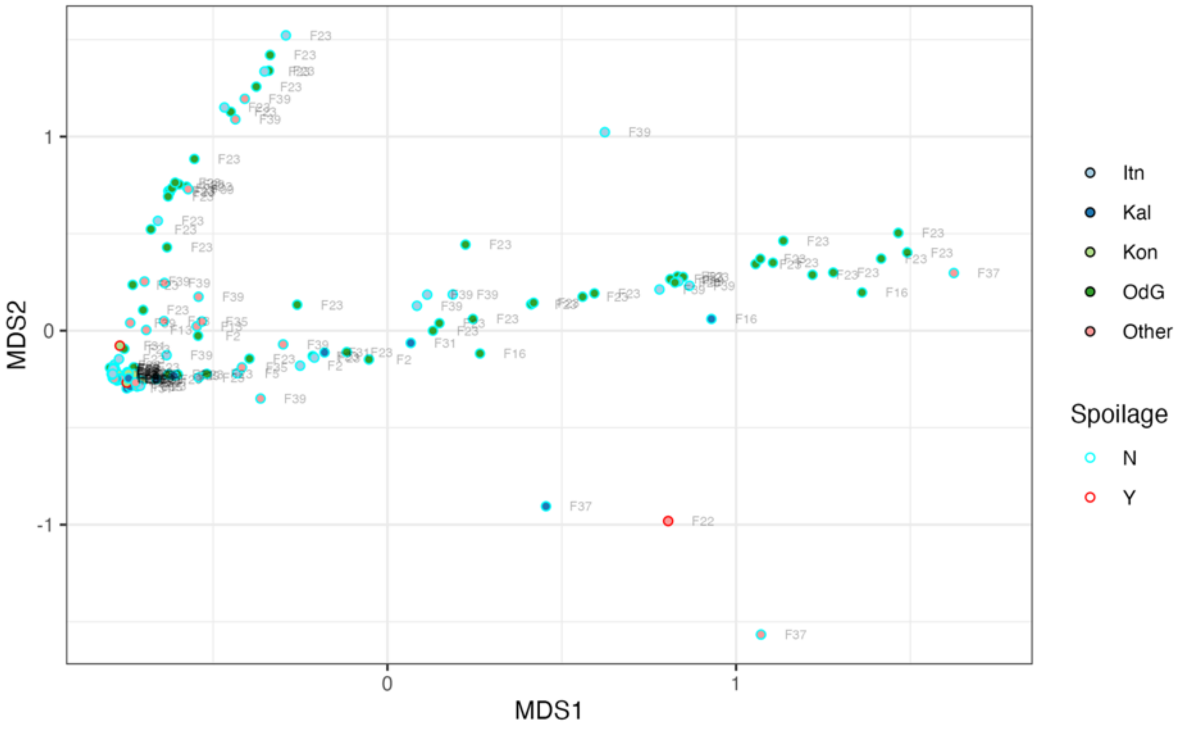


**Supplementary Figure 48.** Configuration plot for the first two dimensions of a non-monotonic Multidimensional Scaling of the Bray-Curtis distance matrix of the composition of fungal communities of Black natural table olives.

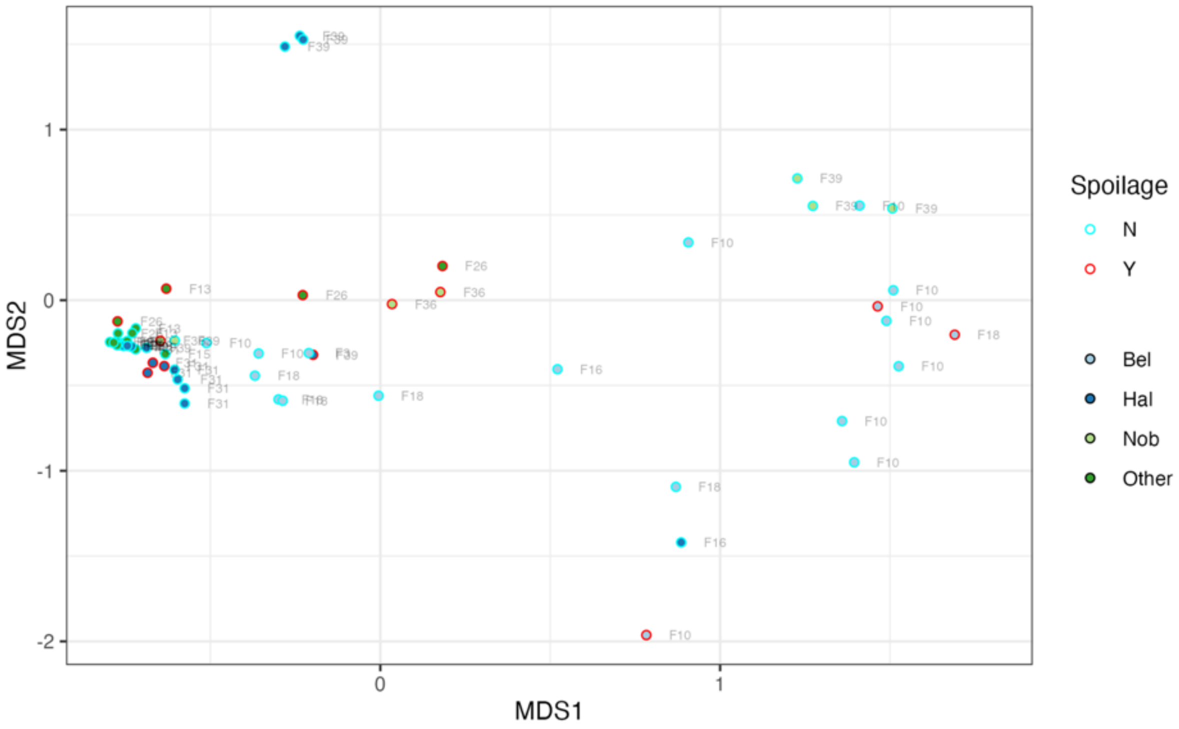


**Supplementary Figure 49.** Configuration plot for the first two dimensions of a non-monotonic Multidimensional Scaling of the Bray-Curtis distance matrix of the composition of fungal communities of Green alkali treated table olives.

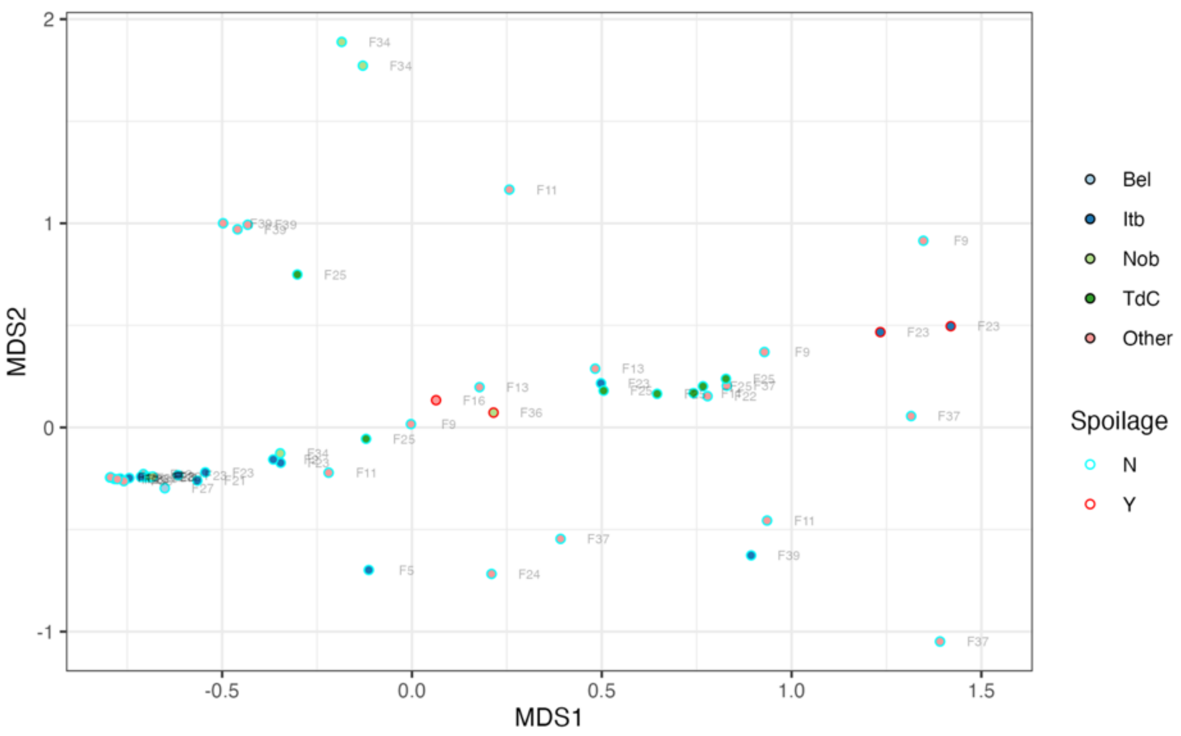


**Supplementary Figure 50.** Configuration plot for the first two dimensions of a non-monotonic Multidimensional Scaling of the Bray-Curtis distance matrix of the composition of fungal communities of Green natural table olives.

**Supplementary table 25**. Results of PERMANOVA to test the hypothesis that variety did not affect the composition of fungal communities of table olives within a given ripeness and trade preparation group.

| Olive ripeness + trade prep. | samples | varieties | F | $R^2$ | Prob |
|---|---|---|---|---|---|
| Black natural | 124 | 5 | 4.500 | 0.130 | 0.001 |
| Green alkali treated | 53 | 4 | 5.888 | 0.265 | 0.001 |
| Green natural | 31 | 3 | 3.649 | 0.207 | 0.005 |
| Turning color natural | 21 | 3 | 2.563 | 0.222 | 0.030 |

**Supplementary Table 26**. Results of PERMANOVA to test the hypothesis that firm did not affect the composition of fungal communities of table olives within selected varieties.

| Olive ripeness + trade prep. | Olive variety | samples | Firms | F | $R^2$ | Prob |
|---|---|---|---|---|---|---|
| Black natural | Itrana nera | 33 | 2 | 12.243 | 0.283 | 0.002 |
| Black natural | Kalamata* | 11 | 2 | 4.181 | 0.317 | 0.017 |
| Green alkali treated | Bella di Daunia | 17 | 2 | 1.909 | 0.080 | 0.115 |
| Green alkali treated | Halkidiki | 20 | 3 | 6.038 | 0.415 | 0.002 |

* results of the betadispers() test were significant.

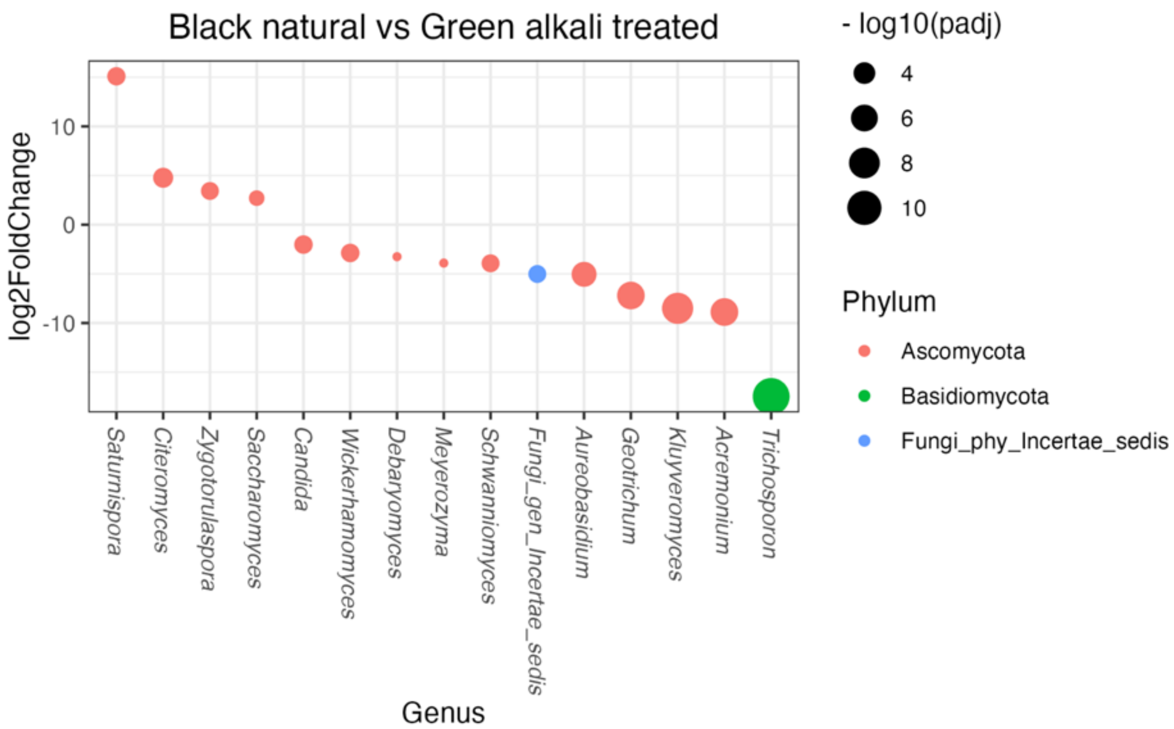


**Supplementary Figure 50**. Differentially abundant fungal genera for the contrast Black natural – Green alkali treated, DeSeq2.

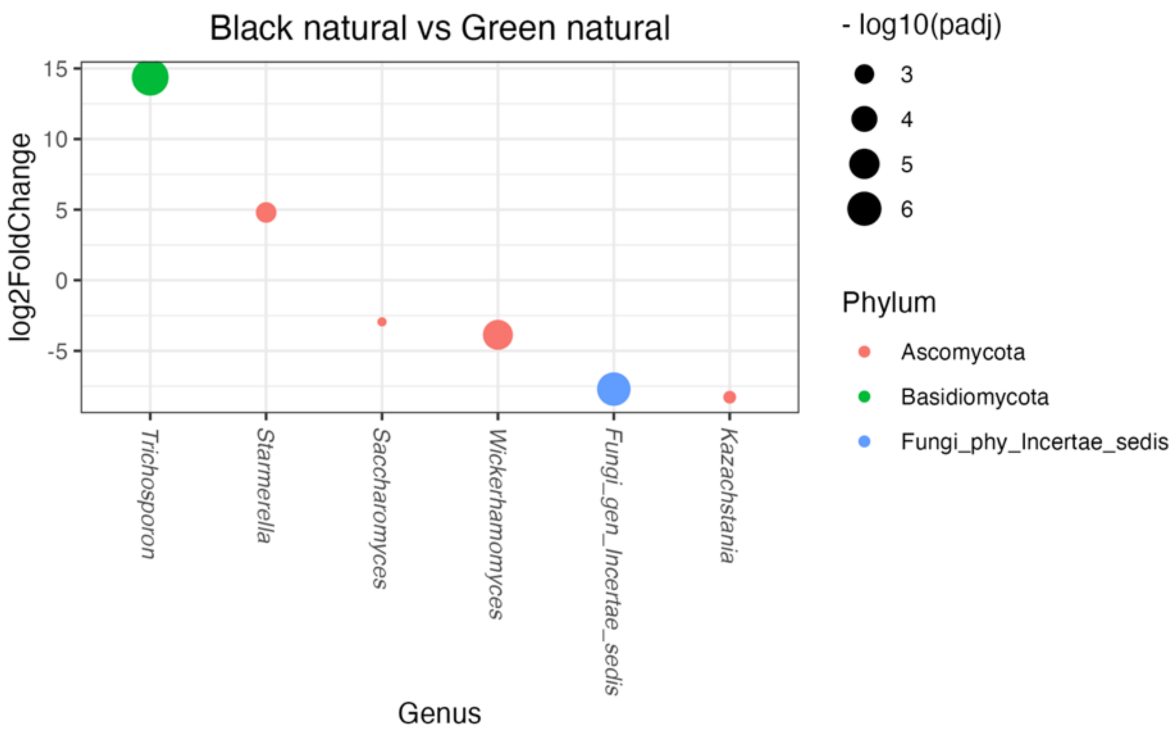


**Supplementary Figure 51.** Differentially abundant fungal genera for the contrast Black natural – Green natural, DeSeq2.

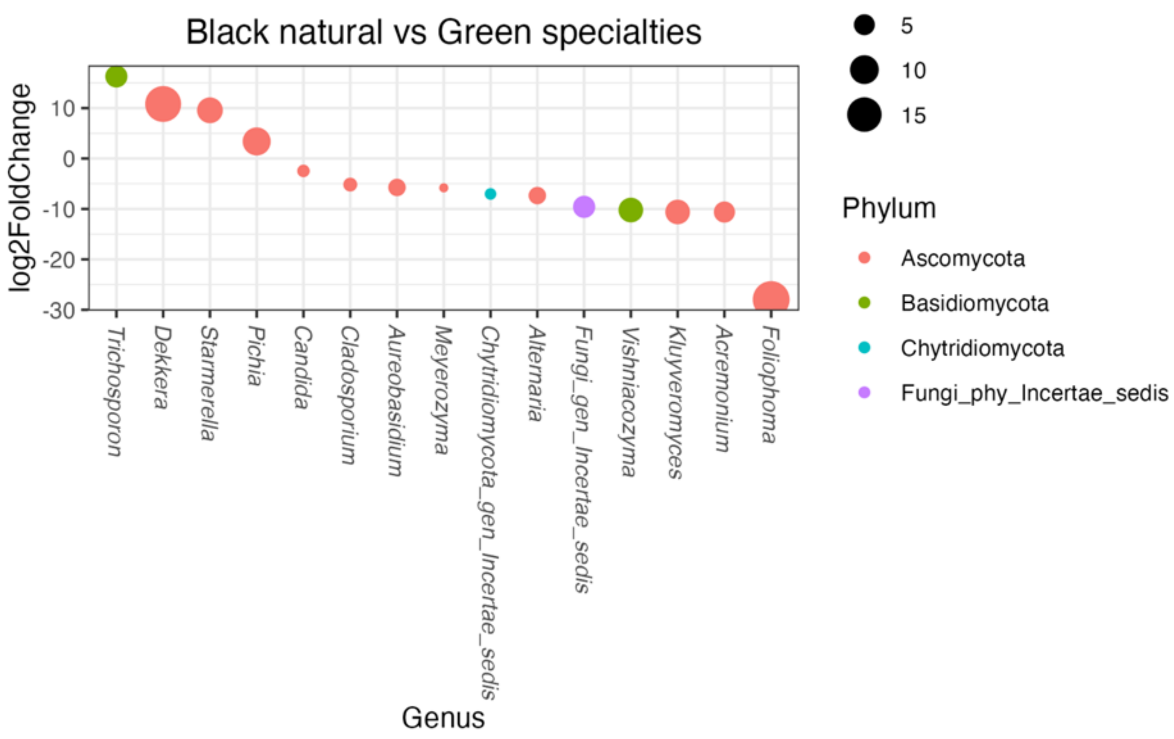


**Supplementary Figure 52**. Differentially abundant fungal genera for the contrast Black natural – Green specialties, DeSeq2.

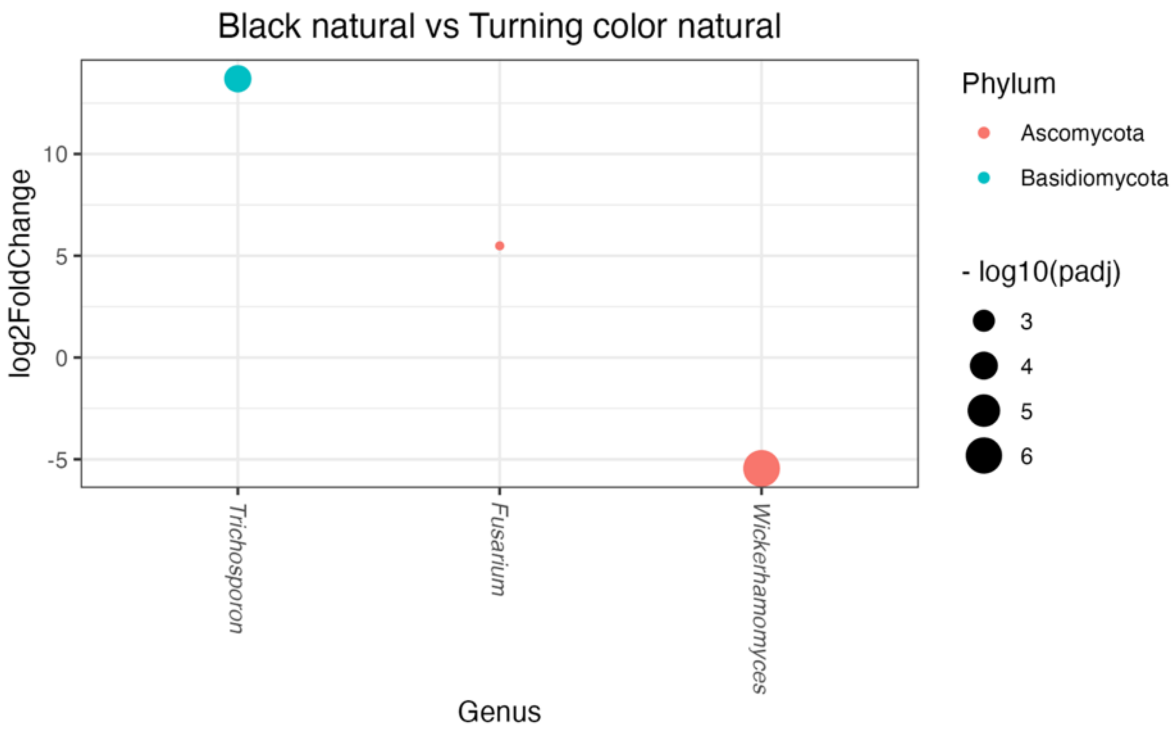


**Supplementary Figure 53**. Differentially abundant fungal genera for the contrast Black natural – Turning color natural, DeSeq2.

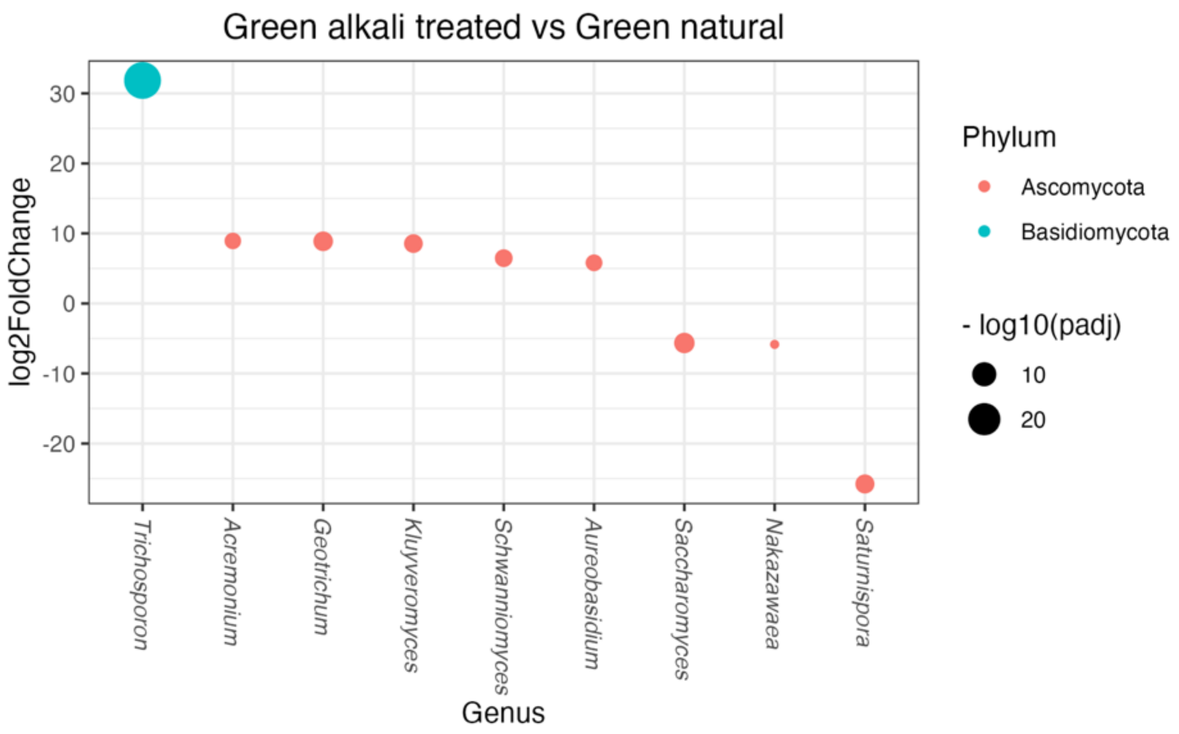


**Supplementary Figure 54**. Differentially abundant fungal genera for the contrast Green alkali treated – Green natural, DeSeq2.

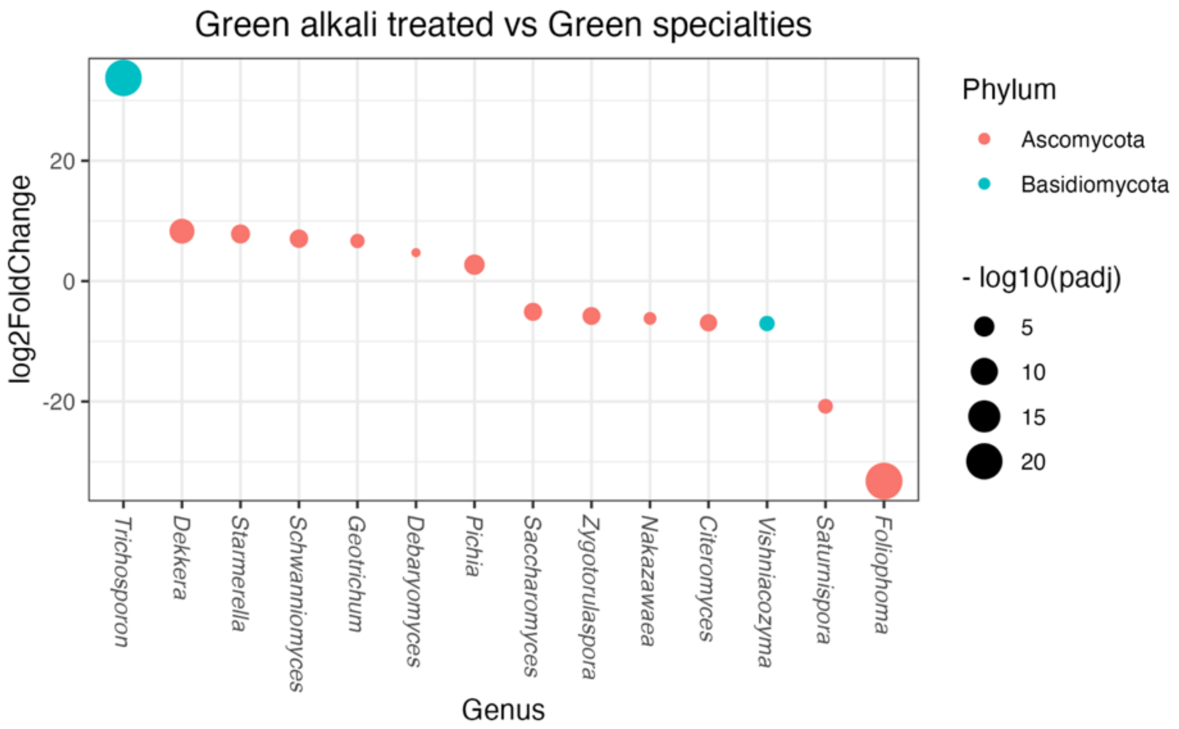


**Supplementary Figure 55**. Differentially abundant fungal genera for the contrast Green alkali treated – Green specialties, DeSeq2.

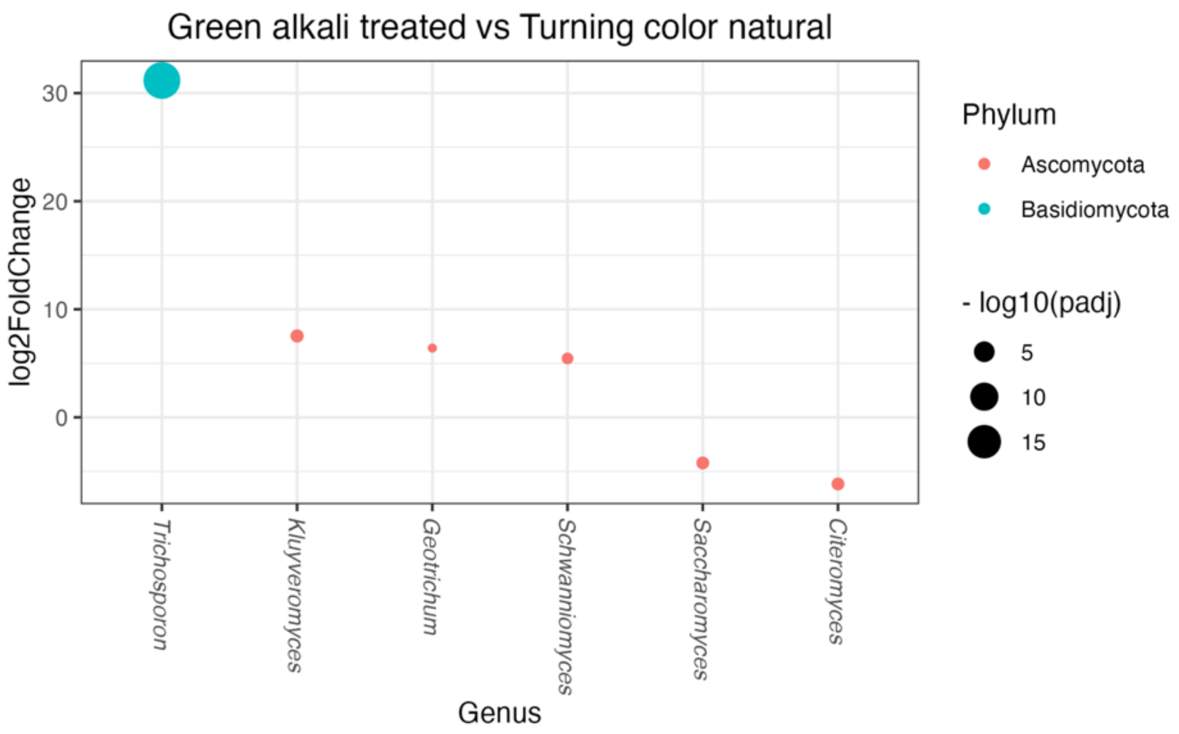


**Supplementary Figure 56.** Differentially abundant fungal genera for the contrast Green alkali treated – Turning color natural, DeSeq2.

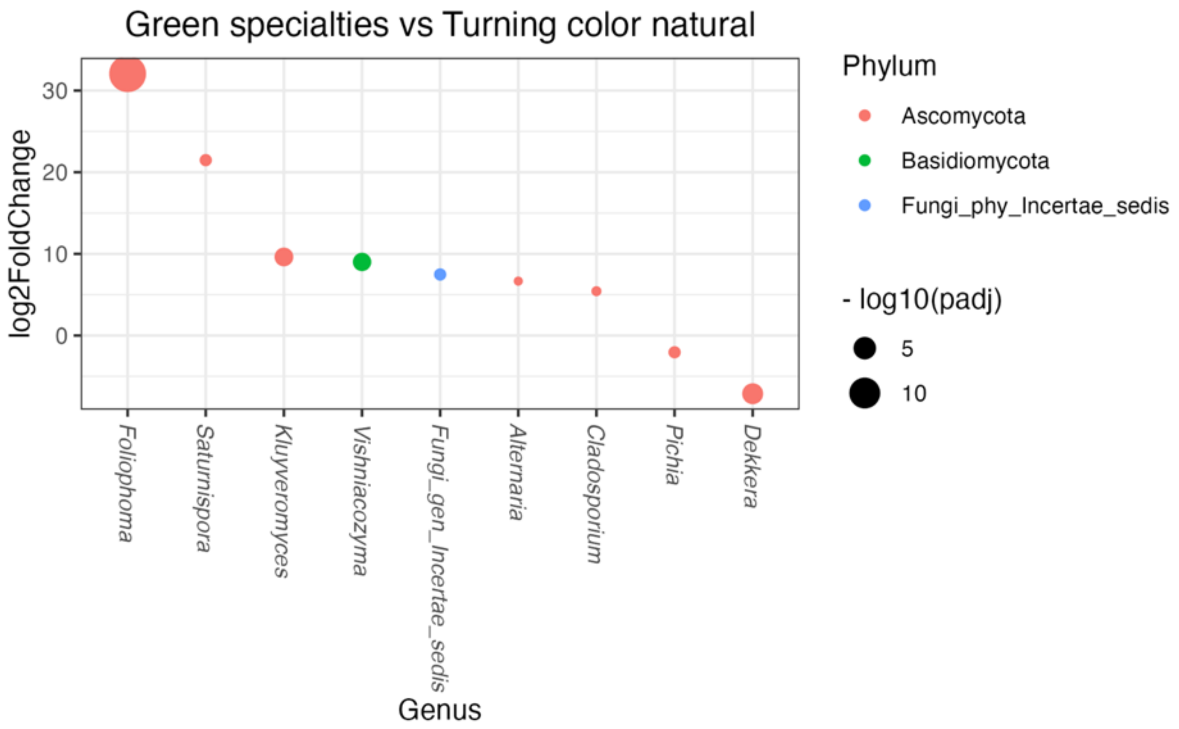


**Supplementary Figure 57.** Differentially abundant fungal genera for the contrast Green specialties – Turning color natural, DeSeq2.

**Supplementary table 26.** Summary results for DeSeq2 differential abundance analysis for fungal genera.

| contrast | Genus | Family | Base Mean | log2FoldChange | lfcSE | stat | p value | padj |
|---|---|---|---|---|---|---|---|---|
| Black_natural vs. Green_alkali_treated | Acremonium | Bionectriaceae | 5.83 | -8.87 | 1.59 | -5.58 | 0 | 0.00 |
| Black_natural vs. Green_specialties | Acremonium | Bionectriaceae | 5.83 | -10.35 | 2.17 | -4.77 | 0 | 0.00 |
| Green_alkali_treated vs. Green_natural | Acremonium | Bionectriaceae | 5.83 | 8.96 | 1.93 | 4.63 | 0 | 0.00 |
| Black_natural vs. Green_specialties | Vishniacozyma | Bulleribasidiaceae | 30.80 | -9.85 | 1.76 | -5.59 | 0 | 0.00 |
| Green_alkali_treated vs. Green_specialtiesl | Vishniacozyma | Bulleribasidiaceae | 30.80 | -6.66 | 1.94 | -3.44 | 0 | 0.00 |
| Green_specialties vs. Turning_color_natural | Vishniacozyma | Bulleribasidiaceae | 30.80 | 8.72 | 2.20 | 3.95 | 0 | 0.00 |
| Black_natural vs. Green_specialties | Chytridiomycota_gen_Incertae_sedis | Chytridiomycota_fam_Incertae_sedis | 5.91 | -6.67 | 2.09 | -3.19 | 0 | 0.01 |
| Black_natural vs. Green_specialties | Cladosporium | Cladosporiaceae | 27.84 | -4.81 | 1.42 | -3.38 | 0 | 0.00 |
| Black_natural vs. Green_specialties | Foliophoma | Coniothyriaceae | 4.08 | -28.11 | 3.12 | -9.01 | 0 | 0.00 |
| Green_alkali_treated vs. Green_specialtiesl | Foliophoma | Coniothyriaceae | 4.08 | -36.26 | 3.44 | -10.55 | 0 | 0.00 |
| Green_specialties vs. Turning_color_natural | Foliophoma | Coniothyriaceae | 4.08 | 32.65 | 3.89 | 8.39 | 0 | 0.00 |
| Black_natural vs. Green_alkali_treated | Debaryomyces | Debaryomycetaceae | 111.80 | -3.17 | 1.07 | -2.97 | 0 | 0.01 |
| Green_alkali_treated vs. Green_specialtiesl | Debaryomyces | Debaryomycetaceae | 111.80 | 4.99 | 1.61 | 3.09 | 0 | 0.01 |
| Black_natural vs. Green_alkali_treated | Meyerozyma | Debaryomycetaceae | 6.32 | -3.86 | 1.27 | -3.04 | 0 | 0.01 |
| Black_natural vs. Green_specialties | Meyerozyma | Debaryomycetaceae | 6.32 | -5.74 | 1.73 | -3.31 | 0 | 0.00 |
| Black_natural vs. Green_alkali_treated | Schwanniomyces | Debaryomycetaceae | 40.39 | -3.87 | 1.05 | -3.68 | 0 | 0.00 |
| Green_alkali_treated vs. Green_natural | Schwanniomyces | Debaryomycetaceae | 40.39 | 6.72 | 1.29 | 5.21 | 0 | 0.00 |

| contrast | Genus | Family | Base Mean | log2FoldChange | lfcSE | stat | p value | padj |
|---|---|---|---|---|---|---|---|---|
| Green_alkali_treated vs. Green_specialtiesl | Schwanniomyces | Debaryomycetaceae | 40.39 | 6.98 | 1.60 | 4.37 | 0 | 0.00 |
| Green_alkali_treated vs. Turning_color_natural | Schwanniomyces | Debaryomycetaceae | 40.39 | 5.43 | 1.51 | 3.60 | 0 | 0.00 |
| Black_natural vs. Green_alkali_treated | Geotrichum | Dipodascaceae | 89.66 | -8.00 | 1.24 | -6.44 | 0 | 0.00 |
| Green_alkali_treated vs. Green_natural | Geotrichum | Dipodascaceae | 89.66 | 8.89 | 1.52 | 5.86 | 0 | 0.00 |
| Green_alkali_treated vs. Green_specialtiesl | Geotrichum | Dipodascaceae | 89.66 | 6.70 | 1.87 | 3.58 | 0 | 0.00 |
| Green_alkali_treated vs. Turning_color_natural | Geotrichum | Dipodascaceae | 89.66 | 6.02 | 1.77 | 3.40 | 0 | 0.01 |
| Black_natural vs. Green_alkali_treated | Fungi_gen_Incertae_sedis | Fungi_fam_Incertae_sedis | 35.24 | -5.02 | 1.35 | -3.70 | 0 | 0.00 |
| Black_natural vs. Green_natural | Fungi_gen_Incertae_sedis | Fungi_fam_Incertae_sedis | 35.24 | -7.71 | 1.40 | -5.50 | 0 | 0.00 |
| Black_natural vs. Green_specialties | Fungi_gen_Incertae_sedis | Fungi_fam_Incertae_sedis | 35.24 | -9.42 | 1.85 | -5.10 | 0 | 0.00 |
| Black_natural vs. Green_alkali_treated | Kodamaea | Metschnikowiaceae | 8.61 | -7.98 | 2.35 | -3.40 | 0 | 0.00 |
| Black_natural vs. Turning_color_natural | Fusarium | Nectriaceae | 7.47 | 5.48 | 1.51 | 3.63 | 0 | 0.01 |
| Black_natural vs. Green_alkali_treated | Wickerhamomyces | Phaffomycetaceae | 5290.43 | -2.88 | 0.74 | -3.89 | 0 | 0.00 |
| Black_natural vs. Green_natural | Wickerhamomyces | Phaffomycetaceae | 5290.43 | -3.80 | 0.77 | -4.94 | 0 | 0.00 |
| Black_natural vs. Turning_color_natural | Wickerhamomyces | Phaffomycetaceae | 5290.43 | -5.49 | 0.95 | -5.79 | 0 | 0.00 |
| Black_natural vs. Green_specialties | Dekkera | Pichiaceae | 17095.94 | 10.84 | 1.23 | 8.83 | 0 | 0.00 |
| Green_alkali_treated vs. Green_specialtiesl | Dekkera | Pichiaceae | 17095.94 | 8.32 | 1.35 | 6.15 | 0 | 0.00 |
| Green_specialties vs. Turning_color_natural | Dekkera | Pichiaceae | 17095.94 | -8.08 | 1.53 | -5.28 | 0 | 0.00 |
| Black_natural vs. Green_specialties | Pichia | Pichiaceae | 52964.03 | 3.40 | 0.51 | 6.71 | 0 | 0.00 |
| Green_alkali_treated vs. Green_specialtiesl | Pichia | Pichiaceae | 52964.03 | 2.71 | 0.56 | 4.85 | 0 | 0.00 |

| contrast | Genus | Family | Base Mean | log2FoldChange | lfcSE | stat | p value | padj |
|---|---|---|---|---|---|---|---|---|
| Black_natural vs. Green_alkali_treated | Saturnispora | Pichiaceae | 3.42 | 15.17 | 3.94 | 3.85 | 0 | 0.00 |
| Green_alkali_treated vs. Green_natural | Saturnispora | Pichiaceae | 3.42 | -25.80 | 4.78 | -5.40 | 0 | 0.00 |
| Green_alkali_treated vs. Green_specialtiesl | Saturnispora | Pichiaceae | 3.42 | -20.84 | 5.94 | -3.51 | 0 | 0.00 |
| Black_natural vs. Green_alkali_treated | Gibellulopsis | Plectosphaerellaceae | 1.67 | -21.68 | 2.13 | -10.17 | 0 | 0.00 |
| Black_natural vs. Green_specialties | Gibellulopsis | Plectosphaerellaceae | 1.67 | -26.21 | 2.90 | -9.05 | 0 | 0.00 |
| Green_alkali_treated vs. Green_natural | Gibellulopsis | Plectosphaerellaceae | 1.67 | 32.77 | 2.59 | 12.67 | 0 | 0.00 |
| Green_alkali_treated vs. Turning_color_natural | Gibellulopsis | Plectosphaerellaceae | 1.67 | 28.55 | 3.04 | 9.39 | 0 | 0.00 |
| Green_specialties vs. Turning_color_natural | Gibellulopsis | Plectosphaerellaceae | 1.67 | 33.08 | 3.62 | 9.14 | 0 | 0.00 |
| Black_natural vs. Green_specialties | Alternaria | Pleosporaceae | 4.68 | -7.15 | 1.76 | -4.06 | 0 | 0.00 |
| Black_natural vs. Green_alkali_treated | Citeromyces | Saccharomycetaceae | 33.01 | 4.77 | 1.15 | 4.16 | 0 | 0.00 |
| Green_alkali_treated vs. Green_specialtiesl | Citeromyces | Saccharomycetaceae | 33.01 | -6.75 | 1.71 | -3.94 | 0 | 0.00 |
| Green_alkali_treated vs. Turning_color_natural | Citeromyces | Saccharomycetaceae | 33.01 | -6.17 | 1.62 | -3.81 | 0 | 0.00 |
| Black_natural vs. Green_natural | Kazachstania | Saccharomycetaceae | 13.21 | -8.01 | 2.32 | -3.46 | 0 | 0.01 |
| Black_natural vs. Green_alkali_treated | Kluyveromyces | Saccharomycetaceae | 1029.03 | -8.48 | 1.34 | -6.32 | 0 | 0.00 |
| Black_natural vs. Green_specialties | Kluyveromyces | Saccharomycetaceae | 1029.03 | -10.56 | 1.83 | -5.76 | 0 | 0.00 |
| Green_alkali_treated vs. Green_natural | Kluyveromyces | Saccharomycetaceae | 1029.03 | 8.54 | 1.63 | 5.24 | 0 | 0.00 |
| Green_alkali_treated vs. Turning_color_natural | Kluyveromyces | Saccharomycetaceae | 1029.03 | 7.52 | 1.91 | 3.93 | 0 | 0.00 |
| Green_specialties vs. Turning_color_natural | Kluyveromyces | Saccharomycetaceae | 1029.03 | 9.60 | 2.29 | 4.20 | 0 | 0.00 |
| Black_natural vs. Green_alkali_treated | Saccharomyces | Saccharomycetaceae | 3813.65 | 2.61 | 0.79 | 3.31 | 0 | 0.00 |

| contrast | Genus | Family | Base Mean | log2FoldChange | lfcSE | stat | p value | padj |
|---|---|---|---|---|---|---|---|---|
| Black_natural vs. Green_natural | Saccharomyces | Saccharomycetaceae | 3813.65 | -2.86 | 0.82 | -3.49 | 0 | 0.01 |
| Green_alkali_treated vs. Green_natural | Saccharomyces | Saccharomycetaceae | 3813.65 | -5.47 | 0.96 | -5.71 | 0 | 0.00 |
| Green_alkali_treated vs. Green_specialtiesl | Saccharomyces | Saccharomycetaceae | 3813.65 | -4.98 | 1.19 | -4.19 | 0 | 0.00 |
| Green_alkali_treated vs. Turning_color_natural | Saccharomyces | Saccharomycetaceae | 3813.65 | -4.22 | 1.13 | -3.75 | 0 | 0.00 |
| Black_natural vs. Green_alkali_treated | Zygotorulaspora | Saccharomycetaceae | 519.43 | 3.34 | 0.91 | 3.67 | 0 | 0.00 |
| Green_alkali_treated vs. Green_specialtiesl | Zygotorulaspora | Saccharomycetaceae | 519.43 | -5.78 | 1.37 | -4.21 | 0 | 0.00 |
| Green_alkali_treated vs. Turning_color_natural | Zygotorulaspora | Saccharomycetaceae | 519.43 | -4.34 | 1.30 | -3.34 | 0 | 0.01 |
| Black_natural vs. Green_alkali_treated | Candida | Saccharomycetales_fam_Incertae_sedis | 8332.70 | -2.07 | 0.52 | -3.99 | 0 | 0.00 |
| Black_natural vs. Green_specialties | Candida | Saccharomycetales_fam_Incertae_sedis | 8332.70 | -2.51 | 0.71 | -3.54 | 0 | 0.00 |
| Green_alkali_treated vs. Green_specialtiesl | Nakazawaea | Saccharomycetales_fam_Incertae_sedis | 12.71 | -6.03 | 1.94 | -3.10 | 0 | 0.01 |
| Black_natural vs. Green_specialties | Aureobasidium | Saccotheciaceae | 27.57 | -5.61 | 1.29 | -4.34 | 0 | 0.00 |
| Green_alkali_treated vs. Green_specialtiesl | Aureobasidium | Saccotheciaceae | 27.57 | -4.88 | 1.43 | -3.42 | 0 | 0.00 |
| Black_natural vs. Green_natural | Starmerella | Trichomonascaceae | 226.48 | 4.91 | 1.17 | 4.21 | 0 | 0.00 |
| Black_natural vs. Green_specialties | Starmerella | Trichomonascaceae | 226.48 | 9.69 | 1.57 | 6.19 | 0 | 0.00 |
| Green_alkali_treated vs. Green_specialtiesl | Starmerella | Trichomonascaceae | 226.48 | 7.85 | 1.72 | 4.57 | 0 | 0.00 |
| Black_natural vs. Green_alkali_treated | Trichosporon | Trichosporonaceae | 220.75 | -17.32 | 2.32 | -7.47 | 0 | 0.00 |
| Black_natural vs. Green_natural | Trichosporon | Trichosporonaceae | 220.75 | 12.94 | 2.42 | 5.35 | 0 | 0.00 |
| Black_natural vs. Green_specialties | Trichosporon | Trichosporonaceae | 220.75 | 17.89 | 3.19 | 5.60 | 0 | 0.00 |
| Black_natural vs. Turning_color_natural | Trichosporon | Trichosporonaceae | 220.75 | 14.80 | 2.98 | 4.96 | 0 | 0.00 |

| contrast | Genus | Family | Base Mean | log2FoldChange | lfcSE | stat | p value | padj |
|---|---|---|---|---|---|---|---|---|
| Green_alkali_treated vs. Green_natural | Trichosporon | Trichosporonaceae | 220.75 | 30.26 | 2.82 | 10.74 | 0 | 0.00 |
| Green_alkali_treated vs. Green_specialtiesl | Trichosporon | Trichosporonaceae | 220.75 | 35.21 | 3.50 | 10.05 | 0 | 0.00 |
| Green_alkali_treated vs. Turning_color_natural | Trichosporon | Trichosporonaceae | 220.75 | 32.12 | 3.32 | 9.69 | 0 | 0.00 |

**Supplementary Table 27.** List of accession numbers for the raw sequences deposited in NCBI SRA and used in this study. The list includes blanks and mock communities.

SRR34676602-SRR34676664
SRR34676666-SRR34676685
SRR34676687-SRR34676735
SRR34676737-SRR34676743
SRR34676745-SRR34676754
SRR34676756-SRR34676784
SRR34676787-SRR34676791
SRR34676793-SRR34676849
SRR34676851-SRR34676852
SRR34676855-SRR34676860
SRR34676863-SRR34676864
SRR34676866-SRR34676883
SRR34676885-SRR34676887
SRR34676890-SRR34676896
SRR34676899-SRR34677072
SRR34677074-SRR34677113
SRR34677116-SRR34677121
SRR34677124-SRR34677132
SRR34677135-SRR34677148
SRR34677150
SRR34677152-SRR34677159
SRR34677161-SRR34677162
SRR34677164-SRR34677166
SRR34677169-SRR34677172
SRR34677174-SRR34677175
SRR34677177
SRR34677179-SRR34677182
SRR34677185-SRR34677186
SRR34677190-SRR34677288
SRR34677293
SRR34677298-SRR34677317
SRR34677319-SRR34677335
